# Highjacked by a pseudoenzyme: How eudicot plants make indole

**DOI:** 10.1101/2024.08.26.609694

**Authors:** Matilde Florean, Hedwig Schultz, Veit Grabe, Katrin Luck, Sarah E. O’Connor, Tobias G. Köllner

**Affiliations:** Department of Natural Product Biosynthesis, Max Planck Institute for Chemical Ecology, Jena, Germany; Microscopic Imaging Service Group, Max Planck Institute for Chemical Ecology, Jena, Germany

## Abstract

Indole is crucial for plant defense, where it is released as a signaling volatile upon herbivore attack and also serves as a starting precursor for defensive specialized metabolites. Indole is known to be synthesized in plants from indole-3-glycerol phosphate by the enzyme indole-3-glycerol phosphate lyase. Here we report that in core eudicots, indole production for plant defense occurs via an alternative pathway. The α subunit of tryptophan synthase (TSA), an enzyme of core metabolism, normally binds to tryptophan synthase β subunit (TSB) to produce tryptophan. However, we show that a non-catalytic TSB paralogue (TSB-like) can highjack TSA to produce indole. The widespread occurrence of *TSB-like* genes in eudicots suggests that this alternative mechanism for indole formation is widespread throughout the plant kingdom.

## Main Text

Indole is a nitrogen-containing aromatic compound that acts as central intermediate in the biosynthesis of the amino acid tryptophan in all forms of life. In plants, indole also serves as a precursor for specialized defense metabolites such as benzoxazinoids (BXD) (*1*), nudicaulins (*2*), and indigoids (*3*). Moreover, many plants release volatile indole upon herbivory attack to either deter the herbivore or to warn neighboring plants of impending attack, thereby priming plant resistance (*4*–*7*). Indole is also released as a flower volatile that is involved in attracting pollinators (*8, 9*). Indole that is utilized for tryptophan formation is produced by the tryptophan synthase α subunit (TSA) from indole-3-glycerol phosphate (IGP) (*10, 11*). TSA is always found as a highly conserved heterotetrameric complex with the tryptophan synthase β-subunit (TSB), which allows indole to be channeled from the active site of TSA into the active site of TSB, where it is then condensed with L-Ser to form tryptophan (*12*). TSA and TSB alone have very low catalytic activity; the heteromeric dimerization provides the mutual allosteric activation that is required for both of these enzymes to work efficiently (*10*–*12*). In contrast to tryptophan biosynthesis, indole for defense or signaling is produced by two types of plant IGP-lyases that have evolved independently from TSA. One type, commonly referred to as IGL (indole-3-glycerol phosphate lyase), is primarily involved in production of volatile indole, while the second type, referred to as BX1 (benzoxazinoneless-1), produces indole for BXD formation, and appears to be restricted to plants that produce BXD natural products (*13*–*16*). Unlike TSA, IGL and BX1 are highly active as standalone proteins and do not require allosteric activation (*17*–*19*). Functional IGL and BX1 enzymes have so far only been reported in the grasses (Poaceae), a family of monocotyledons, and in one species of the Ranunculaceae (*Consolida orientalis*), a family of basal eudicotyledons (Fig. 1A and fig. S1A) (*2, 16, 20, 21*). Recently, the genes responsible for BXD biosynthesis in two core eudicot species, *Lamium galeobdolon* and *Aphelandra squarrosa*, have been discovered (*16, 22*). However, the respective putative IGL/BX1 enzymes produced indole with poor catalytic efficiencies (*16*). Therefore, it was unclear how free indole is produced in these core eudicots. Here, we report the identification of a core eudicot-specific TSB-like pseudoenzyme that itself lacks enzymatic activity but allosterically activates TSA for efficient production of indole for BXD biosynthesis and, more broadly, for plant defense and communication.

**Fig. 1.**
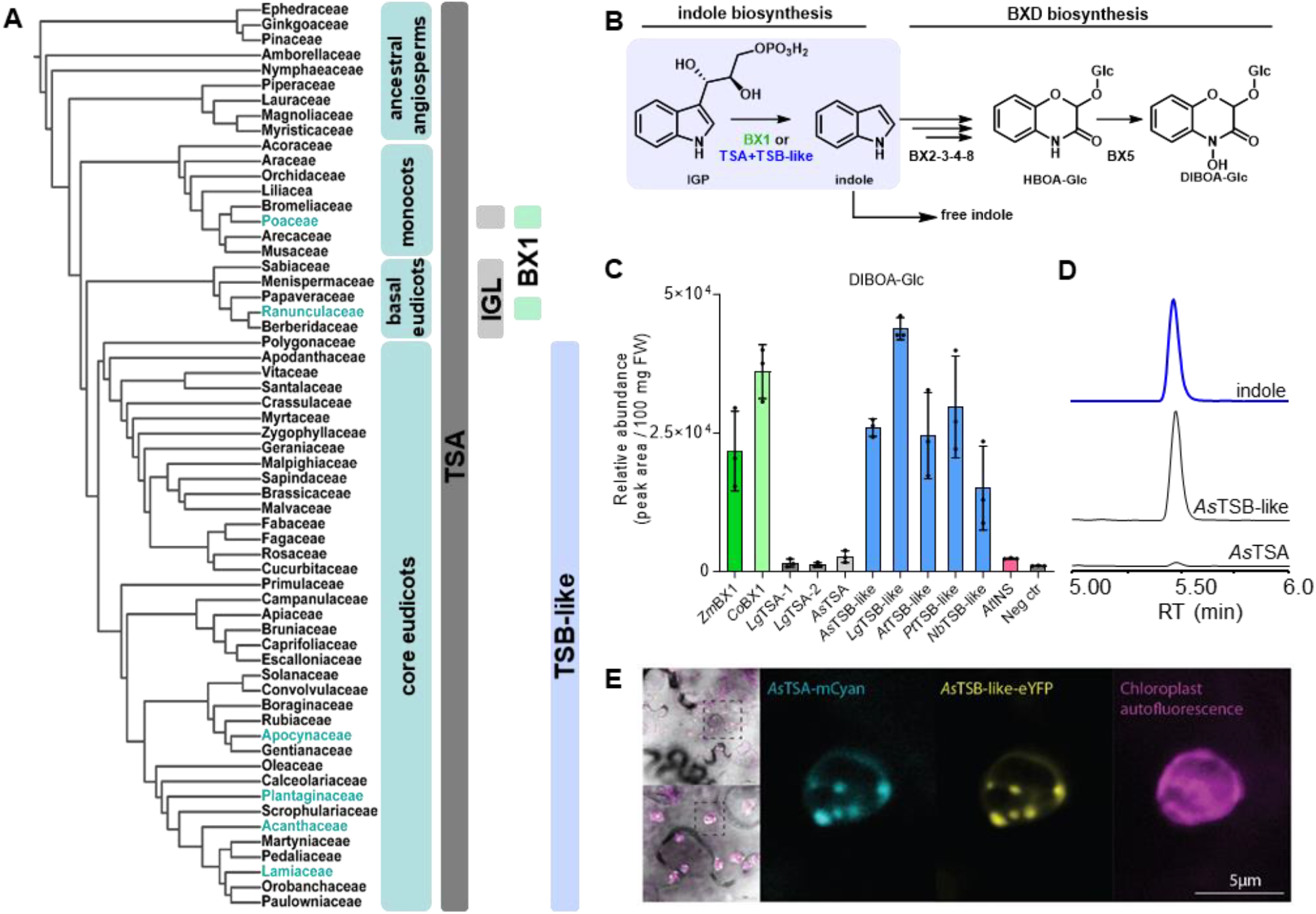
Plants have several ways to produce indole. **A**) Occurrence of different indole biosynthetic enzymes in plants. Plant families with BXD-producing species are colored in green. **B**) Indole as a precursor for benzoxazinoids is produced by BX1 in the monocots and basal dicots or, as reported herein, by TSA – TSB-like complexes in the core eudicots. **C**) BX1 and TSB-like from different species provide indole as a precursor for benzoxazinoids. BX1 genes from *Zea mays* (*Zm*) and *Consolida orientalis* (*Co*), IGL genes from *Lamium galeobdolon* (*Lg*) and *Aphelandra squarrosa* (*As*), TSB-like genes from *A. squarrosa* (*As*), *L. galeobdolon* (*Lg*), *Populus trichocarpa* (*Pt*), *Nicotiana benthamiana* (*Nb*) and *Arabidopsis thaliana* (*At*), and the INS gene from *A. thaliana* (*At*) were transiently expressed in *N. benthamiana* along with the BXD biosynthetic genes *ZmBx1, ZmBx2, ZmBx3, ZmBx4, ZmBx5*, and *ZmBx8* from maize. Plant material was extracted with methanol and analyzed using liquid chromatography-time-of-flight mass spectrometry. Average accumulation of the BXD product DIBOA-Glc (n = 3) and SEM are displayed. **D**) *As*TSB-like promotes the formation of indole. Transient expression of *AsTSB-like* in *N. benthamiana* leads to higher indole accumulation compared to *AsTSA*. Plant material extracted with MeOH was analyzed through liquid chromatography-tandem mass spectrometry. **E**) *AsTSA* and *AsTSB-like* transiently expressed in *N. benthamiana* co-localize in the chloroplast. *As*TrpA-mCyan, *As*TrpB-like-eYFP, and chloroplastic autofluorescence are displayed.

### TSB-like is widespread in the core eudicots and promotes the formation of free indole

In previous work, we have elucidated the BXD biosynthetic pathway in *A. squarrosa* and *L. galeobdolon* (*16, 22*), but the mechanism of the first committed step of the pathway, indole formation, remained elusive (Fig. 1B and fig. S1B). When the pathway was reconstituted by transiently expressing the biosynthetic genes in the heterologous host *Nicotiana benthamiana*, it was observed that putative IGL/BX1 from *L. galeobdolon* and *A. squarrosa* were unable to support the formation of BXD when combined with downstream BXD pathway genes. In contrast, substitution of the *L. galeobdolon* / *A. squarrosa* putative IGL/BX1 genes with BX1 from *Z. mays* or *Consolida orientalis* led to robust production of BXD (Fig. 1C). We therefore concluded that *A. squarrosa* and *L. galeobdolon* do not utilize IGL/BX1 for BXD biosynthesis and must have an alternative mechanism for production of indole. To uncover this mechanism, we screened the *A. squarrosa* and *L. galeobdolon* transcriptomes for genes that are co-expressed with the previously reported BXD pathway genes. This approach revealed a gene in both species that was similar (57% nucleotide sequence identity on average) to *TSB*, named *AsTSB-like* and *LgTSB-like*, respectively (fig. S1C, D and E). Phylogenetic analysis showed that *As*TSB-like and *Lg*TSB-like formed a clade well separated from those of TSB and TSB type II, which are both known to catalyze formation of tryptophan (fig. S2A) (*23, 24*). However, the function of TSB-like was unknown. Transient expression of *AsTSB-like* or *LgTSB-like* together with the maize BXD pathway genes in *N. benthamiana* resulted in efficient BXD biosynthesis (Fig. 1C and D). In contrast, no BXD were formed when the canonical *AsTSB* or *AsTSB* type II gene was used (fig. S2B). Interestingly, a broader BLAST analysis showed that TSB-like are not only present in the BXD-producing species *A. squarrosa* and *L. galeobdolon* but are widespread in all core eudicots examined (Fig. 1A and fig. S2A). Notably, no TSB-like gene was found in monocotyledons and basal eudicotyledons (fig. S2A), which are known to possess functional IGLs for the formation of free indole (Fig. 1A, fig. S1A and fig. S2A) (*16, 18, 20*). Testing of additional *TSB-like* genes from non-BXD-producing species of the core eudicots, including *N. benthamiana, Populus trichocarpa*, and *Arabidopsis thaliana*, showed that all, when transiently expressed in *N. benthamiana*, resulted in indole production (Fig. 1C).

### TSB-like is a pseudoenzyme that allosterically activates TSA for indole production

Because TSA and canonical TSB are known to form a protein complex and mutually activate each other, we hypothesized that the formation of indole promoted by TSB-like might be associated with TSA. Indeed, when recombinant *As*TSB-like protein was incubated with *As*TSA protein and the substrates IGP and L-serine, indole, but not tryptophan, was produced. Conversely, in reactions containing *As*TSA and canonical *As*TSB, tryptophan was the only product. Each enzyme showed negligible activity when assayed alone (Fig. 2A and B), indicating that *As*TSB-like acts as allosteric activator for *As*TSA. Allosteric activation of TSA by TSB-like was also observed for other species, including *A. thaliana, N. benthamiana, P. trichocarpa*, and *L. galeobdolon* (fig. S3A). In species harboring two TSA genes (*e*.*g. L. galeobdolon*) (*16*), allosteric activation of both homologues by TSB-like was observed (fig. S3A). Notably, *A. thaliana* (*25*) and other Brassicaceae plants have both a TSA and a TSA-like enzyme (INS), where the TSA-like enzyme has been proposed to produce indole in a tryptophan-independent auxin pathway. We also showed allosteric activation of both *At*TSA/TSA-like by TSB-like (fig. S3A). To test whether TSB-like and TSA form a protein complex that is characteristic of this allosteric activation, we performed co-purification assays with different combinations of His-tagged or non-tagged recombinant proteins. Incubation of His-tagged *As*TSB-like with untagged *As*TSA or vice-versa followed by nickel affinity purification always resulted in purification of both proteins, regardless of whether the His-tag was fused to the C-or N-terminus of the protein (Fig. 2C and fig. S3B and C). This demonstrates that *As*TSB-like forms a complex with *As*TSA that is stable even under the conditions of the in vitro purification procedure. Competition assays (in vitro and in *N. benthamiana*) also suggested that TSB-like and TSA form a complex. In vitro, incubation of TSA and TSB-like with increasing amounts of TSB resulted in a progressive reduction of indole and increase of tryptophan accumulation (Fig. 2Di and ii). Incubation of TSA and TSB with increasing amounts of TSB-like resulted in increased indole accumulation, and in this case, minor levels of tryptophan (Fig. 2D iii and iv), most likely due to the TSB-like/TSB product-substrate cross talk (fig. S4A and B). In *N. benthamiana*, co-expression of *AsTSB-like* together with *AsTSB* and *ZmBx2-4* resulted in an approximately 50% reduction in the amount of BXD produced compared to the control, which expressed only *AsTSB-like* and *ZmBx2-4* (Fig. 2E). Moreover, the subcellular localization of *As*TSA and *As*TSB-like, as evidenced by expression of fluorescence-tagged proteins in *N. benthamiana*, indicated that both proteins were localized close to each other in the chloroplasts (Fig. 1E and fig. S5). Taken together, our results suggest that TSB-like has no enzymatic function but instead binds to TSA, thereby activating this enzyme for efficient indole production (Fig. 2F).

**Fig. 2.**
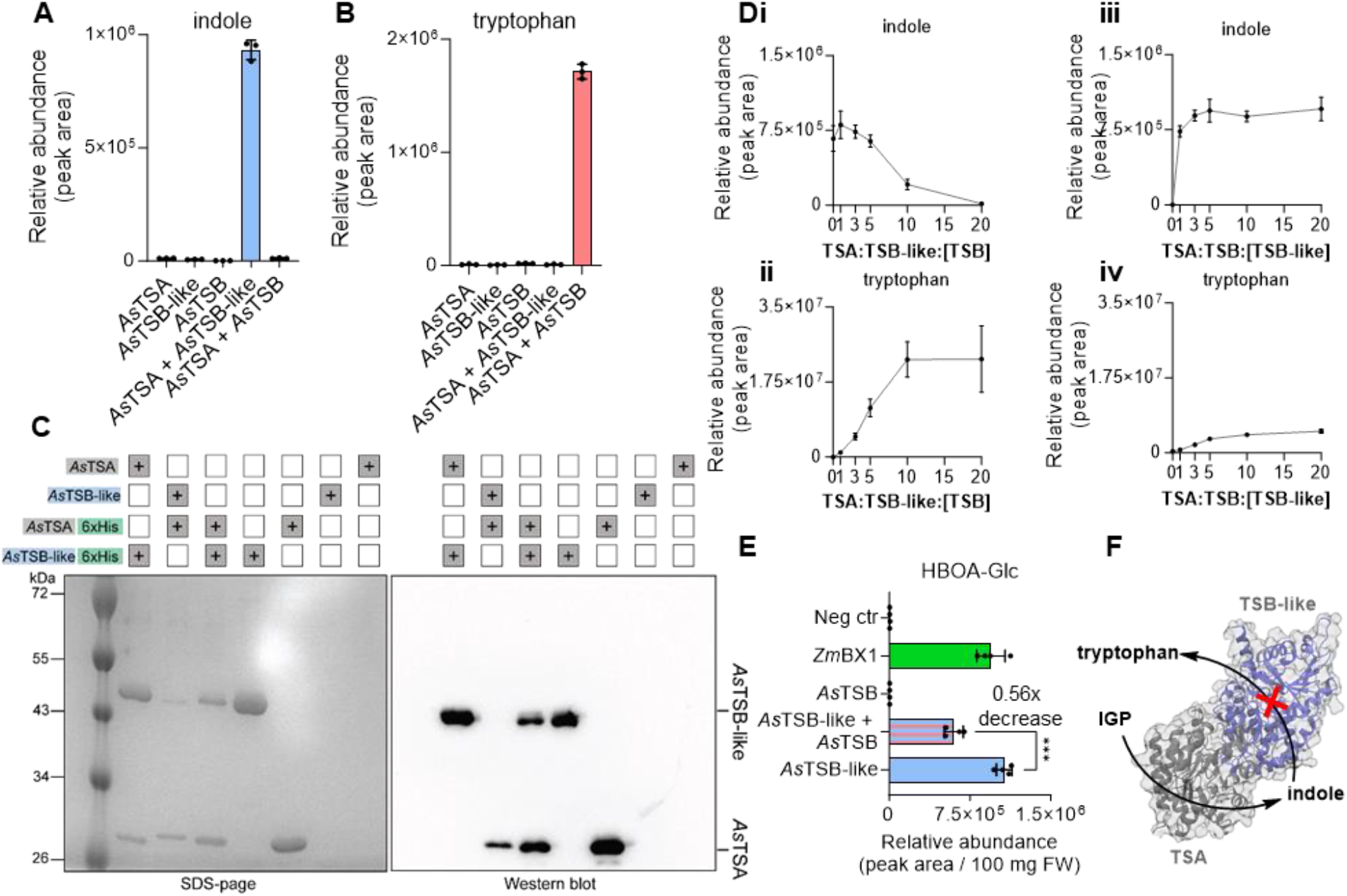
TSB-like binds to TSA, triggering allosteric activation, resulting in indole formation. **A** and **B**) *As*TSB-like induces indole formation by TSA but does not produce tryptophan in vitro. Proteins were expressed in *Escherichia coli*, purified, and assayed with the substrates IGP and L-Ser. Accumulation of indole and tryptophan was measures by liquid chromatography tandem mass spectrometry (LC-MS/MS). Average (n = 3) and SD are displayed. **C**) TSB-like forms a complex with TSA. *E. coli* cultures expressing C-terminal His-tagged or untagged TSA and TSB-like were mixed and His-tagged proteins were retrieved trough affinity purification. Untagged TSA or TSB-like could be co-purified with the corresponding tagged partner as shown in SDS-page and Western blot. **D**) *As*TSB and *As*TSB-like compete for *As*TSA in vitro. Proteins were expressed in *E. coli*, purified, and assayed with the substrates IGP and L-Ser. **i** and **ii**) Equimolar concentrations of TSA and TSB-like were incubated with increasing concentrations of TSB resulting in an increased accumulation of tryptophan and reduced accumulation of indole. **iii** and **iv**) Equimolar concentrations of TSA and TSB were incubated with increasing concentrations of TSB-like resulting in increased accumulation of indole but also of tryptophan. The x axis shows the stochiometric ratio of the protein in brackets. Indole and tryptophan accumulation was measured on LC-MS/MS. Average (n = 3) and SD are displayed. **E**) *As*TSB and *As*TSB-like compete for TSA in *N. benthamiana*. Coexpression of *AsTSB* and *AsTSB-like* with *ZmBx2* – *4* and *ZmBx8* resulted in a decrease of ∼ 50% of the benzoxazinoid HBOA-Glc produced. Plant material was extracted with MeOH and analyzed using liquid chromatography-time-of-flight mass spectrometry. Average (n = 4) and SD are displayed. **F**) Schematic of how TSB-like and TSA produce indole.

### Two conserved residues mediate the different functionalities of TSB-like

Phylogenetic analysis suggested that TSB-like most likely evolved by gene duplication and neofunctionalization of a canonical TSB gene (fig. S2). To understand how TSB-like lost tryptophan synthase activity but retained the capacity to allosterically activate TSA, we identified active site residues that were consistently different between TSB-like and TSB in all species examined (fig. S6A, B and C). We identified a glutamate residue that has been shown to be essential for tryptophan formation and an aspartate residue proposed to play a role in TSB activation (*10*–*12, 26, 27*). In TSB-like, this glutamate residue is almost always replaced by alanine (A195 in *As*TSB-like), with the only exception found in TSB-like from Solanaceae species, which instead contained a serine or a proline at this position (fig. S7A and fig. S8A and B). The aspartate residue was universally replaced by glutamate in TSB-like (E388 in *As*TSB-like) (Fig. 3A, B and C). Mutation of this alanine (A195) in *As*TSB-like to glutamate resulted in restoration of tryptophan biosynthetic activity, although the *As*TSA/*As*TSB-like A195E complex still produced significant amounts of indole (Fig. 3D and fig. S7C). Mutation of glutamate 388 to aspartate in *As*TSB-like resulted in reduced indole formation though tryptophan biosynthesis was still not observed (Fig. 3D and fig. S7B, C). The double mutant *As*TSB-like A195E - E388D in combination with *As*TSA produced high levels of tryptophan relative to indole (Fig. 3D, fig. S7B and C). Transient expression of the double mutant in *N. benthamiana* showed that TSB-like activity was almost completely eliminated in planta, as evidenced by the lack of indole production (Fig. 3E). Introducing the reverse mutations into *As*TSB (*As*TSB E190A and AsTSB E190A D386E) resulted in almost non-functional proteins (fig. S9, S10). Mutagenesis of additional residues in or near the active site did not lead to indole production, suggesting that mutation of residues outside the active site is also essential (fig. S10A and B).

**Fig. 3.**
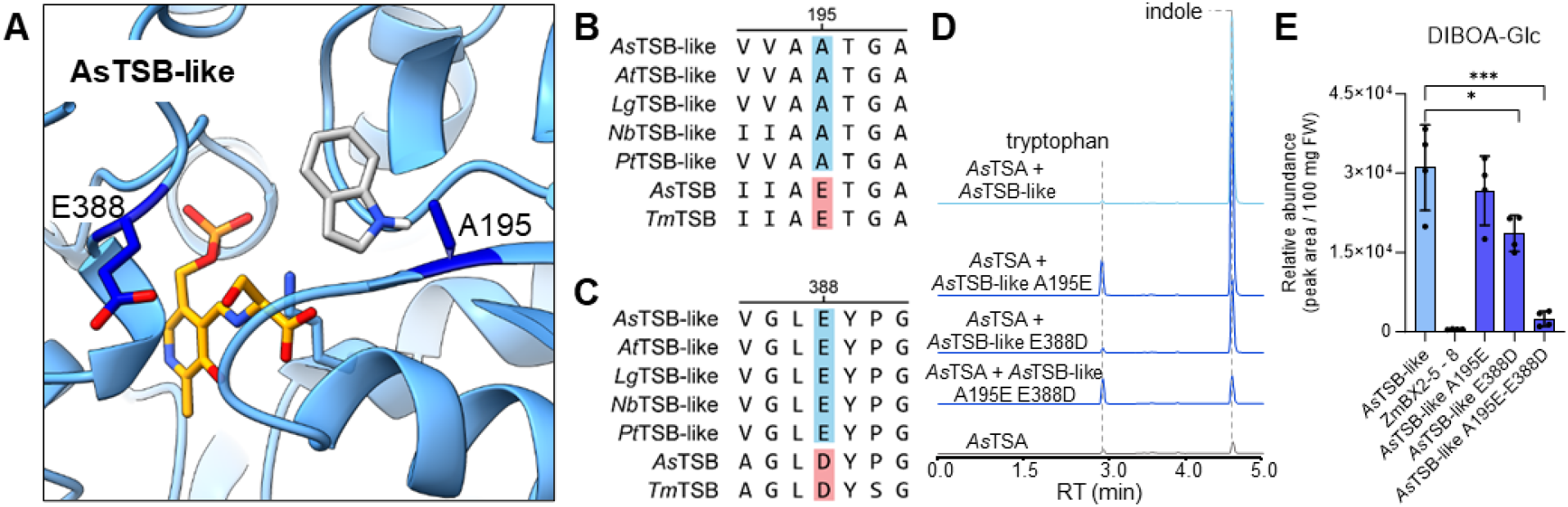
A195 and E388 play a role in AsTSB-like activity. **A**) **Model of the AsTSB-like active** site showing A195 and E388. The structure of *As*TSB-like was modelled on the crystal structure of *Salmonella thypi* TSB (PDB: 7JMQ) in open conformation with pyridoxal phosphate (yellow). Indole (gray) was docked in silico. **B** and **C**) Alignment displaying the conservation of A195 and E388. TSB-like from *A. squarrosa, A. thaliana, L. galeobdolon, N. benthamiana*, and *P. trichocarpa* were compared with *A. squarrosa* and *T. maritima* TSB. Sequences were aligned using the MUSCLE algorithm and the residues of interest were highlighted in blue for TSB-like and in red for TSB. **D**) *As*TSB-like mutants resulted in gain of tryptophan biosynthetic activity and reduction of indole biosynthetic activity. *As*TSB-like A195E gained tryptophan biosynthetic activity but retained indole biosynthetic activity. *As*TSB-like E388D showed highly reduced indole biosynthetic activity and no gain of tryptophan biosynthetic activity. *As*TSB-like A195E E388D showed gain in tryptophan biosynthetic activity and reduction of indole biosynthetic activity. Proteins were expressed in *E. coli*, purified, and assayed on IGP and L-Ser. Reaction products were analyzed on liquid chromatography/tandem mass spectrometry. **E**) The double mutant *As*TrpB-like A195E – E388D showed highly reduced indole production in *N. benthamiana* as evidenced by monitoring production of the benzoxazinoid DIBOA-Glc. *As*TrpB-like mutants were transiently expressed in *N. benthamiana*. Plant samples were extracted with MeOH and analyzed using liquid chromatography/time-of-flight mass spectrometry. Means (n = 4) and SD are displayed. * = p<0.05, *** = p<0.0005.

### TSB-like is involved in plant defense and signaling

Indole is a widespread plant volatile that is often released in response to herbivory or as a characteristic floral scent component. *N. benthamiana* emits indole after herbivory attack (*28*) (Fig. 4A). We could show that, along with indole emission, *TSB-like* expression, but not *TSA* expression, was strongly up-regulated in *N. benthamiana* upon herbivory (Fig. 4B), suggesting that TSB-like promotes indole formation in response to biotic stress. A meta-analysis of literature data across core eudicot species for which both metabolome and transcriptome data were available, revealed that both herbivory-induced and floral scent-related indole emissions were always accompanied by an up-regulation of *TSB-like* expression, whereas *TSA* expression remained unchanged or showed smaller fold change differences compared to TSB-like (fig. S11). These observations are consistent with a recently reported study from tea that showed the upregulation of a TSB-like protein after herbivore attack, and that this protein interacts with TSA (*29*). Along with the absence of IGL/ BX1 genes, these data suggest that indole emission in core eudicots is dependent on the action of TSB-like.

**Fig. 4.**
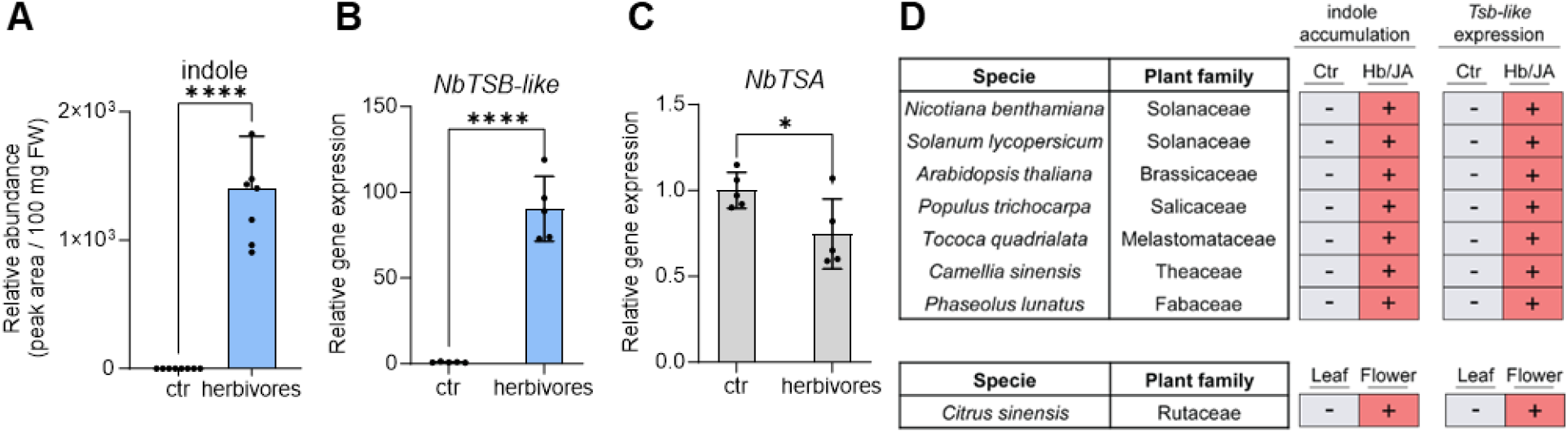
Indole accumulation/emission is accompanied with *TSB-like* expression in the core eudicots. **A)** *Nicotiana benthamiana* plants treated with *Spodoptera littoralis* caterpillars accumulated indole. Plants were exposed to herbivory for 17 h and methanolic leaf extracts were analyzed using liquid chromatography-tandem mass spectrometry. Average (n = 8) and SD are shown (**** = p < 0.0001). **B** and **C**) Herbivory by *S. littoralis* caterpillars induces the expression of *NbTSB-like* and not *NbTSA*. Transcript accumulation of *NbTSB-like* and *NbTSA* was measured by quantitative PCR. Average (n = 3) and SD are shown (**** = p < 0.0001, * = p < 0.05). D) Meta-analysis showing the accumulation/emission of indole and the expression of *TSB-like* in different families of the core eudicots upon herbivory (Hb) or jasmonic acid (JA) treatment. (+) indicates an increased accumulation/emission of indole or *TSB-like* expression compared to the control (Table S1).

## Conclusion

Despite the biological importance of indole in plant defense and communication, the mechanism underlying its formation in the vast and economically important clade of the core eudicots had remained unknown. Recently, the widespread distribution of *TSB-like* genes has been noticed (*30*), but the function of these genes in the plant kingdom nevertheless remained elusive (fig. S12). In this work we report that the pseudoenzyme TSB-like, a catalytically “dead” paralog of TSB, appears to be responsible for indole biosynthesis in the core eudicots. TSB-like most likely evolved from TSB through a loss of tryptophan biosynthetic activity. The resulting catalytically inactive TSB-like mimics the interaction of TSB with TSA, thereby allosterically activating TSA to allow indole biosynthesis, but without subsequent conversion to tryptophan. Therefore, TSB-like has evolved to serve as a switch that toggles between tryptophan and indole biosynthesis by hijacking the pre-existing TSA. Although pseudoenzymes can be challenging to discover, recent work has highlighted that these proteins play essential roles in a number of biological processes such as vitamin B6 biosynthesis (*31*), alkaloid biosynthesis (*32*), and starch breakdown (*33, 34*). In summary, we report the biosynthesis of indole, a fundamental part of the plant defense response (*35*), in core eudicots. The discovery of TSB-like may allow modulation of indole emission by targeted breeding or genetic manipulation, subsequently having a profound impact on food security in economically important eudicots including *e*.*g*. tomato, olive, and coffee.

## Acknowledgments

We thank Maritta Kunert and Sarah Heinicke for help with liquid chromatography/mass spectrometry analyses and the greenhouse team of the Max Plank Institute for Chemical Ecology, especially Danny Kessler and Franz Kaltofen, for rearing the plants. We also thank Susan Schlueter for help with cloning and expression of genes. We thank Dagny Grzech, Chloe Langley, Gabriel Titchiner, and Maite Colinas for helpful discussions.

## Author contributions

Conceptualization: MF, TGK, SEO

Methodology: MF, TGK

Investigation: MF, HS, VG, KL

Visualization: MF

Funding acquisition: SEO

Project administration: TGK, SEO

Supervision: TGK, SEO

Writing – original draft: MF, TGK, SEO

Writing – review & editing: MF, TGK, SEO

## Data and materials availability

Genes described in this study were deposited in NCBI GenBank with the accession numbers given in Table S2. All other data are available in the main text or supplementary material.

