## Supplemental methods and figures for "Highjacked by a pseudoenzyme: How eudicot plants make indole"

**The PDF file includes:**

Materials and Methods
Figs. S1 to S12
Tables S1 to S4
References (*36-47*)

**Other Supplementary Materials for this manuscript include the following:**

Data S1 to S3

### Materials and Methods

#### Chemicals

All chemicals used in this study were purchased molecular biology grade or higher from Sigma Aldrich, Thermo Fisher, or Tokyo Chemical Industry (TCI) unless otherwise stated. Benzoxazinoid (BXD) standards were synthesized or isolated as reported in *(22)*.

#### Plant material and growth

*Aphelandra squarrosa* plants were cultivated in a greenhouse under a 14-hour light / 10-hour dark photoperiod, with temperatures maintained at 21-25°C during the day and 16-22°C at night, and relative humidity ranging from 40-70%. *Lamium galeobdolon* plants were grown in similar conditions but with a 16-hour light / 8-hour dark photoperiod, daytime temperatures of 20-24°C, nighttime temperatures of 16-20°C, and relative humidity between 45-60%. *Nicotiana benthamiana* plants were grown on a 16-hour light / 8-hour dark photoperiod at a constant temperature of 22°C and 60% relative humidity. *N. benthamiana* plants were grown for 3 weeks prior to gene candidate infiltration.

#### Plant metabolite extraction

Collected plant material was snap-frozen in liquid nitrogen and ground to fine powder with 3 mm Tungsten Carbide Beads using a TissueLyser (Quiagen) or, when more material was needed, liquid nitrogen frozen samples were ground to a fine powder in a pre-chilled mortar. Tissue samples (100 mg ± 5%) were extracted with 500 µl MeOH (LC-MS grade). Samples were vortexed vigorously and then incubated at 25°C, shaking for 15 min. Samples were then centrifuged at maximum speed in a tabletop centrifuge for 15 minutes before filtering with a 0.22 µm PTFE syringe filter for LC-MS analysis.

#### Gene candidate identification

Gene candidates were selected from the previously published *A. squarrosa* and *L. galeobdolon* transcriptomes (BioProject accession PRJNA967136) assembled as reported in *(22)*. Pearson co-expression correlation analyses were performed in Excel. TSB-like candidates in other species were identified based on homology by performing BLAST analysis on public databases: NCBI, SolGenomics, Citrus Genome Database, 1KP, NbenBase.

#### Cloning

Total RNA was extracted from ground plant tissue using the RNeasy Plant Mini Kit (Quiagen) including an on-column DNAse digestion step according to manufacturer instructions. cDNA was synthesized from total RNA using SuperSCript IV VILO Master Mix (Thermo Fisher Scientific), according to manufacturer instructions. Genes were amplified from cDNA using Platinum SuperFi II PCR Master Mix (Thermo Fisher Scientific). Synthetic genes, when used, were ordered from Twist Bioscience and used as a template for PCR amplification. PCR products were purified using DNA Clean and Concentrator-5 (Zymo) or Zymoclean Gel DNA Recovery Kit (Zymo). Amplified genes were inserted with In-Phusion HD Cloning (Takara Bio) in p3Ω1 vector (*BsaI*-HF digested) for expression in *N. benthamiana*. For expression in *Escherichia coli*, the following vectors were used: pOPINF (*HindIII*-HF/*KpnI*-HF digested) for N-terminal His-tagged sequences, pOPINE (*NcoI*-HF and *KpnI*-HF digested) for C-terminal His-tagged sequences, and pET28a for alternative N-terminal (*BamHI*-HF and *NotI*-HF) or C-terminal (*NcoI*-HF and *XhoI*-HF digested) His-tagging.

For subcellular localization studies, *AsTSA* and *AsTSB-like* were cloned with a C-terminal fused fluorescent protein. *AsTSB-like* was cloned with C-terminal eYFP fluorescent protein and *AsTSA* was cloned with C-terminal mCeruleans fluorescence protein. The gene of interest was separated from the fluorescent protein by a AGCGGC linker. The fusion constructs were cloned under the control of the strong constitutive *Solanum lycopersicum* Ubiquitin10 (*Sl*Ubq10) promoter and terminator in 3$\alpha$1 vector through Golden Braid using *BsaI*-HF and T4 DNA *(36)*. Vectors harboring the sequences of interest were transformed in *E. coli* Top10 with the heat shock method and plated on LB-agar plates with appropriate antibiotic selection. Overnight colonies were inoculated in liquid LB with appropriate selection and incubated at 37°C, 250rpm for 6-7h. Plasmid DNA was isolated using Wizard Plus SV Minipreps DNA Purification System Kit (Promega) following manufacturer instructions. Each construct was checked through Sanger sequencing to verify sequence of the inserted gene. All the primers used in this study are reported in Table S3.

***Agrobacterium tumefaciens* mediated transient transformation of *N. benthamiana***

Electrocompetent *Agrobacterium tumefaciens* GV3101 (Goldbio) cells were mixed with 50 ng of sequence-confirmed plasmid and incubated on ice for 15 minutes. The cells were electroporated using a BioRad Micropulser. The transformed cells were recovered in 1 mL of LB medium and incubated at 28°C, 200 rpm for 3 hours before plating on LB-agar plates containing the appropriate selection marker. Plates were incubated at 28°C for 48 hours. Single colonies were inoculated into liquid LB medium with the appropriate selection and incubated overnight at 28°C, 200 rpm. For *N. benthamiana* transient transformation, the overnight cultures were pelleted by centrifugation at 4000 rpm for 10 minutes at 14°C. The cell pellet was resuspended in infiltration medium (10 mM MES, 10 mM $MgCl_2$, 100 µM acetosyringone, pH 5.7) to an OD600 of 0.6-0.7 and incubated at 28°C, 200 rpm for 1.5 hours. Equal volumes of the prepared infiltration solutions were mixed to achieve the desired transformation mix containing each construct at OD600 of 0.1. The transformation mix was infiltrated into the abaxial side of 3 week old *N. benthamiana* leaves using a needleless 1 mL syringe. The infiltrated plants were maintained in a growth chamber under growth lights up to 5 days post-infiltration, when samples were collected. In all transformations, a construct encoding the silencing repressor protein p19 was co-infiltrated to enhance expression.

**Small scale heterologous expression of candidate genes in *E. coli***

Gene candidates were expressed as previously described in **Florean et al., 2023** with minor modifications. In brief, *E. co*li DE3 (ThermoFisher Scientific) cells were transformed with sequence-confirmed plasmids using the heat-shock method, plated on LB-agar plates with appropriate selection and grown at 37°C overnight. Single colonies were inoculated in liquid LB medium with selection and grown at 37°C, 250 rpm, overnight. The seed culture (1 mL) was used to inoculate 100 mL 2 x YT medium with selection and the culture was grown at 37°C, 250 rpm shaking, until OD600 = 0.5-0.6. Cultures were then incubated at 18°C, 250 rpm, for 20 minutes before addition of 500 µM IPTG. Induced cultures were incubated at 18°C, 250 rpm, overnight. Cultures expressing TSB and TSB-like were retrieved by centrifugation (4000 x g, 4°C, 15 minutes) and resuspended in A1 buffer (50mM TRIS-HCl, 50mM glycine, 5% v/v glycerol, 0.5 M NaCl, 20 mM imidazole, pH = 8) with 0.2 g/L lysozyme, 1 tablet / 50 mL EDTA-free protease inhibitor and 100 µM pyridoxal phosphate (PLP) and disrupted by sonication on ice (Bandelin UW 2070). Cell debris was removed by centrifugation at 35,000 × g at 4°C for 20 min and His-tagged proteins were purified from the supernatant using NiNTA agarose (Qiagen) beads according to the manufacturer's instructions. Proteins were eluted using elution buffer B1 (A1 buffer + 500 mM imidazole, pH = 8). Ultimately, elution buffer was exchanged for protein storage buffer (20 mM HEPES, 150 mM NaCl, pH 7.5, 10% glycerol) using Amicon concentrator columns (Merck Millipore). Protein were aliquoted and stored at -20°C.

**Large scale heterologous expression of candidate genes in *E. coli***

For large scale heterologous expression, 1 L of 2 x YT media was inoculated with 10 mL of seed culture and induced as described above. Pelleted cells were resuspended in 20 mL of A1 buffer with 0.2 g/L lysozyme, 1 tablet / 50 mL EDTA-free protease inhibitor and 100 μM PLP. Cells were disrupted by sonication on ice (Bandelin UW 2070). Cell debris was removed by centrifugation at 35,000 × g at 4°C for 20 minutes and His-tagged proteins were purified on an ÄKTA pure FPLC system (GE Healthcare) equipped with a 5 mL HisTrap column (Cytiva). The FPLC system was programmed as described in *(37)*. In brief, the column was equilibrated with 5x column volumes of buffer A1. The protein sample was loaded at a flow rate of 2 mL / min. Subsequently the column was washed with buffer A1 (flow rate = 5 mL / minute) for a total of 10 column volumes. The protein was eluted with 5x column volumes of buffer B1 and the elution monitored using UV absorption at 280nm.

**Protein concentration determination**

Concentration of PLP-dependent protein was calculated using Pierce Rapid Gold BCA Protein Assay Kit (Thermo Fisher Scientific) following manufacturer instructions. Plates were read on a CLARIOstar Plus (BMG Labtech) plate reader. Concentration of non-PLP dependent proteins was determined spectrophotometrically measuring absorbance at 280 nm on a IMPLEN Nanodrop.

**SDS-Page and Western blot**

SDS-page analyses were performed using Novex 12%, Tris-Glycine Plus WedgeWell gels (Invitrogen) according to manufacturer instructions. Gels for SDS-page were stained with Quick Coomassie Stain (Serva). Gels for Western Blot analysis were transferred on a Power Blotter Select Transfer Stack PVDF Mini Size membrane using Power Blotter XL transfer station (Invitrogen). Blotted membranes were blocked in TBS + 1 mL/L Tween buffer (TBST) + 5% (w/v) skimmed milk at room temperature for 1h. Blocking solution was removed and incubated in TBST + 3% (w/v) skimmed milk and anti-Histidine antibody coupled with Horseradish peroxidase (BioRad) as per manufacturer instructions. Western blots were imaged with Clarity Western ECL Substrate (BioRad) as per manufacturer instructions.

**IGP *in vitro* biosynthesis**

Indole-3-glycerol phosphate (IGP) was synthesized *in vitro* as described by *(38)* by incubating recombinantly purified *E. coli* phosphoribosyl transferase (TrpD) and phosphoribosyl anthranilate isomerase – indole-3-glycerol phosphate synthase (TrpF-TrpC fusion gene) with 0.5 mM $MgCl_2$, 0.4 mM DTT, 3 mM anthranilic acid, and 3 mM 5-phospho-D-ribose-diphosphate. The reaction was performed in $KPO_4$ buffer, 25 mM, pH = 7.5 at 30°C, shaking for 1 hour. The reaction was stopped by heat inactivation at 95°C for 10 minutes and proteins were precipitated by centrifugation. IGP was stored at -20°C and used within one day of synthesis.

***In vitro* assays**

*In vitro* assays for indole and tryptophan biosynthesis were performed in $KPO_4$ buffer, 25 mM, pH = 7.5 with 10 nM of each protein, and saturating concentrations of IGP, 1 mM L-serine, 0.2 mM PLP. Reactions were started by addition of substrate. The reactions were incubated 15 minutes at 30°C, 300 rpm and quenched by addition of one isovolume of MeOH. Proteins were precipitated by centrifugation and samples were analyzed through LC-MS.
In vitro reactions to check tyrosine biosynthesis were performed in in $KPO_4$ buffer, 25 mM, pH = 7.5 with 50 nM of each protein, 1 mM phenol in DMSO, 1.5 mM L-serine and 0.2 mM PLP. Reactions were started by addition of the substrate and incubated 1 hour at 30°C, 300 rpm. Reactions were quenched by addition of one isovolume of MeOH:1M HCl. Proteins were precipitated by centrifugation and samples were analyzed on LC-qTOF.

**Liquid chromatography-quadrupole time-of-flight mass spectrometry (LC-qTOF-MS) analysis**
Samples were analyzed as described in *(22)* with minor variations. Liquid chromatography-quadrupole time-of-flight mass spectrometry (LC-qTOF-MS) analyses were conducted on a Thermo Scientific UltiMate 3000 ultra-high performance liquid chromatography (UHPLC) system coupled to an Impact II UHR-Q-ToF (Ultra-High Resolution QuadrupoleTime-of-Flight) mass spectrometer (Bruker Daltonics). Chromatographic separation was performed using a reverse-phase Phenomenex Kinetex XB-C18 column (100 x 2.1 mm, 2.6 µm; 100 Å) at 35°C. The mobile phase consisted of water + 0.1% formic acid (A) and acetonitrile (B) run at a 0.3 mL/minute flow. 2 µL of sample were injected. The chromatographic separation was performed starting at 5% B for 1 min, linear gradient from 5% to 50% B in 7 minutes, 100% B for 2.5 minutes, 5% B for 2.5 minutes. Mass spectrometry acquisition was performed in positive or negative electrospray ionization mode depending on the compound of interest as described in *(22)*.

**Liquid chromatography-tandem mass spectrometry (LC-MS/MS) analysis**
Targeted analysis of indole and tryptophan was performed using Thermo Scientific UltiMate 3000 ultra-high performance liquid chromatography (UHPLC) system coupled to a Bruker EVOQ Elite tandem mass spectrometer. Chromatographic separation was performed using a reverse-phase Phenomenex Kinetex XB-C18 column (100 x 2.1 mm, 2.6 µm; 100 Å) at 35°C. The mobile phase consisted of water + 0.1% formic acid (A) and acetonitrile (B) run at a 0.3 mL/minute flow with a sample injection of 1 µL. The chromatographic separation was performed starting at 5% B for 30 seconds, linear gradient from 5% to 70% B in 4 minutes, 100% B for 2 minutes, 5% B for 2 minutes. Mass spectrometry acquisition was performed in positive mode using a heated electrospray ionization source (HESI), with a spray voltage of 4000 V, cone temperature of 350°C, cone gas flow of 20 psi, probe temperature of 400°C, probe gas flow of 45 psi and nebulizer gas flow of 50 psi. Indole and tryptophan were detected using multiple reaction monitoring (MRM) transitions. For indole the transition from 118 *m/z* to 91 *m/z* using a collision energy of 19 eV was used. For tryptophan 205.1 *m/z* to 188 *m/z* with a collision energy of 5 eV, 205.1 *m/z* to 146 *m/z* with a collision energy of 13 eV and 205.1 *m/z* to 118 *m/z* with a collision energy of 23 eV were used. Data was analyzed using Bruker MS Data Review version 8.2.1 software.

**Confocal laser scanning microscopy for subcellular localization analysis**
*A. tumefaciens* strains harboring *AsTSB-like-eYFP* or *AsTSA-mCeruleans* construct were infiltrated in 3 week old *N. benthamiana* plants as described above. Plant leaf disks were analyzed 48 hours post-infiltration. Micrographs of the freshly punched leaf discs were acquired using a

cLSM 880 Axio Imager 2 (Zeiss, Oberkochen, Germany) equipped with a C-Apochromat 40x/1.20 water immersion objective. The leaf discs were water mounted in 3d-printed object slides with 400 µm deep circular wells and covered with a 170 µm thick cover glass. The fluorophores were scanned in two line-sequential tracks with two channels each. The first track contained the excitation with 458 nm Argon laser (10% transmission) for mCyan and 405 nm laser diode (1%) for chlorophyll auto fluorescence combined with a MBS 405 and a MBS 458/514. Emission of mCyan and chlorophyll were detected between 460-499 nm (650 detector gain) and 639-743 nm (650 gain), respectively, with a pinhole adjusted to 1 Airy Unit. The line-sequential second track contained excitation of eYFP with a 514 nm Argon laser (3%) combined with a MBS 458/514, and its emission was detected between 517-597 nm (600 gain). Additionally, the second track contained a transmitted light channel T-PMT (400 gain). The majority of the micrographs were acquired unidirectional with an averaging of 8, a pixel dwell time of 0.76 µs, a resolution of 1024x1024 and a resulting pixel scaling of 50x50 nm.

**Herbivory treatment**

Three to four *Spodoptera littoralis* caterpillar (second to third instar) were starved for 24 hours, then placed on three weeks old *N. benthamiana* leaves and let to feed on the plants for 17 hours. Afterwards, caterpillars were removed and plant tissue was immediately snap-frozen in liquid nitrogen. Tissue was grinded to a fine powder and used for metabolite extraction or qPCR analysis.

**qPCR analysis**

Primers for qRT-PCR analysis were designed to have a $T_m$ of 60°C, a GC content of 40-60%, and a length of 20-21 bases using the primer design software in Geneious Prime (modified Primer3 2.3.7 version) resulting in amplicon sizes between 105 and 134 bp. The specificity of the primers was confirmed by agarose gel electrophoresis, melting curve analysis, and by sequence verification of the cloned PCR amplicons. The efficiencies of the primers (95.7%-103.6%) were determined using a standard curve. Three common housekeeping genes were tested *(39)*. The most stable gene (*PP2A*) according to the standard deviation was used to calculate the relative quantities. All samples were run on a CFX Connect Real-Time PCR Detection System (Bio-Rad Laboratories, Hercules, CA, USA) in an optical 96-well plate. The qRT-PCR were performed with the Biozym Blue S'Green qPCR Kit Separate ROX according to manufacturer instructions. cDNA was diluted 1:10 for analysis. Five biological samples per treatment were analyzed in triplicate. The following PCR conditions were applied for all reactions: Initial incubation at 95°C for 3 min followed by 40 cycles of amplification (95°C for 5 seconds, 60°C for 20 seconds). Reads were taken during the extension step of each cycle and melting curve data were recorded at the end of cycling at 65–95°C. Normalized fold expression was calculated with the ΔΔCP method *(40)*. Data and calculations are reported in Table S4.

**Protein modelling**

Protein models were generated using AlphaFold2 using MMSeq (https://colab.research.google.com/github/sokrypton/ColabFold/blob/main/AlphaFold2.ipynb) with default parameters. Alternatively, models were created by homology modeling using SWISS-MODEL (https://swissmodel.expasy.org/). PLP and ligands were introduced in the models in Pymol by aligning the obtained protein model with crystal structures of orthologous enzymes co-crystalized with PLP and ligands. Protein figures were generated with Chimera X v1.3.

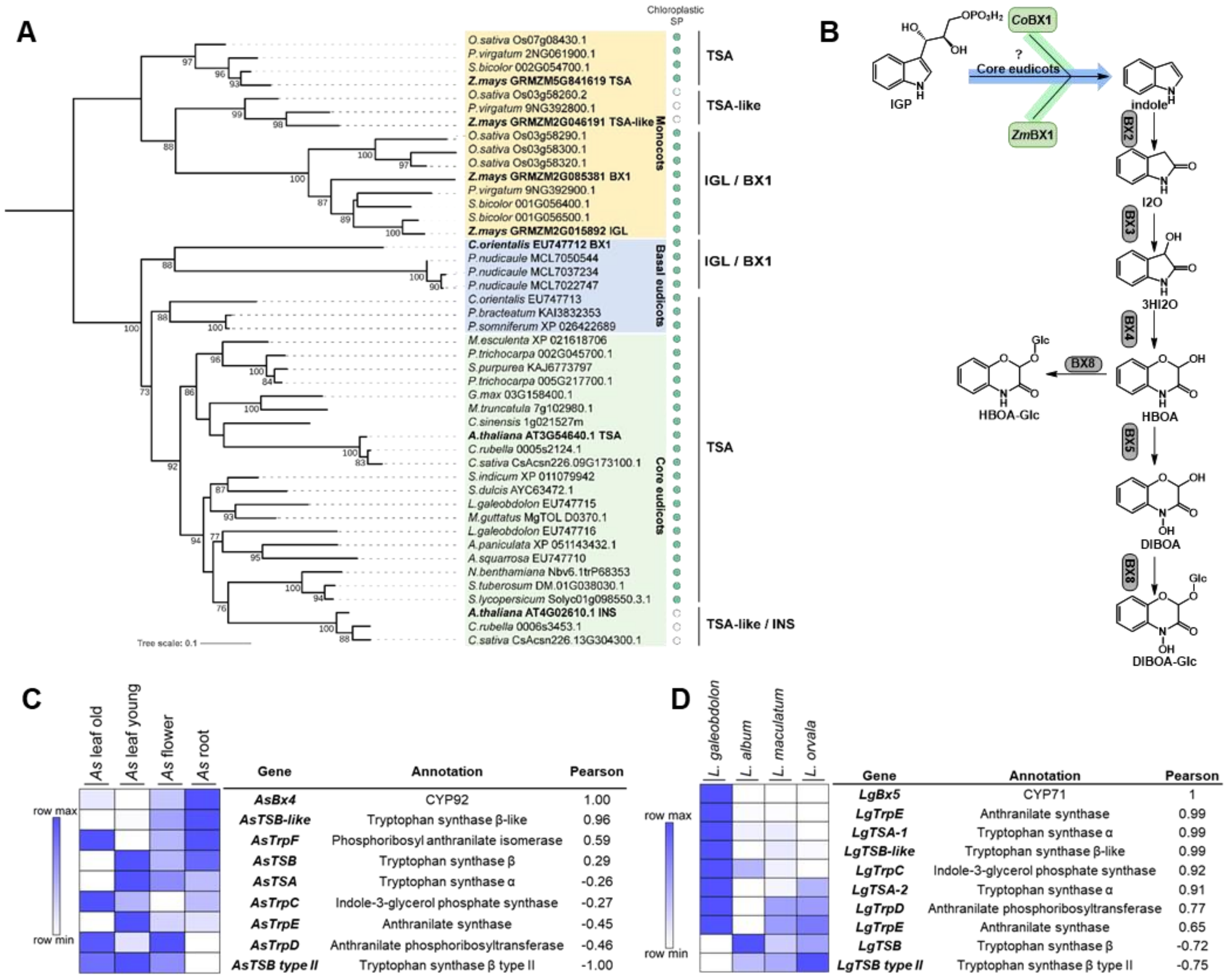


**E**

| % AA seq ID | *Lg*TSB type II | *Lg*TSB | *Lg*TSB-like | *As*TSB type II | *As*TSB | *As*TSB-like |
|---|---|---|---|---|---|---|
| *Lg*TSB type II | - | 21.9 | 23 | 78.8 | 21.2 | 22.1 |
| *Lg*TSB | 21.9 | - | 56.8 | 22.4 | 79.2 | 54.7 |
| *Lg*TSB-like | 23 | 56.8 | - | 23.2 | 54 | 70.8 |
| *As*TSB type II | 78.8 | 22.4 | 23.2 | - | 22.1 | 22.8 |
| *As*TSB | 21.2 | 79.2 | 54 | 22.1 | - | 53.7 |
| *As*TSB-like | 22.1 | 54.7 | 70.8 | 22.8 | 53.7 | - |

**Fig. S1.**

**Indole biosynthesis is initiated by different classes of enzymes. A**) Distribution of TSAs and IGLs among grasses (monocot) and eudicot. Presence (green dot) or absence (white dot) of a chloroplastic localization peptide is reported. Amino acid sequences were aligned with WebPrank and a Maximum Likelihood phylogenetic tree was inferred using iQTree software. The tree was midpoint rooted. Sequences used are reported in Data S1. **B**) BXD biosynthetic pathway up to DIBOA-Glc. **C**) Heatmap displaying the expression of genes involved in indole and tryptophan biosynthesis in *Aphelandra squarrosa* and the corresponding Pearson correlation values with *AsBx4*. **D**) Heatmap displaying the expression of genes involved in indole and tryptophan biosynthesis and the corresponding Pearson correlation values with *LgBX5* in different *Lamium* species. Among the displayed species, only *L. galeobdolon* is a BXD producer. Two *TSA* genes are expressed in *L. galeobdolon*. **E**) Table displaying amino acid sequence identity between TSB-like, TSB, and TSB type II of *A. squarrosa* and *L. galeobdolon*.

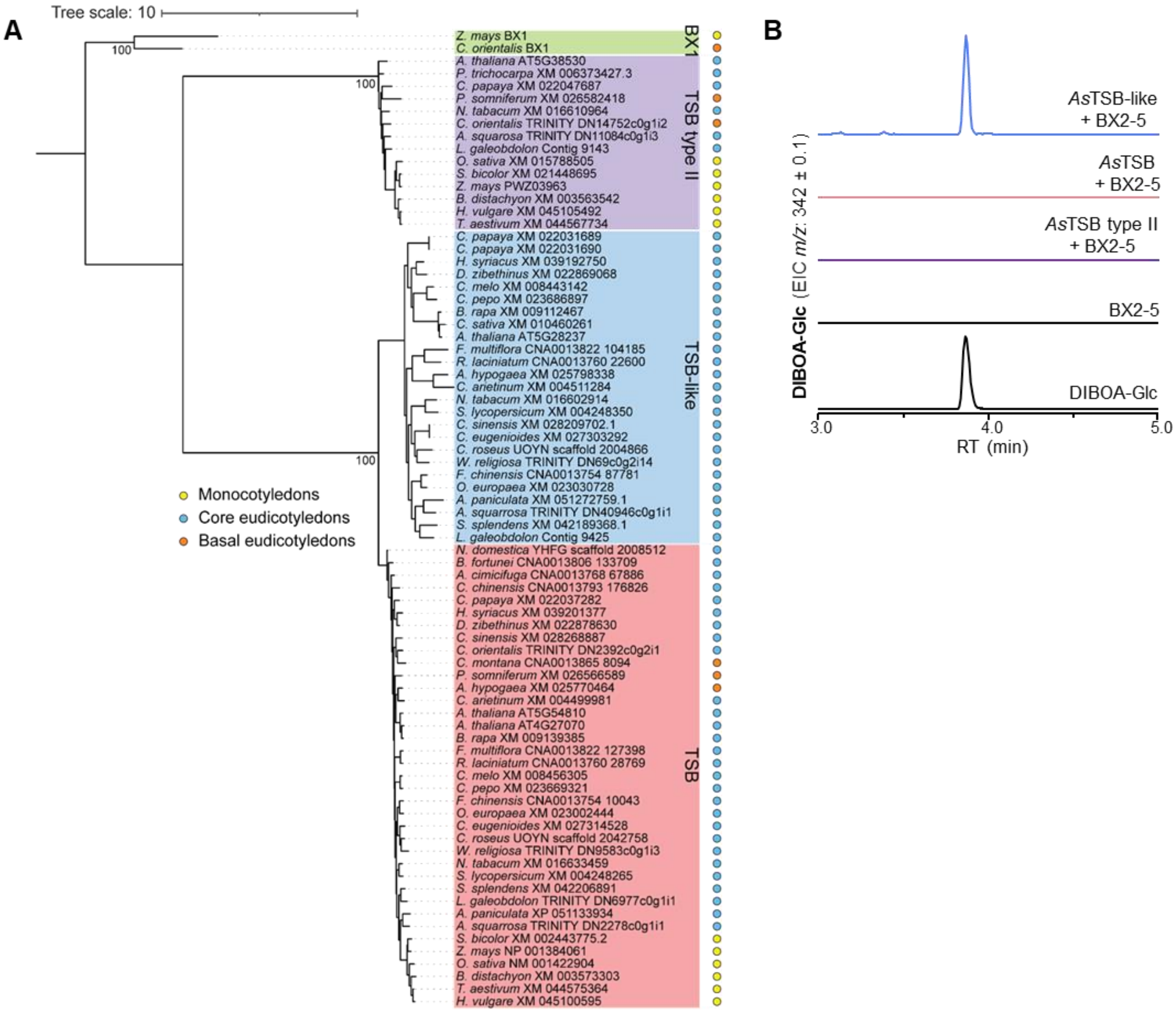


**Fig. S2.**

**TSBs form three distinct clades, and only TSB-like proteins mediate indole formation. A**) TSBs, TSBs-like, and TSB type II form three separate clades. Occurrence of TSBs, TSB-like, and TSB type II in monocotyledons (gray circle), basal eudicot (orange circle), and core eudicot (blue circle) is reported. TSB and TSB type II are present in monocotyledons, basal eudicotyledons, and core eudicot, while TSB-like occur only in core eudicot. Amino acid sequences were aligned with WebPrank algorithm and a Maximum Likelihood tree was inferred using iQTree. The tree was midpoint rooted. Sequences used are reported in Data S2. **B**) AsTSBs were tested for indole biosynthetic activity in *N. benthamiana*. *AsTsB-like*, *AsTsB*, or *AsTsB type II* were coexpressed with *ZmBx2-5*. Leaf methanolic extracts were analyzed using LC-qTOF and DIBOA-Glc accumulation was confirmed with an authentic standard.

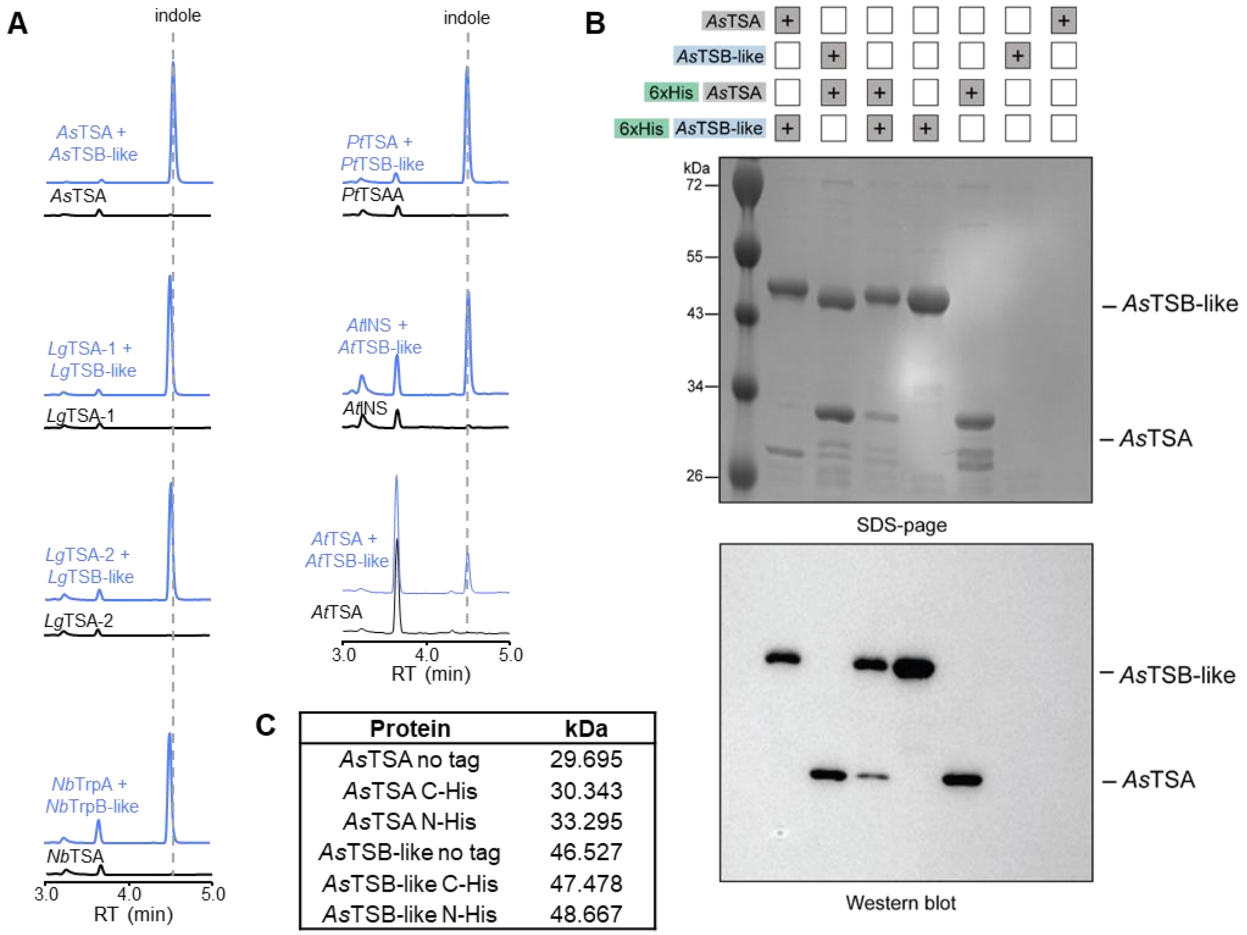


| Protein | kDa |
|---|---|
| *As*TSA no tag | 29.695 |
| *As*TSA C-His | 30.343 |
| *As*TSA N-His | 33.295 |
| *As*TSB-like no tag | 46.527 |
| *As*TSB-like C-His | 47.478 |
| *As*TSB-like N-His | 48.667 |

**Fig. S3.**

**Testing TSA and TSB-like interaction and allosteric activation. A**) Proteins were expressed in *Escherichia coli* and purified TSA/IGL and TSB-like from *Aphelandra squarrosa*, *Lamium galeobdolon*, *Populus trichocarpa*, *Nicotiana benthamiana*, and *Arabidopsis thaliana* were assayed with IGP. Indole produced by TSAs alone (black) and indole produced by TSA + TSB-like (blue) was measured using liquid chromatography/time-of-flight mass spectrometry. **B**) *N*-terminal His tagging of *As*TSA and *As*TSB-like still allows TSA-TSB-like complex formation. *E. coli* cultures expressing *N*-terminal His-tagged or untagged TSA and TSB-like were mixed and His-tagged proteins were retrieved trough affinity purification. Untagged TSA or TSB-like co-purified with the corresponding tagged partner (SDS-page) although only one protein was His-tagged (Western-blot). **C**) Table with the size of *As*TSA and *As*TSB-like with N and C-terminal 6xHis tag or untagged.

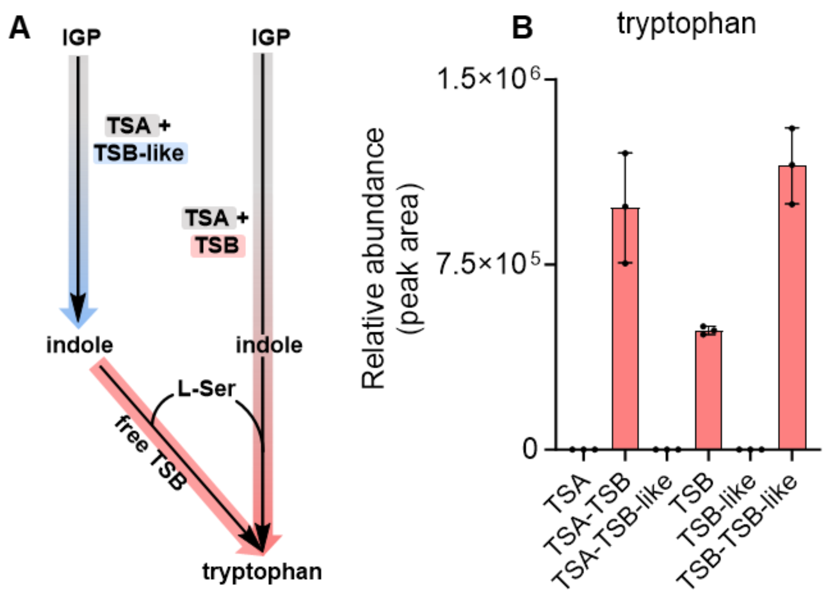


**Fig. S4.**

**Indole can be used as a substrate both from TSA-TSB complex as well as from TSB alone.** **A**) Scheme depicting the cross talk between the product of TSA-TSB-like reaction (indole), which act as a substrate for tryptophan biosynthesis by TSA-TSB or in minor amount for TSB alone. The cross-talk between the product and substrate of the two reactions explains the tryptophan increase concomitant with indole increase in Fig 2D iii and iv. **B**) Both, the TSA-TSB complex as well as TSB alone synthesize tryptophan from free indole and L-Ser. Co-incubation of TSB and TSB-like in the presence of indole and L-serine did not result in altered tryptophan formation compared to TSB alone. Proteins were expressed in *E. coli*, purified, and assayed on indole and L-Ser. Reaction products were analyzed using liquid chromatography/time-of-flight mass spectrometry. Means (n = 3) and SD are displayed.

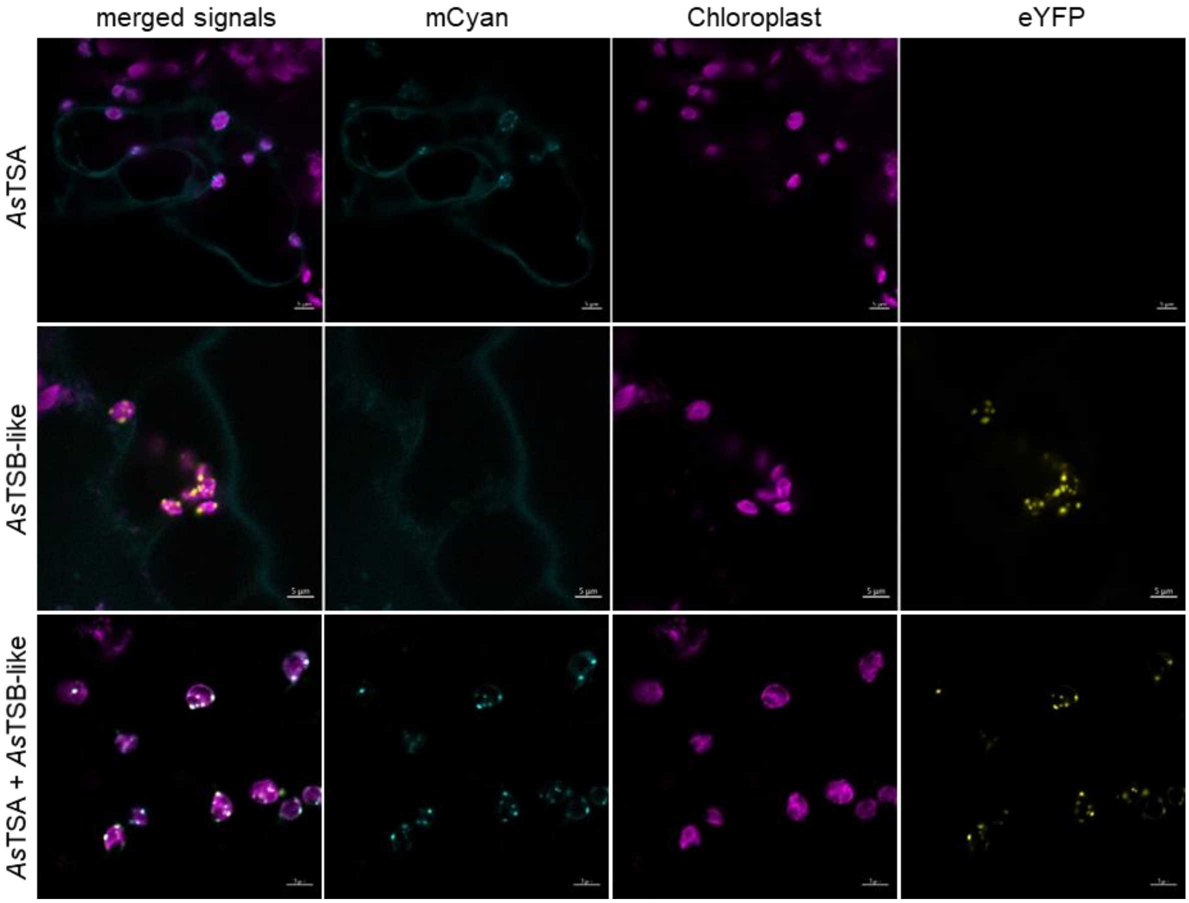


**Fig. S5.**

**Subcellular localization of *As*TSA and *As*TSB-like.** *As*TSA fused to a *C*-terminal mCyan and *As*TSB-like fused to a C-terminal eYFP were transiently expressed in *Nicotiana benthamiana* under control of the *Solanum lycopersicum* Ubq10 promoter and terminator. Chloroplast localization was assessed using chloroplast autofluorescence. *N. benthamiana* was transiently transformed with the single *AsTSA* or *AsTSB-like* construct or co-infiltrated with both.

A

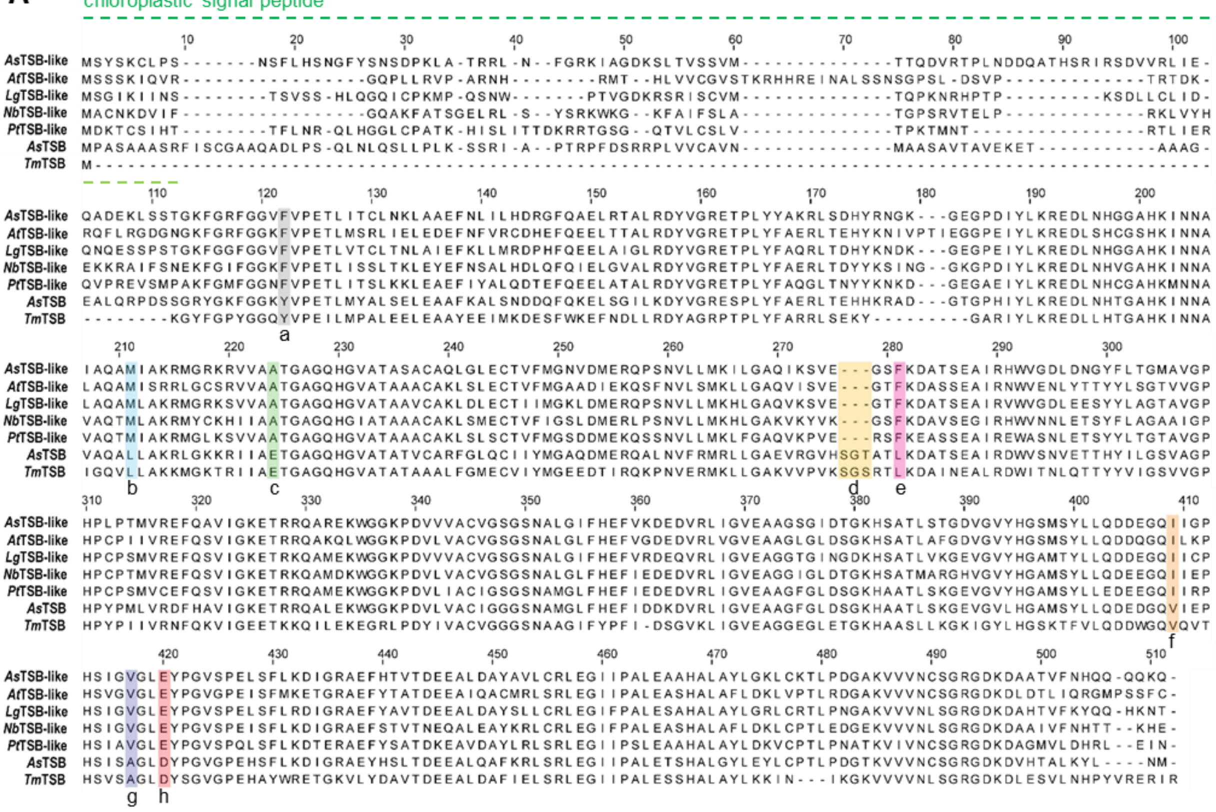


B

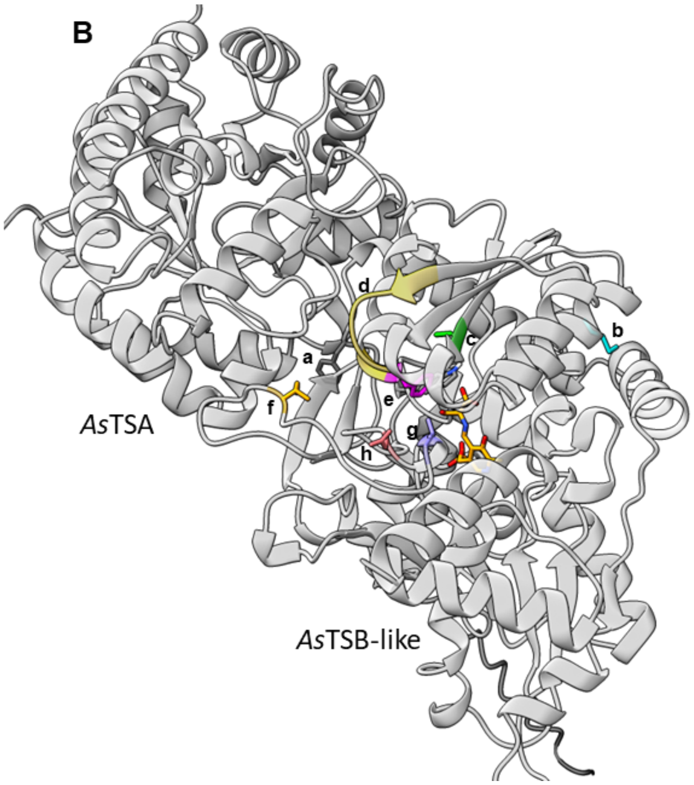


C

| Residue | Location |
| --- | --- |
| a | Near active site |
| b | Near active site |
| c | Active site |
| d | Near active site |
| e | Active site |
| f | Near active site |
| g | Active site |
| h | Active site |

**Fig. S6.**

**Highly conserved residues that differ between TSB and TSB-like. A**) Muscle alignment displaying *Aphelandra squarrosa* (*As*), *Arabidopsis thaliana* (*At*), *Lamium galeobdolon* (*Lg*), *Populus trichocarpa* (*Pt*), *T. maritima* (*Tm*) TSB-like or TSB. **B**) Residues highlighted in (A) are shown in a model of *As*TSA-*As*TSB-like complex. The complex was modelled as a multimer using Alphafold MMSeq. **C**) Table indicating the location of the residues selected for mutagenesis.

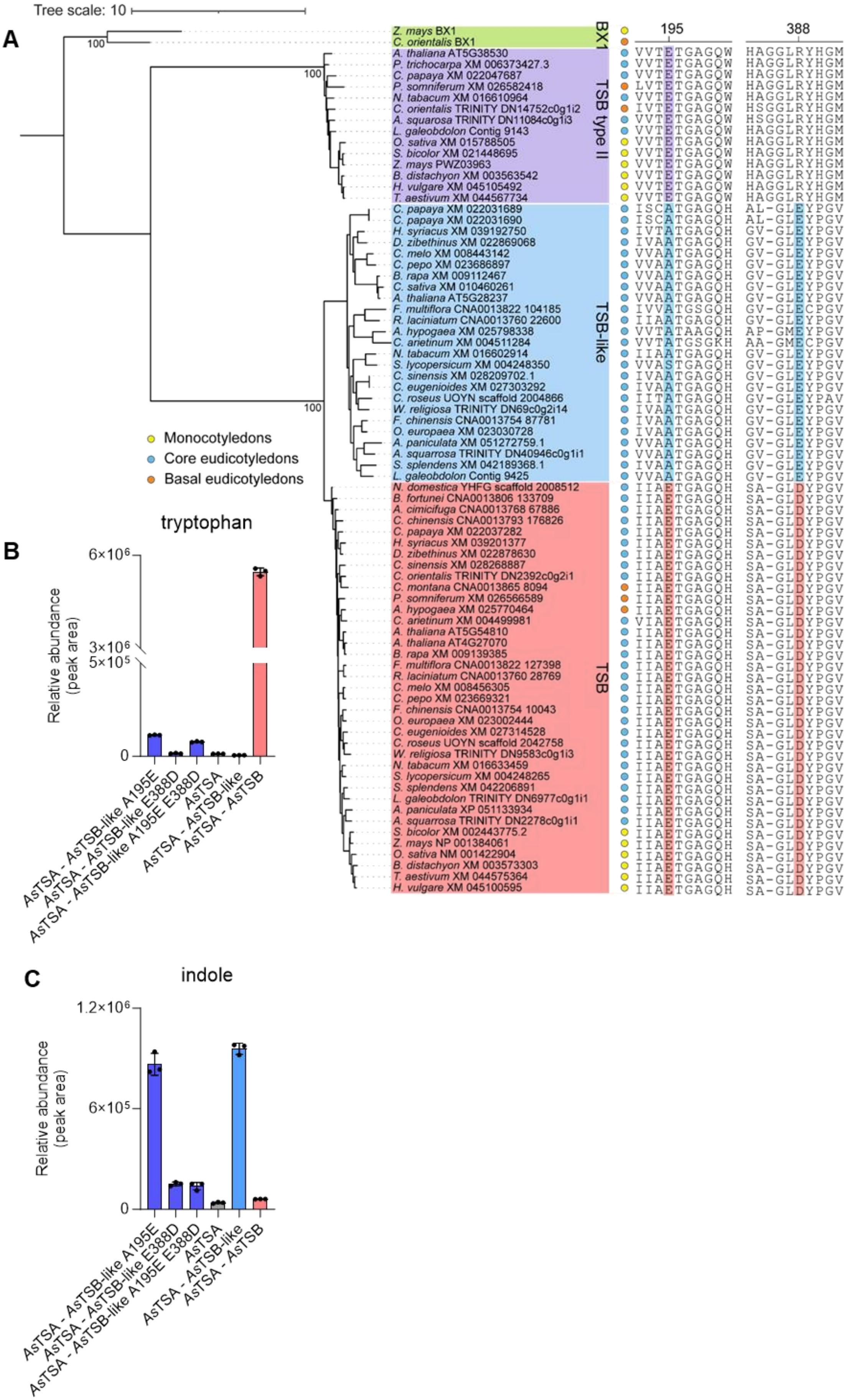

Tree scale: 10
A
100
100
100
BX1
TSB type II
TSB-like
TSB
195
388
Z. mays BX1
C. orientalis BX1
A. thaliana AT5G38530
P. trichocarpa XM 006373427.3
C. papaya XM 022047687
P. somniferum XM 026582418
N. tabacum XM 016610964
C. orientalis TRINITY DN14752c0g1i2
A. squarosa TRINITY DN11084c0g1i3
L. galeobdolon Contig 9143
O. sativa XM 015788505
S. bicolor XM 021448695
Z. mays PWZ03963
B. distachyon XM 003563542
H. vulgare XM 045105492
T. aestivum XM 044567734
C. papaya XM 022031689
C. papaya XM 022031690
H. syriacus XM 039192750
D. zibethinus XM 022869068
C. melo XM 008443142
C. pepo XM 023686897
B. rapa XM 009112467
C. sativa XM 010460261
A. thaliana AT5G28237
F. multiflora CNA0013822 104185
R. laciniatum CNA0013760 22600
A. hypogaea XM 025798338
C. arietinum XM 004511284
N. tabacum XM 016602914
S. lycopersicum XM 004248350
C. sinensis XM 028209702.1
C. eugenioides XM 027303292
C. roseus UOYN scaffold 2004866
W. religiosa TRINITY DN69c0g2i14
F. chinensis CNA0013754 87781
O. europaea XM 023030728
A. paniculata XM 051272759.1
A. squarrosa TRINITY DN40946c0g1i1
S. splendens XM 042189368.1
L. galeobdolon Contig 9425
N. domestica YHFG scaffold 2008512
B. fortunei CNA0013806 133709
A. cimicifuga CNA0013768 67886
C. chinensis CNA0013793 176826
C. papaya XM 022037282
H. syriacus XM 039201377
D. zibethinus XM 022878630
C. sinensis XM 028268887
C. orientalis TRINITY DN2392c0g2i1
C. montana CNA0013865 8094
P. somniferum XM 026566589
A. hypogaea XM 025770464
C. arietinum XM 004499981
A. thaliana AT5G54810
A. thaliana AT4G27070
B. rapa XM 009139385
F. multiflora CNA0013822 127398
R. laciniatum CNA0013760 28769
C. melo XM 008456305
C. pepo XM 023669321
F. chinensis CNA0013754 10043
O. europaea XM 023002444
C. eugenioides XM 027314528
C. roseus UOYN scaffold 2042758
W. religiosa TRINITY DN9583c0g1i3
N. tabacum XM 016633459
S. lycopersicum XM 004248265
S. splendens XM 042206891
L. galeobdolon TRINITY DN6977c0g1i1
A. paniculata XP 051133934
A. squarrosa TRINITY DN2278c0g1i1
S. bicolor XM 002443775.2
Z. mays NP 001384061
O. sativa NM 001422904
B. distachyon XM 003573303
T. aestivum XM 044575364
H. vulgare XM 045100595
Monocotyledons
Core eudicotyledons
Basal eudicotyledons
B
tryptophan
Relative abundance (peak area)
6×10^6
3×10^6
5×10^5
0
AsTSA - AsTSB-like A195E
AsTSA - AsTSB-like E388D
AsTSA - AsTSB-like A195E E388D
AsTSA
AsTSA - AsTSB-like
AsTSA - AsTSB
C
indole
Relative abundance (peak area)
1.2×10^6
6×10^5
0
AsTSA - AsTSB-like A195E
AsTSA - AsTSB-like E388D
AsTSA - AsTSB-like A195E E388D
AsTSA
AsTSA - AsTSB-like
AsTSA - AsTSB

**Fig. S7.**

**Mutation of two conserved residues in TSB-like and TSB leads to activity changes in vitro.** **A**) Phylogenetic tree displaying conservation of residues 195 and 388 among different classes of TSBs. Residues are highlighted with colors according to the TSB clade they belong to. *Solanum lycopersicum* displays a serine instead of the otherwise conserved alanine 195. Amino acid sequences were aligned with WebPrank algorithm and a Maximum Likelihood tree was inferred using iQTree. **B** and **C**) Activity of *As*TSB-like mutants. Tryptophan (**B**) and indole (**C**) biosynthetic activity of *As*TSB-like mutants. Proteins were expressed in *E. coli*, purified, and assayed with IGP and L-serine. Reaction products were analyzed using liquid chromatography/time-of-flight mass spectrometry. Means (n = 3) and SD are displayed.

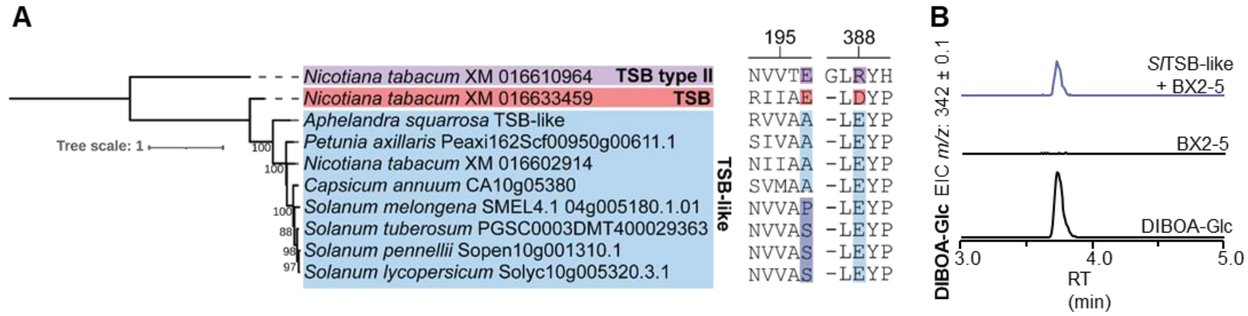


**Fig. S8.**

**A195 is not strictly required for TSB-like activity. A**) Phylogenetic tree displaying the conservation of residue 195 and 388 among the Solanaceae. *As*TSB-like is used as a reference sequence to mark the TSB-like clade. While most TSB-like proteins have an alanine at position 195, members of the *Solanum* genus have either a proline or serine residue at this position. E388 is conserved in all TSB-like. The tree was constructed using amino acid sequences, alignment was performed with WebPrank, and a Maximum Likelihood tree was inferred using iQtree. The tree was midpoint rooted. Sequences used are reported in Data S3. **B**) *S. lycopersicum TSB-like* was expressed in *N. benthamiana* with *ZmBx2-5*. Leaf methanolic extracts were analyzed using liquid chromatography/time-of-flight mass spectrometry and DIBOA-Glc accumulation was confirmed with an authentic standard.

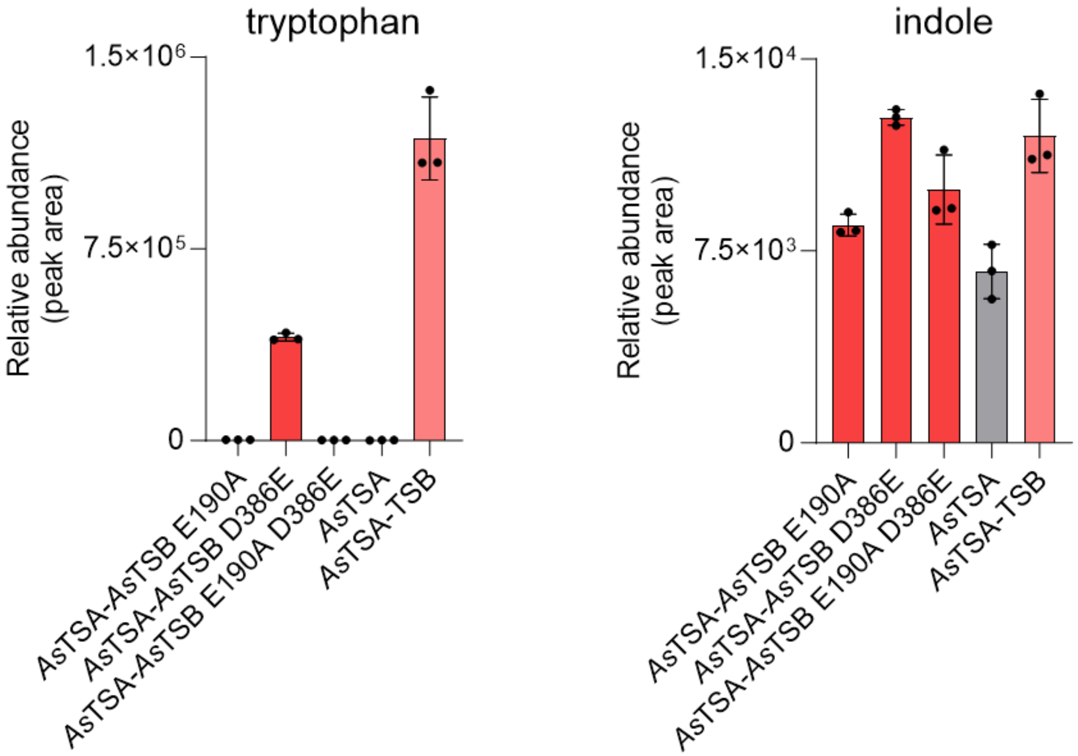


**Fig. S9.**

**The mutation of E190 into alanine and D386 into glutamate is not sufficient to convert *As*TSB into *As*TSB-like.** *As*TSB E190A showed a substantial loss of tryptophan biosynthetic activity, while *As*TSB D386E showed reduced but still significant tryptophan biosynthesis without a significant gain in indole formation. The double mutant *As*TSB E190A D386E did not show tryptophan biosynthetic activity, but this mutant showed no gain of indole biosynthetic activity. Proteins were expressed in *Escherichia coli*, purified, and assayed with IGP and L-serine. Reaction products were analyzed using liquid chromatograpy/time-of-flight mass spectrometry. Means (n = 3) and SD are displayed.

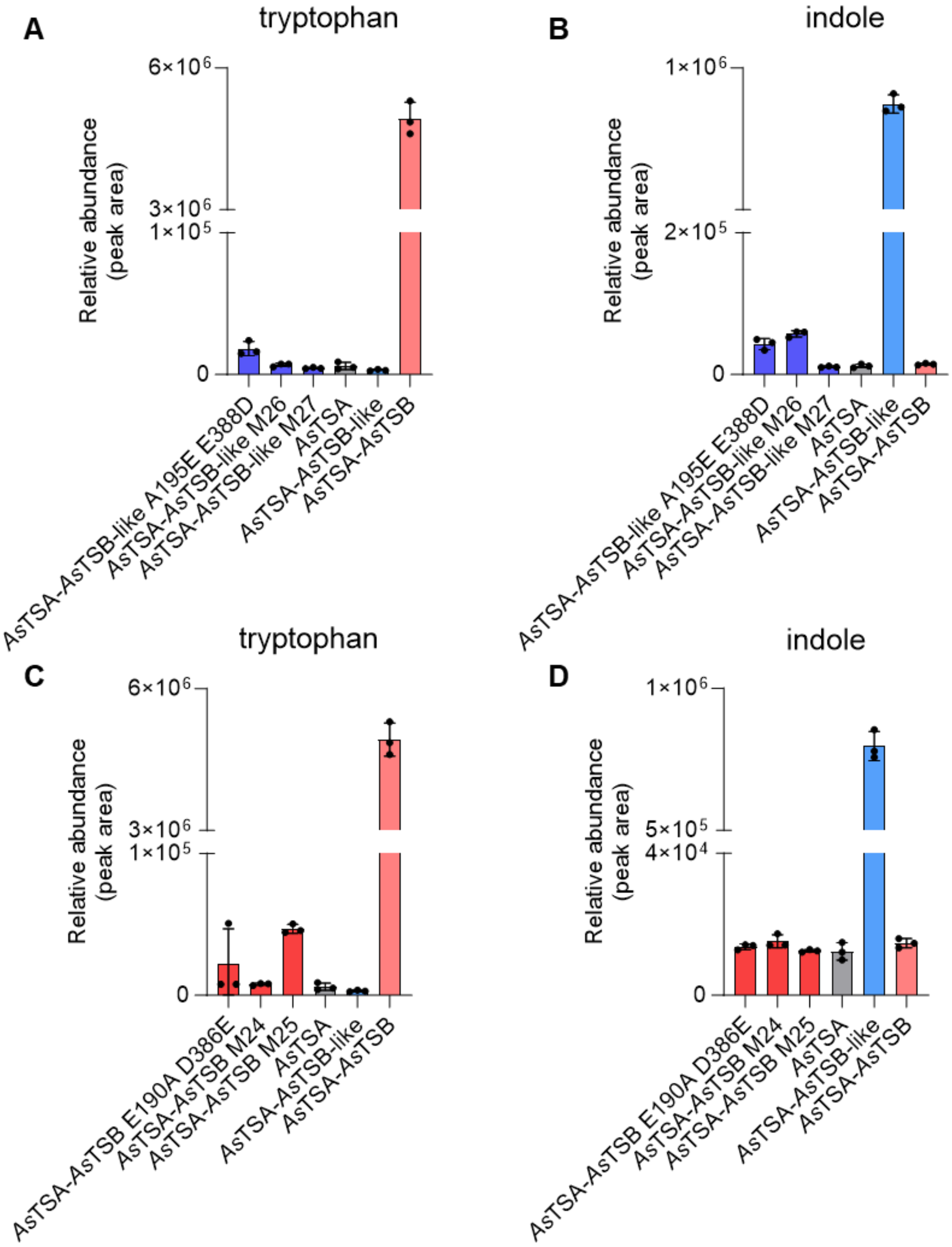


**Fig. S10.**

**Mutagenesis of highly conserved active site and near-active site residues that differ between TSB and TSB-like clades did not substantially change the activity of the resulting proteins.** **A** and **B**) Extended mutagenesis on *As*TSB-like background did not improve tryptophan biosynthetic activity. TSB-like M26 consisted of swapping positions c, e, g, h and M27 positions b, c, d, e, f, g, h (SI Fig. 6) from TSB-like to TSB residues. Proteins were expressed in *Escherichia coli*, purified, and tested in combination with *As*TSA on IGP and L-serine. Reaction products were analyzed using liquid chromatography/time-of-flight mass spectrometry (LC-qTOF-MS). Means (n = 3) and SD are displayed. *As*TSB-like M26 and *As*TSB-like M27 did not show improved tryptophan biosynthesis compared to the double mutant *As*TSB-like A195E E388D. **C** and **D**) Extended mutagenesis on *As*TSB background did not improve indole biosynthetic activity. TSB M24 consisted of swapping positions c, e, g, h and M25 positions b, c, d, e, f, g, h (SI Fig. 6) from TSB to TSB-like residues. Proteins were expressed in *E. coli*, purified, and tested in combination with *As*TSA on IGP and L-serine. Reaction products were analyzed using LC-qTOF-MS. Means (n = 3) and SD are displayed. *As*TSB M24 and *As*TSB M25 did not show improved indole biosynthesis compared to the double mutant *As*TSB E190A D386E.

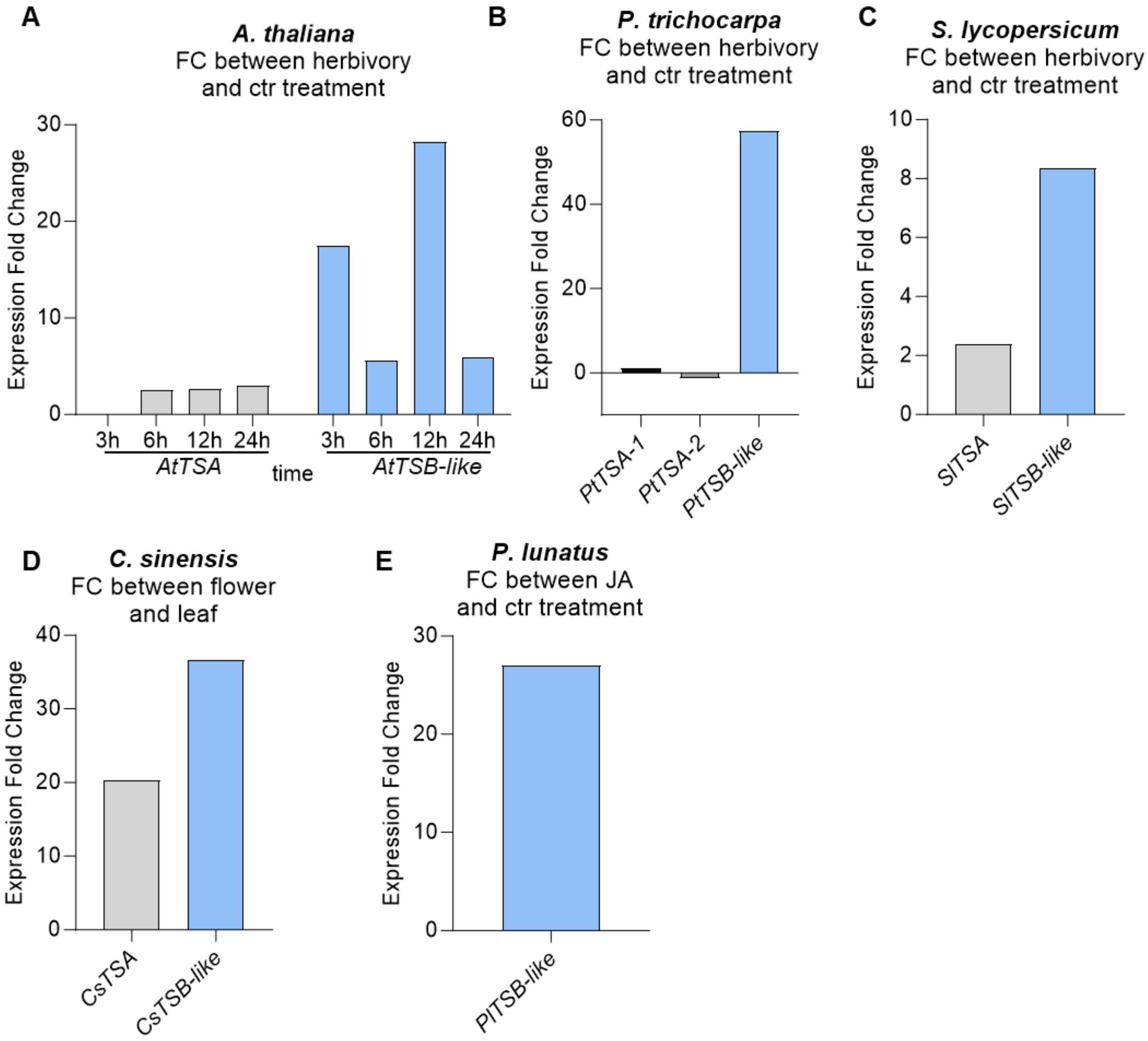


**Fig. S11.**

**Expression differences of *TSA* and *TSB-like* genes between conditions in which indole emission is induced and the control treatment in different plant species.** Expression differences are showed as Fold Change (FC). **A**) FC of *Arabidopsis thaliana TSA* and *TSB-like* over the course of 24 h exposure to *Pieris rapae* caterpillars as reported by *(41, 42)*. **B**) FC of *Populus trichocarpa TSA* and *TSB-like* after 24 h of *Lymantria dispar* caterpillar feeding as reported by *(43)*. **C**) FC of *Solanum lycopersicum TSA* and *TSB-like* after 40 days of *Tuta absoluta* exposure as reported by *(7, 44)*. **D**) FC of *Citrus sinensis* (sweet orange) *TSA* and *TSB-like* between the indole-producing flowers and the indole non-producing leaves (*45*, www.orangeExpDB.com). **E**) FC of *Phaesous lunatus* TSB-like between jasmonic acid (JA) and control treatment. No *TSA* was reported as differentially expressed in *P. lunatus* plants treated with JA as reported by *(46, 47)*.

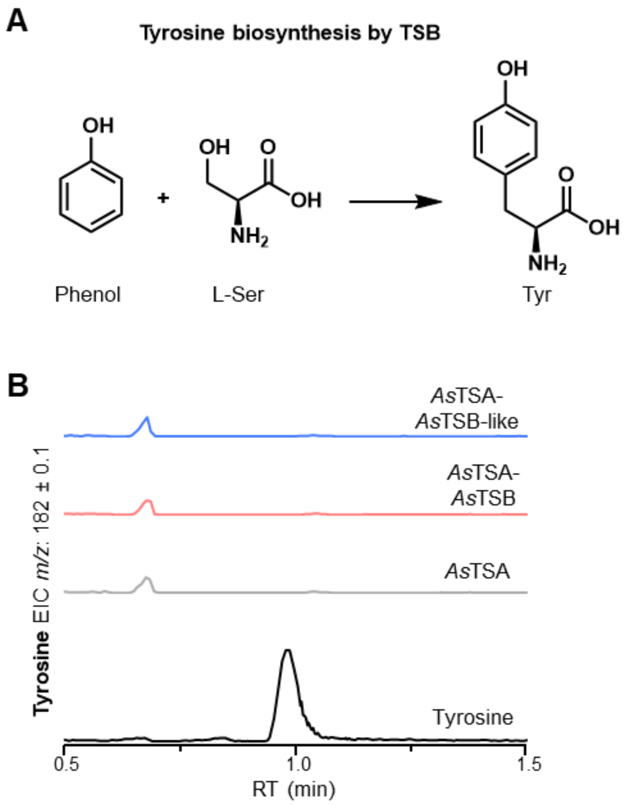


**Fig. S12.**

***As*TSB-like shows no tyrosine biosynthetic activity. A**) Tyrosine formation catalyzed by TSB as proposed by *(30)*. **B**) *As*TSB-like and *As*TSB show no tyrosine biosynthetic activity. Proteins were expressed in *Escherichia coli*, purified, and assayed with phenol and L-serine. Reactions were quenched with MeOH:1MHCl and reaction products were analyzed using liquid chromatography-time-of-flight mass spectrometry. Extracted ion chromatograms (EIC) are shown.

**Table S1. (separate file)**

Meatanalysis of TSB-like expression and indole emission among different species.

**Table S2. (separate file)**

NCBI accessions numbers and nucleotide sequence of the genes characterized in this study.

**Table S3. (separate file)**

Table of primers used in this study.

**Table S4. (separate file)**

qPCR results along with raw data and primer efficiency calculation for *N. benthamiana TSA*, *TSB-like* and housekeeping genes.

**Data S1.**

Sequences used for phylogenetic analysis of Fig S1 A, provided in FASTA format.

```
>ATHALIANAAT3G54640.1
MAIAFKSGVFFLQSPKSQIGFRHSSPPDSSLSFKRFTPMASLSTSSPTLGLADTFTQLKKQGKVAFIPYITAGD
PDLSTTAEALKVLDACGSDIIELGVPYSDPLADGPVIQAAATRSLERGTNLDSILEMLDKVVPQISCPISLFTY
YNPILKRGLGKFMSSIRAVGVQGLVVPDVPLEETEMLRKEALNNDIELVLLTTPTTPTERMKLIVDASEGFI
YLVSSIGVTGARSSVSGKVQSLLKDIKEATDKPVAVGFGISKPEHVKQIAGWGADGVIVGSAMVKLLGDA
KSPTEGLKELEKLTKSLKSALL
>SOLYC01G098550.3.1_SLYCOPERSICUM_ITAG4.0
MSFHSYTTVRPYLTPSPSASPLFLHFQVERRFGDGMASFLKATHFIHSSNNHQNHPFSQSYTQTIKISTSRKPL
MAALSATATVGLSETFTRLKEQGKVALIPYITAGDPDLATTAEALKVLDRCGSDIIELGVPYSDPLADGPVI
QAAASRSLTKGTNFAKVISMLEDVVPQLSCPIALFTYYNPILKRGTEKFMATVRDTGVHGLVVPDVPLEET
EMLRKEAARHNIELVLLTTPTTPKIRMKAITEASEGFVYLVSAVGVTGARASVSSKVQSLLLDIKEATSKPV
AVGFGISKPEHVKQVAEWGADGVIVGSAMVRILGEAKSPEEGLKELEVFTTSLKSALS
>NBENTHA_NBV6.1TRP68353
MASFLKATQFIHSTNKLENHAFPHSYNNTQRLKISTSVKPPMAALSTTPTVGLSETFTRLRKQGKVALIPYIT
AGDPDMSTTAEALKVLDRCGSDIIELGVPYSDPLADGPVIQAAATRSLSRGTNFAKVLSMLKDVVPQLSCPI
ALFTYYNPILKRGTEKFMATVRDAGVHGLVVPDVPLEETEILRKEAARHNIELVSLCARYLSQFPTIMRCNK
VLLTTPTTPTIRMKAITEASEGFVYLVSSTGVTGARESVSSKVQPLLIDIKEATPKPVAVGFGISKPEQVKQV
AGWGADGVIVGSAMVRILGEAKSPEEGLKELEVFATSLKSALS
>POTRI.005G217700.1_PTRICHOCARPA_V4.1
MAVALKSTPSFLQLKKPETYFIVRNKPPIVSTRRFAPMASLTATRSLGIGETFSNLKKQGKVALIPYITAGDP
DLSTTAEALKLLDACGCDIIELGVPYSDPLADGPVIQAAATRSLARGTNFEAITSMLKEVVPQVSCPIALFTY
YNPILKRGIEKFMSTVNDIGVHGLVVPDVPLEETQVLRKEAVKNGLELVLLTTPTTPTERMKAIVEAADGF
VYLVSSVGVTGTRASVSDRVQTLLQDIKETTTKPVAVGFGISKPEHVKQVAGWGADGVIVGSAMVKLLGE
AKSPEEGLKELESFTKSLKAALP
>ORANGE1.1G021527M_CSINENSIS_V1.1
MAALQATTNFVHLKNPHAHYLPRLPCHKSTLSLKRFTPMAALTASPTVGLAETFTRLKKQGKVALIPYITA
GDPDLSTTAEALKLLDSCGSDIIELGVPYSDPLADGPVIQAAATRSLARGTNFNAILSMLKEVVPQMSCPIAL
FTYYNPILKRGVDNFMSTVRDIGIRGLVVPDVPLEETESLQKEAMKNKIELVLFTTPTTPTDRMKAIVEASE
GFVYLVSSIGVTGARASISGHVQTLLREIKESSTKPVAVGFGISKPEHVQQVAGWGADGVIVGSAMVKLLG
EAQSPEEGLKELEKFAKSLKSALP
>MEDTR7G102980.1_MTRUNCATULA_MT4.0V1
MAIAFKSSCFLQFNKPNTGFGLSFSSRKPVIISVKRYTSVAAIKTMETVGISETFNRLKKQGKVALIPFITAGD
PDLSTTAEALKVLDSCGSDIIELGVPYSDPLADGPVIQAAATRALARGSNFDSIISMLNEVIPQISTPIALFTYY
NPILKRGTGKFMSIVRDTGVHGLVVPDVPLEETKTLREEAKKHGIELVLLTTPTTPTDRMKAIVDAAEGFVY
```

LVSSVGVTGARASVSGKVQALLQDIKEATTKPVAVGFGISTPEHVKQIVGWGADGVIVGSAMVRLLGEAK
SPEEGLKELEKFTRSLKSALD
>AT4G02610.1_ATHALIANA_TAIR10
MDLLKTPSSTVGLSETFARLKSQGKVALIPYITAGDPDLSTTAKALKVLDSCGSDIIELGVPYSDPLADGPAI
QAAARRSLLKGTNFNSIISMLKEVIPQLSCPIALFTYYNPILRRGVENYMTVIKNAGVHGLLVPDVPLEETET
LRNEARKHQIELVLLTTPTTPKERMNAIVEASEGFIYLVSSVGVTGTRESVNEKVQSLLQQIKEATSKPVAV
GFGISKPEHVKQVAEWGADGVIVGSAMVKILGESESPEQGLKELEFFTKSLKSALVS
>CSACSN226.13G304300.1_CSATIVA_ACSN-226_V1.1
MDSLKNPPTTVGLSETFARLKTQGKVALIPYITAGDPDLSTTEKALKVLDSCGSDIIELGVPYSDPLADGPAI
QAAARRSLLKGTNFNSIITMLKEVIPLLSCPIALFTYYNPILRRGIENYMTIIKDAGVHGLLVPDVPLEETETL
RIEARKQQIELVLLTTPTTPKERMSAIVEASEGFIYLVSSVGVTGTRESVNEKVQSLLQQIKEATSKPVAVGF
GISKPEHVKQVAEWGADGVIVGSAMVRILGESESPERGLKELEVFTKSLKSALVS
>CSACSN226.09G173100.1_CSATIVA_ACSN-226_V1.1
MAIAFKSGVFFLQSPKTQFGFRHSSSSPDSSLSFKRLTPMASLSTSSPTLGLADTFTQLKKQGKVAFIPYITAG
DPDLSTTAEALKVLDACGSDIIELGVPYSDPLADGPVIQAAATRSLEKGTNLDNIFEMLDKVVPQISCPISLFT
YYNPILKRGLGKFMSSIRAVGVQGLVVPDVPLEETEMLRKEALANDIELVLLTTPTTPTERMKRIVDASEGF
IYLVSSIGVTGARASVSGKVQSLLKDIKEATDKPVAVGFGISKPEHVKQIVEWGADGVIVGSAMVKLLGDA
NSPTEGLRELEKLTKSLKSALL
>CARUB.0006S3453.1_CRUBELLA_V1.1
MDLLQNPPATTTVGLSETFARLKTQGKVALIPYITAGDPDLSTTAKALKVLDSCGSDIIELGVPYSDPLADG
PAIQAAARRSLLKGTNFNSIITMLKEVIPQLSCPIALFTYYNPILRRGVENYMTIIKEAGVHGLLVPDVPLEET
ETLRSEARKQQIELVLLTTPTTPKHRMTAIVEASEGFIYLVSSVGVTGTRESVNEQVQSLLQQIKEATSKPVA
VGFGISKPEHVKQVAEWGADGVIVGSAMVKILGESESSEKGLKELEVFTKSLKSALVS
>CARUB.0005S2124.1_CRUBELLA_V1.1
MAIAFKSGVFFLQSPKTQFGFRHSSTSPDASLSFKRLTPMASLSTSSPTLGLADTFTQLKKQGKVAFIPYITA
GDPDLSTTAEALKVLDACGSDIIELGVPYSDPLADGPVIQAAATRSLEKGTNLDNILEMLDKVVPQISCPISL
FTYYNPILKRGLGKFMSSIRAVGVQGLVVPDVPLEETEMLRKEALNNDIELVLLTTPTTPTERMKRIVDASE
GFIYLVSSIGVTGARTSVSGKVQSLLKDIKEATDKPVAVGFGISKPEHVKQIAGWGADGVIVGSAMVKLLG
DAKSPTEGLKELEKLTKSLKSALL
>Z.MAYSIGL
MASAIKAASTSSRWSSSPAAVHSSPLSKRLPAAVAMPGRRRSVATVRAVAAVAPAAPAAPAKLTAGAGGR
CLPVSQTMSRLRAQGKTAFIPYITAGDPDLATTAEALRLLDACGADVIELGVPFSDPYADGPVIQASAARAL
ASGTTPDGVLAMLKEVTPELSCPVVLFSYFNPIVRWGLADFAAAVKEAGVHGLIVPDLPYGNSCALTLRTE
AIKNSLELVLLTTPSTPADRMEEITRASRGFVYLATVNGVTGPRANVNTRVQSLIQEVKQVTDIPVAVGFGI
SKPEHVKQIAEWGADGVIIGSAMVRQLGEAASPKEGLKRLEKYARSMKNALPCQ
>Z.MAYSTSA-LIKE
MANGGAAAGKLTVAETFSNLREQGKSAFIPFITAGDPDLVTTSKALKILNSCGSDVIEVGVPYSDPLADGPV
IQASATRALKKGTTLDSVIEMLKGVTPELSCPIVLFTYYNPILKRGVGNFMSTIKQAGIHGLVVPDLPLEETT
LLRSEAIMHNIELVLLTTPTTPTDRMKGIAQASEGFLYLVSAVGVTGARSNVNLRVEHLLREIKKVTDKPVA
VGFGVSTPEHVKQIVGWGADGVIVGSAIVKQLCEAATPEEGLERLEEYARSMKAAMP
>Z.MAYSBX1
MAFAPKTSSSSSLSSALQAAQSPPLLLRRMSSTATPRRRYDAAVVVTTTTTARAAAAAVTVPAAPPQAPAP
APVPPKQAAAPAERRSRPVSDTMAALMAKGKTAFIPYITAGDPDLATTAEALRLLDGCGADVIELGVPCSD
PYIDGPIIQASVARALASGTTMDAVLEMLREVTPELSCPVVLLSYYKPIMSRSLAEMKEAGVHGLIVPDLPY
VAAHSLWSEAKNNNLELVLLTTPAIPEDRMKEITKASEGFVYLVSVNGVTGPRANVNPRVESLIQEVKKVT
NKPVAVGFGISKPEHVKQIAQWGADGVIIGSAMVRQLGEAASPKQGLRRLEEYARGMKNALP
>ASQUARROSA_TSA
MAAAALKASCFVQPKASFDTRGRRRSLLAVPTSTSFKCKPPMAALTTAPTLSISETFSKLKQRGEVALIPYIT
AGDPDLSTTAKALKILDSSGADIIELGVPYSDPLADGPVIQAAATRALARGASFEKVIGMLKDVVPQLSSPIA
LFIYYNVILKRGVKKFVTTLKETGVHGIIVPDVPLEETELLRNEAVKYNIEMVLLTTPTTPTERMKAIAEVAQ
GFIYLVSSVGVTGARASVSDKVPSLLHEIREVTNKPVAVGFGISKPEHVKQVAEWGADGVIIGSAMVKVLG
EANSPEEGLKDLEAFMKSLKAAATRGDSVLL
>POTRI.002G045700.1_PTRICHOCARPA_V4.1
MAAALKSTPSFLQLKKPETHFLVRHKPTIVSTRRFAPMASLTAIRSLGIGETFSNLKKQGKVALIPYITAGDP
DLKTTAEALKVLDACGCDIIELGVPYSDPLADGPVIQAAATRSLARGTNFEAITSMLREVVPQVSCPIALFTY
YNPILKRGIEKFMSTVKDIGVHGLVVPDVPLEETGVLRKEAVKNKLELVLLTTPTTPTERMKAIVEAADGF

VYLVSSVGVTGARASVSDRVQTLLRDIKESTTKPVAVGFGISKPEHVKQVAAWGADGVIVGSAMVKLLGE
AKSPEEGLKELESFTKSLKAALP
>ZMTSA_GRMZM5G841619
MAFALKAAAAGSASFSAAGPRRRAAATGRVSFRSAAPVVAVRAAAAAAAAVAEDKRSISGTFAELRQQG
KTALIPFITAGDPDLATTAKALRILDACGSDVIELGVPYSDPLADGPVIQASATRALAKGTTFEDVISMVKGV
IPDLSCPVALFTYYNPILKRGVPNFMSIVKEAGVHGLVVPDVPLEETDVLRSEAAKNNLELVLLTTPTTPNE
RMEKIAQASEGFIYLVSTVGVTGTRANVSGKVQSLLQDIKKVTEKPVAVGFGVSTPEHVRQIAGWGADGVI
IGSAVMKTLEEAASPEEGLKKLEEFAKNLKAALP
>SOBIC.002G054700.1_SBICOLOR_V3.1.1
MAFALKAAATGSASFSAAGPRRGAASATGRVSFRGAAPAVAVRAAAAAAAAVAEDKRSISGTFAELREQ
GKTAFVPFITAGDPDLSTTAKALKILDACGSDVIELGVPYSDPLADGPVIQASATRALAKGTTFEDVISMVK
GVIPELSCPVALFTYYNPILKRGIPKFMSIVKEAGVHGLVVPDVPLEETDVLRSEAAKNNLELVLLTTPTTPN
ERMEKIAQASEGFIYLVSTVGVTGTRANVSGKVQSLLQDIKKVTEKPVAVGFGVSTPEHVQQIAGWGADG
VIVGSAMVRLLGEAASPEEGLKKLEELAKNLKAALP
>LOC_OS07G08430.1_OSATIVA_V7.0
MAFALKASTASAAAASTASASASSLSVAAAAPGRRGGAAGRVSFRGVPAPMVAIRAEAAAVGEDERVISG
TFAKLKEQGKTAFIPFITAGDPDLATTAKALKILDACGSDLIELGVPYSDPLADGPVIQASATRALSKGTTFE
DVISMVKEVIPELSCPVALFTYYNPILKRGIANFMTVVKEAGVHGLVVPDVPLEETNILRSEAAKNNLELVL
LTTPTTPTERMEKITKASEGFIYLVSTVGVTGARANVSGKVQSLLQDIKQVTDKAVAVGFGISTPEHVKQIA
GWGADGVIIGSAMVRQLGEAASPEEGLKKLEELAKSLKAALP
>PAVIR.2NG061900.1_PVIRGATUM_V5.1
MAFALKAASTAGSASLAAAGPRRAAAPAAGRVSFRGASAAGPVVAVRAAAAAASAVAHDRRSISGTFAE
LREQGKTALIPFITAGDPDLATTAKALKILDACGSDVIELGVPYSDPLADGPVIQASATRALAKGTTFEDVIS
MVKEVIPELSCPVSLFTYYNPILKRGIPNFMTIVKEAGLRGLVVPDVPLEETDFLRSEAAKNNLELVLLTTPT
TPNERMEKIAQASEGFIYLVSTVGVTGTRANVSGKVQSLLQDIKKVTEKPVAVGFGVSTAEHVKQIAGWG
ADGVIVGSAMVRLLGESASPEEGLKKLEELAKNLKAALE
>PAVIR.9NG392800.1_PVIRGATUM_V4.1
MADHGVVAGKRTVAGTFSRLREQVKTAFIPFITAGDPDLATTSKALKILDSCGSDVIEVGVPYSDPLADGPV
IQASATRALKKGTTLESVIGMLKGVAPELSCPIVLFTYYNPILKRGVRNFMATIRQAGINGLVVPDLPLEETV
LLRSEAIMHSIELVLLTTPTTPTERMIEIVKASEGFLYLVSAVGVTGARSNVNLRVEHLLREIKKVTDKPVAV
GFGVSTPEHVKQIAGWGADGVIIGSAIVRQLCEAATPEEGLKRLEEYTRNITAAMPLR
>LOC_OS03G58260.2_OSATIVA_V7.0
MEMEDSGRGVVGAGKRGVAETFSRLREQGKTAFIPFITASDPDLATTSKALKILDSCGSDVIELGVPYSDPL
ADGPVIQAAATRALKKGATFDSVIAMLKGVIPELSCPIVIFTYYNPILKRGVSNFMAIIKQAGVHGLVVPDLP
LEETALLRNEAVMHGIELVLLTTPTTPTERMKEIAKASEGFIYLVSSVGVTGARSNVNLRVEYLLQEIKKVT
DKPVAVGFGISTPEHVKQIAGWGADGVIIGSAIVRQLGEAASPEEGLKRVEEYAKNMKAAMP
>SOBIC.001G056500.1_SBICOLOR_V3.1.1
MASAIKAASTSRWSSSPAVQSSPLPKRVAMPGRRRSVATVRSVAAVAPAAPAPARLTGAGLTVSQTMSKL
RAQGKTAFIPYITAGDPDLATTAEALRLLDACGADVIELGVPFSDPYADGPVIQASAARALASGTTPDAVLA
MLQEVTPELSCPVVLFSYFNPIVHWGLPDFAAAVKDAGVHGLIVPDLPYGASCALRTEAIKNNIELVLLTTP
STPADRMEEITKASQGFVYLVSVNGVTGPRANVNTRVESLIQEVKQVTDKPVAVGFGISKPEHVKQIAEWG
ADGVIIGSAMVRQLGEAASPKEGLKRLEKYARSMKNALPCQ
>SOBIC.001G056400.1_SBICOLOR_V3.1.1
MALFAVQAASTSQSSSSSPAVLQQSPLPSRRAAAAAVKKMPQRKKAAAVVRAVAAPPPAPPVPGPPKPAG
ERCRLPVSQTMSRLKAQGKTAFIPYITAGDPDLATTAEALRLLDACGADVIELGVPFSDPYADGPVIQASMA
RARTTGGGATPDGVLAMLREVTPELSCPVVLFSYFNPIVRWGLPGFAAAVKDAGAHGLIVPDLPYADTCA
LRSEAIKNDLELVLLTTPATPEERMKEITEASEGFVYLVSVNGVTGSRADVNTRVESLIQEVKQVTDKPVAV
GFGISKPEHVKQIAEWGADGVIIGSAMVKQLGEATSPKEGLKRLKEYARSMKNALQ
>LOC_OS03G58290.1_OSATIVA_V7.0
MAFTTMKASPMSASSSSAPVLRRCVAQPARVAAARRLAAAAASVALEASPVPAAAAAAVERRMSVSQTM
SKLKEKGKTAFIPYITAGDPDMGTTAEALRLLDACGADVIELGVPFSDPYADGPVIQASASRALAAGATPEA
VLSMLKEVTPELSCPVVLLSYLGPILRRGAANFTAAAKEAGVQGLIVPDLPYVDTCTFRSEAIKSNLELVLL
TTPATPGERMKIITEASGGFVYLVSVNGVTGPRPKVNTRVEHLLQDIKLVTDKAVCVGFGISTPDHVRQIAG
WGADGVIIGSAMVRQLGEAASPKQGLKRLEEYARRMKDALP
>LOC_OS03G58320.1_OSATIVA_V7.0

MAFTVKASSPSSPATTRLTGALHGGAARVAARKLPAAAVAASLTLDRAPAPAGAERGMSSSVSRTMSRLR
EKGKAAFIPYITAGDPDMGTTAEALRVLDACGADVIELGVPFSDPYTDGPVIQASAARALAAGATMDGVM
SMLAEVTPELSCPVVLFSYFGPIVRRGPANFTAAAKEAGVQGLIVPDLPYVETSTFRSEAIKNNLELVLLTTP
ATPADRMKAITAASGGFVYLVSVNGVTGSIQNVNPRVEHLLQEIKQVTDKAVCVGFGISTPDHVRQIADW
GADGVIIGSAMVRQLGEAASPKQGLKRLEKYARSLKDALP
>LOC_OS03G58300.1_OSATIVA_V7.0
MAFTVKASSPSSPAASSSSSSAPAKLGAAPGRVAVRKLTAAATSLRLDRAPAAPATERGLSSVSRTMSRLM
EKGKTAFIPYITAGDPDMGTTAEALRLLDACGADVIELGVPFSDPYNDGPVIQASAARALSAGATMDGIMS
MLAEVTPELSCPVVLFSYLGPIVRRGPANFTAAAKEAGVQGLIVPDLPYLEACSFRSEVIKNNLELVLLTTPT
TPPDRMKAITAASGGFVYLVSVNGVTGSRQDVNPRVEHLLQEIKQVTDKAVCVGYGISTPDHVRQIAEWG
ADGVIIGSAMVRQLGEAASPKQGLKRLEKYARSLKNALP
>PAVIR.9NG392900.1_PVIRGATUM_V4.1
MAFAIKAASTSHLTFFPAVHRPSSSLTTPAASVTKIPAGRKKSAAVTRAVAAVAPPPAPAPARPAGKRCLSV
SQTMSRLKAQGKTAFIPYITAGDPDLATTAEALRLLDACGADVIELGMPFSDPYADGPVIQASAARALASG
TTTDGLLAMLKEVTPELSCPVVLFSYFNPIVRWGLADFAAAAKDAGVEGLIIPDLPYAATCTLRSEAMKNK
LELVLLTTPATPEERMKEITRASEGFIYLVSVNGVTGPRANVSTRVESLIQEVKKVTNKPVAVGFGISKPEHV
KQAREIAKWGADGVIIGSAMVKQLGEAASPKEGLQRLEDYARSMKNALP
>GMAXGLYMA_03G158400.1
MALALKSSCFLQLKKPEAGFNVCFSSKKAIISVKRHTPVAAIRTMEAVGLSATFTRLKKEGKVAFIPYVTAG
DPDLSTTAEALKVLDSCGSDIIELGVPYSDPLADGPVIQAAATRSLAKGTNLNAIIDMLKEVVPQLSCPIALF
TYYNPILKRGTDKFMSTIRDSGVHGLVVPDVPLEETETLRTEAKKHGIELVLLTTPTTPTNRMRAIVDVAEG
FVYLVSSVGVTGARASVSGSVQSLLKEIKEATTKPVAVGFGISKPEHVKQVVVWGADGVIVGSAIVKVLGE
AKSPQEGLKELEVFTRSLKAALP
>MGUTTATUSMGTOL_D0370.1
MAAALKATCFLQLKTSYSLPERRSSTTNTSFKFKPIMASLATAAPTVGLAETFSRLKQQGKVAFIPYITAGD
PNLSTTAEALKVLDLCGSDIIELGMPYSDPLADGPVIQAAATRSLARGTNFDKIIAMLKEVVPELSCPVALFS
YYNPILKRGVENFMTILNDTGVHGLVVPDVPLEETEILRKEAIKKNIELVLLTTPTTPTARMKAIVEVSEGFV
YLVSSIGVTGARASVSEKVQSLLQEIKEASDKPVAVGFGISTPEHVKQVAGWGADGVIVGSAMVKILGDAK
SPEEGLKELEAFTRSLKSALL
>STUBEROSUMSOLTU_DM.01G038030.1
MASFLKATHFIHSNNNHQNHPFSQSYTQRIKISTSRKPLMAALSATATVGLSETFTRLKEQGKVALIPYITAG
DPDLATTAEALKLLDRCGSDIIELGVPYSDPLADGPVIQAAASRALSRGTNFAKVISMLEDVVPQLSCPIALF
TYYNPILKRGIEKFMATVRDTGVHGLVVPDVPLEETEMLRKEAARHNIELVLLTTPTTPKIRMKAITEASEG
FVYLVSAVGVTGARASVSAKVQSLLIDIKEATSKPVAVGFGISKPEHVKQVAEWGADGVIVGSAMVRILGD
AKSPEEGLKELEVFTTSLKSALS
>LGALEOBDOLONEU747715
MASSLKATGFLQLRTNYSLPEYPLYPSSLTSSTNRPFKLRPIMATLTAAPTVGLSKTFSRLKQQGKVAFIPYI
TAGDPDLSTTAEVLKVLDSCGSDIIELGVPYSDPLADGPVIQAAASRALARGTNFDKIIAMLKEVVPQLSCP
VALFTYYNPILKRGAGKFMATLQDTGIHGLVVPDVPLEETEILRKEAISKNIELVLLTTPTTPKARMQSIVEV
SEGFVYLVSSVGVTGARASVSDRVQNLLKEIKEATDKPVAVGFGISKPEHVKQVADWGADGVIVGSAMV
KILAEAKSPEEGIKEIETYTKSLKSALS
>LGALEOBDOLONEU747716
MAANSLKSICFPQLKTTNQSISQRSSSRVSLCYKPPMATLQTATKAGISETFSRLKQQGKVALIPYITAGDPD
LSTTAEALKVLDSCGSDIIELGVPYSDPLADGPVIQAAATRSLSRGADFDKIIGMLKEVIPQLSCPVGLFTYY
NPILKRGVEEFMTTLKDTGINGLIVPDVPLEETEMLRKEATKNNIELVLLTTPTTPSDRMKEIVEAAEGFIYL
VSTVGVTGTRTSVSDKVQSLLHEIKGETNKAVAVGFGISKPEQVKQVAEWGADGVIVGSAMVKILGEAKT
PEEGLKELEKFTKTLKSALV
>C.ORIENTALISBX1
MALAITSSAFSLVCQKPAVIQKSSETRGSLTISPSSLTISPSSVSISETFASLRQQGKVALVPYITAGDPDLSTT
AEALKVLDYCGSDVIELGVPCTDPFLDGPVIQAACKRSLGGGANMKSIFSMLQKVSPQLSCPILLFTYYKQI
LKCGIGRFMAATNDAGVRGLLVPDAPLEHTEVLRAEASKYGIEIVLLTTPITPKERMKKIVQVAQGFVYLVS
SVGVTGARPSVNPRVQSLLQEXKEVTNKPVVVGFGISKPEHVKQIARWGADGVIVGSAMVKLLGEAKTAN
EGLKELEAFTMSLKTALSENNSLLMI
>C.ORIENTALISEU747713
MAAAFKSTCFLQSSNPTNNLFLRSSTQKLKTSNISTKSRISMASLAVAPPTAATAVSLSETFIRLKEQGRVAFI
PFITAGDPDLSTTAEALKVLDSCGSDIIELGVPYSDPLADGPVIQAAATRALARGTNFDAILAMLEGVVPQLS

CPLALFSYYNPILKRGIGKFMTTIKDVGVHGLVVPDVPLEETAMLREEALKNQIELILLTTPTTPTDRMKAIV
EASEGFVYLVSSIGVTGSRASVSSRVQSLLQEIKETSNKPVAVGFGISTPEQVKQIAGWGADGVIVGSAMVE
LLAHAKTPEEGLKELEAFTKSLRAAIP
>PNUDICAULEMCL7050544
MSSSTHLQDSATTVGISQTFTRLRRQGQVAFIPYVTAGDPNLQTTAEALKVLDSSGADIIELGIPFSNPFADG
PVNQAAARRALANGTNFDGIISMLKEVVPQISSPIALFTYYNQILDVGIAEFISAIQSVGVHGLVVPDIPFKDT
AILRREAISLHHKIELVLLTKPDSSTERMKAIVEASQGFVYLVSSMGVTGPRPVVNSQVQDLIREIKKATTKP
VAVGFGLSKPEHVKQVAEWGADGVIVGSALVKLLGEAKSPKEGLKELETFAKSLKAALPLRVPSALPLRV
PSELCAQLLTLAK
>PNUDICAULEMCL7022747
MVCSSTYLQDSSTTVGISQTFTRLRRQGQVAFIPYVTAGDPNLQTTAEALKVLDSSGADIIELGIPFSNPFAD
GPVNQAAARRALANGTNFDGIISMLKEVVPQISSPIALFTYYNQILDVGIAEFISAIQSVGVHGLVVPDIPFKD
TAILRREAISLHHKIEVVLLTKPDSSTERMKAIVEASQGFVYLVSSMGVTGPRPVVNSQVQDLIREIKKATTK
PVAVGFGLSKPEHVKQVAEWGADGVIVGSALVKLLGEAKSPKEGLKELETFAKSLKDALPLRVPSALPLR
VPYELCAQLLTLAK
>PNUDICAULEMCL7037234
MELTNLYYSPRHQNCKNRPLNRRNNQTIQSLVMVCSSTYLQDSSTVSISQTFTRLRRQGQVAFIPYVTAGDP
NLPTTAEALKVLDSSGADIIELGIPFSNPFADGPVNQAAARRALANGTNFDGIISMLKEVVPQISSPIALFTYY
NQILDVGIAEFISAIQSVGVHGLVVPDIPFKDTAILRREAISLHHKIELVLLTKPDSSTERMKAIVEASQGFVY
LVSSMGVTGPRPVVNSQVQDLIREIKKATTKPVAVGFGLSKPEHVKQVAEWGADGVIVGSALVKLLGEAK
SPKEGLKELETFAKSLKDALPLRVPYELCAQLLTLAK
>PSOMNIFERUMXP_026422689
MAIAALKTSCFLRQSKGTEENLFVRFQIQKKSVVSIKSSSTPVMASLSVSSPATIGLSETFAKLKKQGKVAFIP
YITAGDPDLSTTAEALKVLDSCGSDIIELGVPYSDPLADGPVIQAAATRSLARGTNFDAILSMLEEVVPQLSC
PIALFTYYNPILKRGVGKFMTTIKDVGVHGLVVPDVPLEETEILRKEALSNNIELVLLTTPTTPTERMKSIVD
ASEGFVYLVSSIGVTGARSSVSLRVESLLQDIKKASGKPVAVGFGISKPEHVKLVAGWGADGVIVGSAMVK
ILGEAKSPEEGLKELEAFTKSLKAALP
>PBRACTEATUMKAI3832353
MAIAALKTSCFLRQSKGTEGNLFVRFQIQKKSVVSIKSSSTPIMASLSVSSPATIGLSETFAKLKKQGKVAFIP
YITAGDPDLSTTAEALKVLDSCGSDIIELGVPYSDPLADGPVIQAAATRSLARGTNFDAILSMLEGVVPQLSC
PIALFTYYNPILKRGVGKFMTTIKDVGVHGLVVPDVPLEETEILRKEALKNNIELVLLTTPTTPTERMKSIVD
ASEGFVYLVSSIGVTGARSSVSLRVESLLQDIKKASGKPVAVGFGISKPEHVKLVAGWGADGVIVGSAMVK
ILGEAKSPEEGLKELENFTKSLKAALP
>SCOPARIA_DULCIS_AYC63472.1
MAAAALKASSFLQLRSNNYNNNNNKGGISQRRCWSTNNRALKYKPPMAALATAPTVGLSETFSRLKQKG
KVALIPYITAGDPDLSTTAEALKVLDASGSDIIELGVPYSDPLADGPVIQAAATRSLAKGTNFDKIIAMLKEV
VPQLSCPIALFSYYNPILKRGVEKFMTTLQETGIHGLVVPDVPLEETEILRKEASNKNIELVLLTTPTTPKERM
KSIVEASGGFVYLVSAVGVTGARASVSDKVQSLLTEIKEATDKPVAVGFGISKPEHVKQVAGWGADGVIV
GSAMVKVLGEAKSPEEGLKELSLFTKSLKEPLQ
>ANDROGRAPHIS_PANICULATA_XP_051143432.1
MAAALKATCFLQRKTDSSCSGRRLSFAFPTSKSFKYKPPMAALTTSPTLSISETFTKLKQRGEVALIPYITAG
DPDLSITAEALKILDSSGSDIIELGVPYSDPLADGPVIQGAATRALAKGTTFEKIIDMLKDVVPQLSCPIALFT
YYNPILKRGVEKFMTTLKDTGVHGVVVPDVPLEETQILRKEASSKNIELVLLTTPTTPAERMKAIAEASEGF
LYLVSSVGVTGARSSVSQRVESLLRDIKESTNKPVAVGFGISKPEHVKQVAEWGADGVIVGSAIVRLLGEA
KSPEQGLKELETFTKSLKAPLIK
>SESAMUM_INDICUM_XP_011079942
MAAALQTTCFLRLKNNYNVIQCRSSTNRAFKCKPPMATLAAAPTVGLSETFSRLKQKGKVAFIPYITAGDP
DLPTTAEALKVLDSCGSDIIELGVPYSDPLADGPVIQAAATRSLARGTNFDKIIAMLKEVVPQLSCPVALFSY
YNPILKRGTEKFMTTLKDTGIHGLVVPDVPLEETEILRKEASNNNIELVLLTTPTTPTIRMKAIVEASEGFVYL
VSAVGVTGARASVSEKVQSLLQEIKEVTNKPVAVGFGISKPEHVKQVASWGADGVIVGSAMVKILGEAKS
PEEGLNELEAFTKSLKSALL
>SALIX PURPUREA_KAJ6773797
MAVALKSTPSFLQLKKPETHFLLRNKPLIVSTRRFAPMASLTATRSLGIGETFSNLKKQGKVALIPYITAGDP
DLSTTAEALKMLDACGCDIIELGVPYSDPLADGPVIQAAATRSLARGTNFEAITSMLKEVVPQVSCPIALFT
YYNPILKRGIEKFMSTVKDIGVHGLVVPDVPLEETQVLRKEAVKNGLELVLLTTPTTPKERMRAIVEAADG

FVYLVSSVGVTGTRASVSDKVQTLLQDIKETTTKPVAVGFGISKPEHVKQVAGWGADGVIVGSAMVKLLG
EAKSPEEGLKELESFTKSLKAALS
>MANIHOT ESCULENTA_XP_021618706
MSIALKSTTSFLNLKKPETHLPIRFPSYKSSIVSTRRLTPMATLTTAPTLGLRDTFSNLKKQGKVAFIPYITAG
DPDLSTTAEALKVLDSCGSDIIELGVPYSDPLADGPVIQAAATRSLARGTNFNAITSMLKEVVPQLSCPIALF
TYYNPILKRGIEKFMSTVQDIGVHGLVVPDVPLEETEVLRTEAVKHNIELVLLTTPTTPSERMKAIVEASEGF
VYLVSSVGVTGTRASVSNRVQTLLQDIKEVTTKPVAVGFGISKPEHVKQIAEWGADGVIVGSAMVKLLGE
AKSPQEGLKELENLTKSLKSALP

**Data S2.**

Sequences used for phylogenetic analysis of Fig S2A, provided in FASTA format.

>ZEA_MAYS_PWZ03963
MLMARLFLTATGQEQRASLLCTPKHRVAASRRSLRFTTRASSNAGASVSIPKQWYNLIADLPVKPPPPLHP
QTHQPLNPSDLSPLFPDELIRQEVTDERFVDIPEEVIDVYKLWRPTPLIRARRLEKLLGTPAKIYYKYEGTSPA
GSHKPNTAVPQAWYNAAAGVKNVVTETGAGQWGSALSFASSLFGLNCEVWQVRASFDQKPYRRLMMET
WGAKVHPSPSTATEAGKRILEADPSSPGSLGIAISEAVEVAATSADTKYCLGSVLNHVLLHQTVIGEECLEQ
LAALGETPDVVIGCTGGGSNFGGLAFPFLREKLRGRMSPAFRAVEPAACPTLTKGVYAYDFGDTAGLTPL
MKMHTLGHGFVPDPIHAGGLRYHGMAPLISHVYELGFMDAVAIQQTECFQAALQFARTEGIIPAPEPTHAI
AAAIREALECKRTGEEKVILMAMCGHGHFDLAAYEKYLRGDMVDLSHPAEKLEASLAAVPKV
>PAPAVER_SOMNIFERUM_XM_026582418
MRIVSATKKRTQLKLSPDYYNRAATARSIMCSATNKPTEIPRQWYNIVADLPVKPPPALDPETLQPVKPEEL
SHLFPDELLKQDNSTERFIDIPEEVLDIYSLWRTTPLLRAKRLEKLLGTPARIYYKYEGVSPAGSHKPNTAIPQ
AYYHAKQGTKKLVTETGAGQWGSSLAFACNLFGLGCEIWQTRASYDGKPYRKLMMQTWGAKVHPSPSD
KTNVGRSILQKDPSCPGSAAIASTEAVEICTENADTKYCMGSVLNHVVLHQTIIGEECIRQMEALGESPDVII
GCSGGGSNFAGLAFPFLREKLEGKINPIIRAVEPSACPSLTKGVYAYDSSDTTGLTPLMKMHTLGHDFIPDPV
HAGGLRYHGMAPLVSHVYELGLMEAVAMPQLECFDGAIQFARSEGIIPAPESGHAIAEAIKEALHCKETGE
SKVILLLVSGHGHFDLKSFEKYLLGNMVNLSCTDETLRASLAGSSIPSGFELKC
>POPULUS_TRICHOCARPA_XM_006373427.3
MAIQATFIADPVLISSPSRISIRGWEQCIGSFVLKTRPRHPRLSNGGRVRARASLNADLKAVGIPHQWYNVV
ADLSVKPPPPLHPKTFEPVKPEDLSPLFPDELIRQEASTDKFIDIPEEVLDIYSLWRPTPLIRAKRLEKLLDTPA
RIYYKYEGGSPAGSHKPNTAVPQVFYNAQQGIKNVVTETGAGQWGCSLAFACSLFGLDCEVWQVRASYD
QKPYRRLMMETWGAKVHPSPSSITETGRRILQMDPSSPGSLGIAISEAVEVAAKNADTKYCLGSVLNHVLL
HQTVIGEECIKQMEAIGETPDVIIGCTGGGSNFAGLSFPFIREKLNGKINPVIKAVEPAACPSLTKGVYAYDY
GDTAGMTPLMKMHTLGHDFIPDPIHAGGLRYHGMAPLISHVYELGFMEAMAIPQIECFRGAIQFARSEGLIP
APEPTHAIAATIREALHCKETGEAKVILMAMCGHGHFDLKSYEKYLQGKMVDLSFDEEKIRASLDKVPQVT
RN
>ARABIDOPSIS_THALIANA_AT5G38530_TSB_TYPE2
MASQLLLPPNQFTKSVSPQVFITGDCQGFSDLTLKRKSNQATRVSNGSSLRVKAALRSTHNKSVVEIPKQW
YNLVADLSVKPPPPLHPKTFEPIKPEDLAHLFPNELIKQEATQERFIDIPEEVLEIYKLWRPTPLIRAKRLEKLL
QTPARIYFKYEGGSPAGSHKPNTAVPQAYYNAKEGVKNVVTETGAGQWGSSLAFASSLFGLDCEVWQVA
NSYHTKPYRRLMMQTWGAKVHPSPSDLTEAGRRILESDPSSPGSLGIAISEAVEVAARNEDTKYCLGSVLN
HVLLHQTIIGEECIQQMENFGETPDLIIGCTGGGSNFAGLSFPFIREKLKGKINPVIRAVEPSACPSLTKGVYA
YDFGDTAGLTPLMKMHTLGHDFIPDPIHAGGLRYHGMAPLISHVYEQGFMEAISIPQIECFQGAIQFARTEGI
IPAPEPTHAIAATIREALRCKETGEAKVILMAMCGHGHFDLTSYDKYLKGELVDLSFSEEKIRESLSKVPHV
V
>LAMIUM_GALEOBDOLON_LG_66
MSGIKIINSTSVSSHLQGQICPKMPTTSQSNWPTVGDKRSRISCVMTQPNIRHPTPKSDLLCLIDQNQESSPST
GKFGRFGGVFVPETLVTCLTNLAIEFKLLMRDPHFQEELAIGLRDYVGRETPLYFAQRLTDHYKNDKGEGP
EIYLKREDLNHGGAHKINNALAQAMLAKRMGRKSVVAATGAGQHGVATAAVCAKLDLECTIIMGKLDM
ERQPSNVLLMKHLGAQVKSVEGTFKDATSEAIRVWVGDLERSYYLAGTAVGPHPCPSMVREFQSVIGKET
RKQAMEKWGGKPDVVVACVGSGSNALGIFHEFVRDEQVRLIGVEAGGTGINGDKHSATLAKGEVGVYHG
AMTYLLQDDEGQIIVPHSIGVGLEYPGVSPELSFLRDIGRAEFYAVTDEEALDACSLLCRLEGIFPALESAHA
LAYLGRLCRTLPNGAKVVVVNLSGRGDKDAXTVFKYQQHKNT
>LAMIUM_GALEOBDOLON_TRINITY_DN6977_C0_G1_I1
MTFSSSSSACRTQSHLPLHIRSPSLPPARFFQTTPYSLSNSRSPPTLSAVTRMAATAVDKELVAAEAQALLRP
DAFGRFGKFGGKYVPETLMSALTELEAAFKSLADDVEFQKELDGILKDYVGRESPLYFAERLTEHYKRPDG
TGPHVYLKREDLNHTGAHKINNAVAQALLAKRLGKKRIIAETGAGQHGVATATVCARFGLECIIYMGAQD
MERQALNVFRMRLLGAEVRAVHSGTATLKDATSEAIRDWVTNVESTHYILGSVAGPHPYPMMVREFHAVI
GKETRKQALEKWGGTPDVIVACVGGGSNAMGIFHEFVDDKEVRLIGVEAAGFGLDSGKHAATLTRGEIGV
LHGAMSYLLQDEDGQIIEPHSISAGLDYPGVGPEHSFLKDIGRAEYHSITDEEALEAFKRLSKLEGIIPALETS
HAIAYLEKLCPTLPDGAKVVLNCSGRGDKDVQSVIKHLNL
>SALVIA_SPLENDENS_XM_042189368.1
MNLQALSLHHAISLPSLNPNNSTTLGFRLPSHRGFKVTSVMSPQHTWTQTWKTDVLRLIDETEESLFKTGKF
GRFGGVFVPETLITCLNMLLSEFSFCLNDPHFQGELATALRDYVGRETPLYFAQRLTDHYKNSRGEGPEIYL

KREDLNHGGAHKINNAIAQAMLAKRMGRRSIVAATGAGQHGVATAAACAKLDMECTIIMGDVDVARQS
HNVRLMKLLGAQVKSIKGSFKDATSEAIREWVGDLERKYYLAGTAVGPHPCPAMVREFQSVIGKETRKQA
MEKWGGKPDVLVACVGSGSNALGLFHEFVRDEDVRLIGVEAAGTNIDGGTHSATLAKGEIGVYHGAMTY
LLQDDDGQIVGPHSVGVGLEYPGVSPELSFLRDIGRAEIHTATDEEALDAYALLCRLEGICPALEASHALAH
LGKLCSTLPSGTKVVVNCSGRGDKDIETVFKHQQQKSGGELTTFL
>SALVIA_SPLENDENS_XM_042206891
MAFAASSSSAATARTSAQSDLAFRIPSNPPKFAKFAPSISRRSPPISCKMAATAVVDMELDAEALQRPDAFG
RFGRFGGKYVPETLMAALTELEAAFNSLATDHEFQKELDGILKDYVGRESPLYFAERLTQHYKRHDGTGP
HVYLKREDLNHTGAHKINNAVAQALLAKRLGKKRIIAETGAGQHGVATATVCARFGLECIIYMGAQDME
RQALNVFRMRLLGAEVRAVHSGTATLKDATSEAIRDWVTNVETTHYILGSVAGPHPYPMMVREFHKVIGQ
ETRKQALEKWGGKPDVIVACIGGGSNAMGIFHEFVEDEDVRLIGVEAAGFGLDSGKHAATLTKGEIGVLH
GAMSYLLQDEDGQIIEPHSISAGLDYPGVGPEHSFLKDLGRAEYHSITDEEALEAFKRLSRLEGIIPALETSHA
LAYLEKLCPTLPDGTKVVLNCSGRGDKDVHTAINYLKL
>APHELANDRA_SQUARROSA_AS_144
MPASAAASRFISCGAAQADLPSQLNLQSLLPLKSSRIAPTRPFDSRRPLVVCAVNMAASAVTAVEKETAAA
GEALQRPDSSGRYGKFGGKYVPETLMYALSELEAAFKALSNDDQFQKELSGILKDYVGRESPLYFAERLTE
HHKRADGTGPHIYLKREDLNHTGAHKINNAVAQALLAKRLGKKRIIAETGAGQHGVATATVCARFGLQCII
YMGAQDMERQALNVFRMRLLGAEVRGVHSGTATLKDATSEAIRDWVSNVETTHYILGSVAGPHPYPMLV
RDFHAVIGKETRRQALEKWGGKPDVLVACIGGGSNAMGLFHEFIDDKDVRLIGVEAAGFGLDSGKHAATL
SKGEVGVLHGAMSYLLQDEDGQVIEPHSISAGLDYPGVGPEHSFLKDIGRAEYHSLTDEEALQAFKRLSRL
EGIIPALETSHALGYLEYLCPTLPDGTKVVVNCSGRGDKDVHTALKYLNM
>APHELANDRA_SQUARROSA_AS_143
MSYSKCLPSNSFLHSNGFYSNSDPKLATRRLNFGRKIAGDKSLTVSSVMTTQDVRTPLNDDQATHSRIRSD
VVRLIEQADEKLSSTGKFGRFGGVFVPETLITCLNKLAAEFNLILHDRGFQAELRTALRDYVGRETPLYYAK
RLSDHYRNGKGEGPDIYLKREDLNHGGAHKINNAIAQAMIAKRMGRMRVVAATGAGQHGVATASACAQ
LGLECTVFMGNVDMERQPSNVLLMKILGAQIKSVEGSFKDATSEAIRHWVGDLDNGYFLTGMAVGPHPLP
TMVREFQAVIGKETRRQAREKWGGKPDVVVACVGSGSNALGIFHEFVKDEDVRLIGVEAAGSGIDTGKHS
ATLSTGDVGVYHGAMSYLLQDDEGQIIGPHSIGVGLEYPGVSPELSFLKDIGRAEFHTVTDEEALDAYAVL
CRLEGIIPALEAAHALAYLGKLCKTLPDGAKVVVNCSGRGDKDAATVFNHQQQQKQ
>ANDROGRAPHIS_PANICULATA_XM_051272759.1
MSCMNNLHIQSSISPIGKNPNLGNDRLIKFKNVNYEGSAMGSRSVATSPADSSNFNASRKRNSGDLKLIDEV
STTGKFGRFGGVFVPETLITCLRKLAAEFNLILHDHQFQSELAAALRDYVGRETPLYYAERLSEHYRNGAG
EGPEIYLKREDLNHGGAHKINNALGQAMIAKRMGRRRVVAATGAGQHGVATAAVCAKLGLECTVFMGD
ADMLRQSFNVTLMKHLGAELKSVGGTFKDATSEAIRSWVGDLDGGYYLTGTAVGPHPLPSMVREFQSVIG
KETRRQAAEKWGGPPDVVVACVGSGSNALGIFHEFLTDENVRLVGVEAAGTGIDGGRHSATLAKGEVGV
YHGAMSFLLQDNEGQIVVPHSIGVGLEYPGVSPELSFLKEIGRAEFHTVTDEEALQAYGRLCRLEGIIPALEA
AHAVAHLDRLCPTLPGGAKVVVNCSGRGDKDAAAVIELLSSTTLNSEKGKKFGSNSYQELNPT
>ANDROGRAPHIS_PANICULATA_XP_051133934
MSVSSSLASSFTSRRTTADLPSQLRRPLSVAIKANRISPILPSGAAASPPIACRLPSRMSAMAVEKEVVLAEEI
LQRPDSFGRYGKFGGKYVPETLMYALTELEAAFNALSTDEEFQKELSGILKDYVGRESPLYFAERLSEHYK
HSDGTGPHIYLKREDLNHTGAHKINNAIAQALLAKRLGKQRIIAETGAGQHGVATATVCARFGLPCVIYMG
AQDMERQALNVFRMRLLGAEVRGVHSGTATLKDATSEAIRDWVANVETTHYILGSVAGPHPYPMMVRDF
HAVIGKETRKQALEKWGGKPDVLVACVGGGSNAMGLFHEFVDDKDVRLIGVEAAGFGLDSGKHAATLT
KGEVGVLHGAMSYLLQDDDGQIIEPHSISAGLDYPGVGPEHSFLKDVGRAEYHSITDEEALEAFKRLSRLEG
IIPALETSHAIGYLEYLCPTLPDRAKVVLCCSGRGDKDVHTALNYLKV
>OLEA_EUROPAEA_XM_023030728
MAGNNLQSTISVHGRTLYPTCKLLNSSKMLGFSLQSRTDRGLIVCSAVGHNTLNSDELGLRNPTRTSEILELI
DQGEKTLANSGKFGRFGGIFVPETLITCLKKLEVEFNWVLHDPQFQEELSIALKDYVGRETPLYFAERLTNY
YRNKNGEGPEIYLKREDLTHGGAHKINNAMAQAMLAKRMGRKSIVAATGAGQHGVATAAACAKLGLDC
TIFMGSIDMQRQSSNVLLMEHTGAQVKSVAGTFKDATSEAIRSCVGNLQGSYYLAGTAVGPHPCPSMVRE
FQTVIGKETRKQAMEKWSGKPDVLVACVGSGSNALGLFHEFIRDEDVRLIGVEAAGYGIDSGKHSATLAK
GEVGVYHGAMSYLLQDDEGQIIGPHSIGVGLEYPGVSPELSFLKDIGRAEFYSITDEEALEAYTRLCRLEGIF
PALEAAHALAYLEKLCPTLASGTKVVVNCSGRGDKDAGTVFKYQHKMEGEQEQSE
>OLEA_EUROPAEA_XM_023002444
MAFFAAAPCRARTHLASSADCYSLLPLKFDRITPPCHLNPKRHSVFCTLSKQTMAEHHLEPVVLQRPDSFG
RFGKFGGKYVPETLMYALTELETAFNSLASDQKFQNELDGILRDYVGRESPLYFAERLTEHYKRPDGRGPQ

IYLKREDLNHTGAHKINNAVAQALLAKRLGKKRIIAETGAGQHGVATATVCARFGLECIIYMGAQDMERQ
ALNVFRMRLLGAEVRPVHSGTATLKDATSEAIRDWVTNVESTHYILGSVAGPHPYPMMVREFHAVIGKET
RKQALEKWGGKPNVLVACVGGGSNAMGIFHEFVDDKDVRLIGVEAAGFGLDSGKHAATLTKGEVGVLH
GAMSYLLQDDDGQIIEPHSISAGLDYPGVGPEHSFLKDIGRAEYYSITDEEAIEAFKRLSRLEGIIPALETSHA
LAYLEKLCPTLPDGTKVVLNCSGRGDKDVQTAIKLLKL
>WRIGHTIA_RELIGIOSA_WR_6
MACSQSLSLFFPHGGNLALGKPKLPAGLISFGVPSSYTKGYAGSVSCSAVGNDIAAVAGSLSLEALTEKWT
TMTPRPNHTEKETPKLTSGKFGRFGGKFVPETLITCFNMLEAEFNLVLTDADFQRELALALRDYVGRESPLY
FAKRLTDHYKNSNGEGPEIYLKREDLVHGGAHKINNAIAQAMLAKRMGRSSIVAATGAGQHGVATASAC
AKLSLECTIFMGTTDMERQQSNVLLMKHLGAQVKAIDGSFKDAMSEAIRSWVGNLDSSYLLIGTAVGPHP
CPTMVREFQSVIGRETRKQAMEKWGGKPDVLVACVGSGSNALGLFHEFIRDEDVRLVGIEAAGTGLDAGA
HSATLARGEVGVYHGTMSYLLQDEEGQIIGPHSIGVGLEYPGVSPELSFLKDIGRAEFYSVTDEEALDAYGR
LCRLEGICPALEASHALAYLGKLCPTLRNGTKVVVNCSGRGDKDAVTVFNRKLEMTERQVN
>WRIGHTIA_RELIGIOSA_WR_7
MAFYTAAMRPRTTATPRQLPELHRSFNATTYSQKPLKFNQFTSISISCTLAKEVMVTKEEVEPVVLQRPDSL
GRFGKFGGKYVPETLMYALSQLEDAFRTLSANREFQRELEGILKNYVGRESPLYFAERLTEHYRRPNGKGP
YIYLKREDLNHTGAHKINNAVAQALLAKHLGKKRIIAETGAGQHGVATATVCARFGLQCIIYMGAQDMER
QALNVFRMRLLGAEVRAVHSGTATLKDATSEAIRDWVTNVESTHYILGSVAGPHPYPMMVREFHAVIGKE
TRKQALEKWGGNPDVLVACVGGGSNAMGLFHEFIDDKDVRLIGVEAAGYGLDSGKHAATLTKGEVGVL
HGAMSYLLQDEDGQIVEPHSISAGLDYPGVGPEHSFLKDIGRAEYYSVTDEEALEAFKRVSRLEGIIPALETS
HALAYLEKLCPTLPDGTKVVVNFSGRGDKDVQTAIKHLKL
>COFFEA_EUGENIOIDES_XM_027303292
MIMASSPRNVFLQGGNTTEFSEPVLKLRRLPFSAPLGSPYIKAADRPRAPISCSAVADKMPATVGGALDSNS
LTTRRLMTQTFIDQAELMTVSSTGKFGRFGGKFVPETLISCLSKLEAEFILALHDENFQEELATALRDYVGR
ESPLYFAKRLTDYYKNSEGKGPEIYLKREDLVHTGAHKINNAIAQAMLAKRIGRKRIVAATGAGQHGVAT
AAACAKLSLECTIFMGSADVGRQGANVLLIKNLGAQVKVVDGSFKEAMSEAIRDWVGDLETSYFLSGTAV
GPHPCPTIVREFQSVIGKETRKQAEEKWGGKPDVLVACVGSGSNALGMFHEFIRDEDVRLIGVEAAGVGLD
SGAHSATLARGDVGVYHGSMSYLLQDEEGQIIGPHSIGVGLEYPGVSPELSFLKDNGRAEFYAVTDDEALD
AYRRLCQLEGIFPALEASHALAYLEKLCPGLEDGAKVVVNCSGRGDKDAATVYKHNEERV
>COFFEA_EUGENIOIDES_XM_027314528
MAAFSTASVTPETAATPLSLPKPYRPSSAAFLRSLPRKFNRSPKTTSPSVFCTLTKQPMTSTEEEVILQRPDSF
GRFGKFGGKYVPETLMYALSELEDAFKSLSADNEFQNELDGILRDYVGRETPLYFAERLTEHYRRPSGEGP
HIYLKREDLNHTGAHKINNAIGQALLAKRLGKKRIIAETGAGQHGVATATVCARFGLECVIYMGAQDMER
QALNVFRMRLLGAEVRPVHSGTATLKDATSEAIRDWVTNVESTHYILGSVAGPHPYPVMVREFHAVIGKE
TRKQALEKWGGKPDVLVACVGGGSNAMGLFDDFVDDKDVRLIGVEAAGFGIESGKHAATLTKGEVGVL
HGAMSYLLQDEDGQIIEPHSISAGLDYPGVGPEHSFLKDIGRAEYFSVTDEEALEAFKRLSRLEGIIPALETSH
AVAYLEKLCPTLPDGAKVVLNCSGRGDKDVQTAIKHLKL
>SOLANUM_LYCOPERSICUM_XM_004248265
MAFSSTVSPKHCRLSSAASSSSSFPKFQIPLKINNITPCHSSSKFPPSIVCVLTEKQSMAAQAVDEPAAIQRPDS
FGRFGKFGGKYVPETLMHALDELETAFKSLATDEAFQKELDGILRDYVGRESPLYFAERLTEHYKRPNGEG
PLIYLKREDLNHTGAHKINNAVAQALLAKRLGKKRIIAETGAGQHGVATATVCARFGLECIIYMGAQDME
RQALNVFRMRLLGAEVRAVHSGTATLKDATSEAIRDWVTNVETTHYILGSVAGPHPYPMMVRDFHAVIG
KETRKQALEKWGGKPDVLVACIGGGSNAMGLFHEFVDDEDVRLIGVEAAGFGVDSGKHAATLTKGEVGV
LHGAMSYLLQDEDGQIVEPHSISAGLDYPGVGPEHSFLKDLGRAEYYSITDEEALEAFKRLSRLEGIIPALET
SHALAYLEKLCPTLPNGTRVVLNCSGRGDKDVQTAIKYLKV
>SOLANUM_LYCOPERSICUM_XM_004248350
MACNINVESILGQGIFATTSSKKLQAFPSHHTYKANVISCVAIGPTPIPQLPWKLVFHEKERQSLLSNEKFGIY
GGKFVPETLISPLTKLDYEFNSALRDPQFQMNLQVALKDYVGRETPLYFAQRLTDYYKSLNKGIGPDIYLK
REDLNHGGAHKINNAIAQAMLAKRMGCKNVVASTGAGQHGVATAAACAKLSLECTIFMGSLDMERQPSN
VLLMNHLGAKVKCVEGSFKDAMSEGIRNWVNNLETSYFLAGAAIGPHPCPTMVREFQSIIGKETRKQAMD
KWGGKPHVLVACVGSGSNALGLFHEFIQDVDVRLIGVEAGGIGLDSGKHSATMARGEVGVYHGAMSYLL
QDEEGQIIGPHSIGVGLEYPGVSPELSYLKDIGRAEFSTVTDEEAIKAYKRLCILEGIFPALESCHALAFLDKL
CSTLKDGEKVIVNLSGRGDKDAEAVFNHTPKHK
>NICOTIANA_TABACUM_XM_016633459
MAFSSSTAQTASPLSSKHCCRLSSSAASSASYFPKFQIPFKFDKTTSCPSSISCVLTKQESMAAQEAAEPAVLL
RPDSFGRFGKFGGKYVPETLMHALDELETAFKSLATDEAFQKELDGILRDYVGRESPLYFAERLTEHYKRP

DGEGPLIYLKREDLNHTGAHKINNAVAQALLAKRLGKKRIIAETGAGQHGVATATVCARFGLECIIYMGAQ
DMERQALNVFRMRLLGAEVRGVHSGTATLKDATSEAIRDWVTNVETTHYILGSVAGPHPYPMMVREFHA
VIGKETRKQALEKWGGKPDVLVACVGGGSNAMGLFHEFVDDKDVRLIGVEAAGFGIDSGKHAATLTKGE
VGVLHGAMSYLLQDEDGQIVEPHSISAGLDYPGVGPEHSFLKDLGRAEYYSITDEEALEAFKRLSRLEGIIPA
LETSHALAYLEKLCPTLPNGTKVVLNCSGRGDKDVHTAINYLKV
>NICOTIANA_TABACUM_XM_016602914
MACNIEVIVRQAKIAEPRLSYSRKWKGKFAIFSVATGPSRVTELPGKLVYHEKKRAIFSNEKFGIFGGKFVPE
TLISSLTKLDYEFNSALHDPQFQMVLGVALRDYVGRETPLYLAERLTDNYKSRNGGKGPDLYLKREDLNH
VGAHKINNAIAQTMLAKRMDCKNIIAATGAGQHGVATAAACAKLSLECTVFMGSLDMERQPSNVILMKH
LGAKVKSVKGSFKDAVSEGIRHWVNNLETSYFLGGAAIGPHPCPTMVREFQSVIGKETRKQAMEKWGGKP
DVLVACVGSGSNALGLFHEFIEDKDVRLIGVEAGGVGLDTGKHSATMARGQVGVYHGAMSYLLQDDEGQ
IIEPHSIGVGLEYPGVSPELSFLKDTGRAEFYTVTDEQALEAYKRLCRLEGIFPALEASHALAFLDRLCPTLED
GEKVVVNLSGRGDKDAATVFNHTTKNE
>CUCURBITA_PEPO_XM_023669321
MVASVELSCKILTIPPSSSHSNSKISSSFSRFPVHFRRFLPNSSPVFRSCAVTCTLTRETSLAMEDNKTQNLGQ
RPDSFGRFGRFGGKYVPETLMHALAELETAFHSLSGDQEFQKELDGILRDYVGRASPLYFAERLTEHYRRP
NGEGPHIFLKREDLNHTGAHKINNAVAQALLAKRLGKKRIIAETGAGQHGVATATVCARFGLECIIYMGAQ
DMERQALNVFRMRLLGAEVRPVHSGTATLKDATSEAIRDWVTNVETTHYILGSVAGPHPYPMMVRDFHA
VIGKETRQQALEKWGGKPDVLVACVGGGSNAMGLFHEFVNDKDVRLVGVEAAGFGVDSGKHAATLTKG
EIGVLHGAMSYLLQDDDGQIIEPHSISAGLDYPGVGPEHSYLKDAGRAEYYSVTDDEALEAFKRLSRLEGIIP
ALETSHALAYLEKLCPTLADGTKVVLNCSGRGDKDVHTAIKHLQV
>CUCURBITA_PEPO_XM_023686897
MASNIVHNIAAAVTNQLDATRGIARSHVKLGNGIGSPTFTPTRKPTPALKMQPVIDDHPKNSKQTIHEGKFG
KFGGKFVPESLITCLAKLEAEFNLVLNDSKFQEELEVALRDFVGRETPLYHAERLTKYYKDEEGKGPEIYIK
REDLNHCGAHKMNNAIAQVMIAKRMGRRRVVAATGAGQHGVATAAACAKHDLECTIFMGSEDIKKQSS
NVVLIKLLGAQVKSVEGNFKDASSEAIREWVGNLEMSYYLTGTVVGPHPCPAMVREFQSVIGKETRKQAM
EKWGGLPEVLVACIGSGSNALGLFNEFMNEEDVRLIGVEAAGFGLDSGKHSATLSKGHVGVYHGALSYLL
QDDEGQILVPHSVGVGLEYPGVGPELSFLKESGRAEFHTAIDKEAVEAYKRVCKLEGIFPSLEASHAFAYLD
KLCPTLTDGCKVVVNCSGRGDKDAAIVFNYGHHQEC
>CUCUMIS_MELO_XM_008456305
MVASTALNCKIPTISPSPSHSNNKISSSIPHFPVRFRRVSSNPSPLLRSNAVSCTLTREPSLAMEDKLHTLSLQQ
RPDSFGRFGRFGGKYVPETLMHALTELEAAFYSLAGDQDFQKELDGILRDYVGRESPLYFAERLTEHYRRS
NGEGPHIFLKREDLNHTGAHKINNAVAQALLAKRLGKKRIIAETGAGQHGVATATVCARFGLECIIYMGAQ
DMERQALNVFRMRLLGAEVRPVHSGTATLKDATSEAIRDWVTNVETTHYILGSVAGPHPYPMMVRDFHA
VIGKETRKQALEKWGGKPDVLVACVGGGSNAMGLFHEFVNDEDVRLVGVEAAGFGLDSGKHAATLTKG
EVGVLHGAMSYLLQDDDGQIIEPHSISAGLDYPGVGPEHSYLKDLGRAEYHSVTDDEALEAFKRLSRLEGII
PALETSHALAYLEKLCPTLPDGTKVVLNCSGRGDKDVQTAIKYLQV
>CUCUMIS_MELO_XM_008443142
MACNMVKNSAITNQLVAKPHVNFGKLGHGANTLATTKRNYMGTIKMQLVTDNPKKYQNLGIGELGKFG
KFGGKFVPESLITCLGKLEAEFNLVLNDSKFQEELEVALRDFVGRETPLYYAERLTKHYKNEEGKGPEIYIK
REDLNHCGAHKMNNAIAQVMIAKRMGRKSVVAATGAGQHGVATAAACAKHDLDCTIFMGSEDINKQSS
NVLLIKLLGAKVKSVEGNFKDASSEAIREWVGNLETSYYLTGTVVGPHPCPAMVREFQSVIGKETRRQAKE
KWGAKPDVLLACIGSGSNALGLFHEFINEKDVRLIGVEAAGFGLDSGKHSATLSKGHVGVYHGALSYLLQ
DDEGQILNPHSVGVGLEYPGVGPELSFLKESGRAEFETASDTEAVEAYKRLAKLEGIFPSLEASHAFAYLHK
LCPTLPDGCKVVVNCSGRGDKDAAIVFNYHQQNH
>CICER_ARIETINUM_XM_004499981
MASSITTLNSSRLLPISKEQNYPSSHFHSNLLKVSSFPSRTSSSSYSLSCSVTKDPSSVLPLLDHTKLDNGSVV
HQRPDSFGRFGKFGGKYVPETLMHALTELEASFYSLTADEDFQRELAGILKDYVGRESPLYFAERLTEHYK
RPNGEGPHIYLKREDLNHTGAHKINNAVAQALLAKSLGKKRVIAETGAGQHGVATATVCARFGLECIIYM
GAQDMERQSLNVFRMRLLGAEVRPVHAGTATLKDATSEAIRDWVTNVETTHYILGSVAGPHPYPMMVRE
FHAVIGKETRKQALEKWGGKPDVLIACVGGGSNAMGLFHEFVDDKDVRLIGVEAAGFGLDSGKHAATLT
KGEVGVLHGAMSYLLQDDDGQIVEPHSISAGLDYPGVGPEHSFLKDLGRAEYHSITDNEALEAFKRVSRLE
GIIPALETSHALAYLEKICPTLPNGTKVVVNFSGRGDKDVHTAIKYLKM
>CICER_ARIETINUM_XM_004511284
MTTCKFQGAIVRIDNHKKGGPKERASKGLQTVVTRHHFKVKVPQLTTTYTPLPKTPLKDKAKEIIIDTNSTN
SAGKFGRFGGKFVPETLIQCLNQLEAEFNNTLHDQVFQTELSTTLRDYVGRETPLYHAQRLSEYYKSKNGG

IGAEIYLKREDLNHSGSNKMNNALAQVMIAKRIGRKSVVTATGSGKHGLATAAACAKFGLECIVFMAAKD
KDRYSSNVRLMNLLGARVEVVNGSFKDASSEAFRCWVGDIENNYHLTGSAVGPHPCPTMVREFQSVIGKE
TRKQALEKWGGKPDVVVACVGTGSNALGIFHEFIQDTDVRLFGVEGGGLGLDSGKHSSTLAKGEVGVYH
GSITFLLQDDYGQIVPPHSIAAGMECPGVGPELSYLKESGRAEFFVATDEEALDAYERLCKLEGIFPSLEAAH
ALAILDKLVPTLSNKNVVVNCSGRGDNDDAAIFFNTCIPQ
>ARACHIS_HYPOGAEA_XM_025798338
MMTCKLQLGAFASSSNHSRQQGAPNKVRSCLMVATDDHQDTFFKTNLPTKLAKTYVPTIKPLLEKELQES
NNNTDAPIINSGKFGRFGGKFVPETLVVCLSQLEAEFNKALHDEAFKAELAEALKDFAGRETPLYYARRLS
EYYKMNNNGKGPDIYLKREDLNHGGSHKMNNALAQAMIAKRMGRKSVVTATAAGQHGIATAAACAKL
GLECTVFMAAKEMEKESSNVQLMKLLGAKVEGVNGSFKDAASEAFRCWVGEMENSYHLTGSAVGPHPC
PTMVREFQSVIGRETRKQALEKWGGKPDVLVACVGTGSNALGLFHEFLEDKDVRLIGVEAGGCGLESGRH
SSTLATGEVGVYHGALSYLLQDQDGQIIGPHSIAPGMEYPGVSPELSYLKETGRAEFCVATDQEALDAYER
LCKLEGIFPSLEAAHALAILDNLVPTLCDGSKVVVNCSGCGDKDAAVVFDRKLFHGKF
>ARACHIS_HYPOGAEA_XM_025770464
MASSITSSSSRLFPITRESQPTSHSTLNFSKFASFSSSSSASGSKASSYISCSLTRDPSVLPLEEQSKLSNGSVLF
QRPDSLGRFGKFGGKYVPETLIHALTELEAAFHSLSADEDFQKELAGILRDYVGRESPLYFAERLTEHYKRA
NGEGPHIYLKREDLNHTGAHKINNAVAQALLAKCLGKKRIIAETGAGQHGVATATVCARFGLECIVYMGA
QDMERQALNVFRMRLLGAEVRPVHSGTATLKDATSEAIRDWVTNVETTHYILGSVAGPHPYPMMVREFH
AVIGKETRKQALEKWGGKPDVLIACVGGGSNAMGLFHEFVDDKDVRLIGVEAAGFGLDSGKHAATLTKG
EVGVLHGAMSYLLQDDDGQIVEPHSISAGLDYPGVGPEHSFLKDIGRAEYYSITDEEALEAFKRVSRLEGIIP
ALETSHALAYLEKVCPTLPNGAKVVVNFSGRGDKDVHTAIKYLKV
>DURIO_ZIBETHINUS_XM_022878630
MAASAAITAAIRLSNPLTSSSASSSSSSSSFSSSISPFNFNNFASPVTPVKSFSVNCTLTREPVPAAPVSMESDP
TGWQRPDTFGRYGKFGGKYVPETLMYALSELETAFHSLSNDEKFQAELAGILKDYVGRESPLYFAERLSEH
YKRPNGEGPDIYLKREDLNHTGAHKINNAVAQTLLAKRLGKNRIIAETGAGQHGVATATVCARFGLQCIV
YMGAQDMERQALNVFRMRLLGAEVRAVHSGTATLKDATSEAIRDWVTNVETTHYILGSVAGPHPYPMM
VREFHAVIGKETRKQALEKWGGKPDVLVACVGGGSNAMGLFHEFINEKDVRLIGVEAAGFGLDSGKHAA
TLTRGEVGVLHGAMSYLLQDEDGQIVEPHSISAGLDYPGVGPEHSFLKDKGRAEYYSVTDEEALEAFKRLS
QLEGIIPALETSHALAYLEKLCPTLPNGTKVVVNCSGRGDKDVHTAIKHLQV
>DURIO_ZIBETHINUS_XM_022869068
MACNIKALSLNHQIVSSARNLTVDRKMGTASAVSVPVFPDTVTKTTTLSIKNPQSTEKLLTKTVDGKEVYS
QGKFGRFGGKFVPETLISCLGKLEAEFNLVLHDSEFQEELAIALRDYVGRETPLYFAQRLTNHYKNSKGEGP
EIYLKREDLNHVGAHKINNTIAQAMIAKRMGRKSIVAATGAGQHGVATAAACAKLSLECTIFMGSADMEK
QASNVLLMKLLGAKVEPVEGTFKDASSQAIREWVGNLETSYHLIGTVVGPHPCPTMVREFQSVIGKETRRQ
AMEKWGGKPDILVACVGSGSNALGLFHEFIKDEDVRLIGVEAGGFGLDSGKHAATLARGDVGVYHGAMS
YLLQDEEGQILGPHSIGVGLEYPGVGPEVSFLKETGRAEFYSATDKEAIDAYRRLCQLEGIFPALEASHALAF
LEKLCPSLANGTKVVVNISGRGDKDADIVFQYEADRLSGLMA
>HIBISCUS_SYRIACUS_XM_039201377
MAAAATSTAALRPRNPFFSSSVYSSSKFPVNINKVEPLTSPVKKFSIYCTLTREAVPADTLPMDSDPNGWQR
PDSFGRFGKFGGKYVPETLMSALSELETAFHSLSKDDKFQEELAGVLKDYVGRESPLYFAERLSEHYKRPN
GEGPDIYLKREDLNHTGAHKINNAVAQALLAKHLGKKRIIAETGAGQHGVATATVCARFGLQCIVYMGAQ
DMERQELNVFRMRLLGAEVRAVHSGTATLKDATSEAIRDWVTNVETSHYILGSVAGPHPYPMMVREFHG
VIGVETRKQALEKWGGKPDVLVACVGGGSNAMGLFHEFVDDKDVRLIGVEAAGFGLDSGKHAATLTKGE
VGVLHGAMSYLLQDEDGQIVEPHSISAGLDYPGVGPEHSFLKDVGRAEYHSVTDEEALEAFKRLSRLEGIIP
ALETSHALAYLEKLCPTLPNGTKVVVNCSGRGDKDVHTAIKYLKV
>HIBISCUS_SYRIACUS_XM_039192750
MAFSRNALSFNHQLISGSTTKHPTVDTKAGTTSVVSLPMIPQTLPVLTINNPLSSVGTVDKKEVVKNIPGKF
GKFGGKYVPETLITCLGKLEAEFNLVLHDSGFQEDLATALRDYVGRETPLYFAQRLTDHYNKNSHGEGPEI
YLKREDLNHGGAHKINNAIAQAMIAKRMGRKTIVTATGAGQHGVATAAACAKLSLDCTIFMGATDMEKQ
ASNVQLMKLLGARVEPVDHGTFKDASSEAIRAWVGELETSYYLTGTAVGPHPCPSMVREFQSVIGKETRR
QAMEKWGGKPDVLVACIGSGSNALGLFHEFIKDEDVRLIGVEAGGFGLDSGKHAATLARGDVGVYHGAM
SYLLQDDEGQILGPHSIGVGLEYPGVGPEVSFLKETGRAEFHTATDKEAVDAYRRLCRLEGIFPALEASHAL
AFLEKLCPTLPHGTKVVVNISGRGDKDAAIVSQY
>ARABIDOPSIS_THALIANA_AT5G28237_TSB_3_4
MSSSKIQVRGQPLLRVPARNHRMTHLVVCGVSTKRHHREINALSSNSGPSLDSVPTRTDKRQFLRGDGNGK
FGRFGGKFVPETLMSRLIELEDEFNFVRCDHEFQEELTTALRDYVGRETPLYFAERLTEHYKNIVPTIEGGPE

IYLKREDLSHCGSHKINNALAQAMISRRLGCSRVVAATGAGQHGVATAAACAKLSLECTVFMGAADIEKQ
SFNVLSMKLLGAQVISVEGTFKDASSEAIRNWVENLYTTYYLSGTVVGPHPCPIIVREFQSVIGKETRRQAK
QLWGGKPDVLVACVGSGSNALGLFHEFVGDEDVRLVGVEAAGLGLDSGKHSATLAFGDVGVYHGSMSY
LLQDDQGQILKPHSVGVGLEYPGVGPEISFMKETGRAEFYTATDEEAIQACMRLSRLEGIIPALEASHALAF
LDKLVPTLRDGAKVVVNCSGRGDKDLDTLIQRGMPSSFC
>ARABIDOPSIS_THALIANA_AT5G54810_TSB_1
MAASGTSATFRASVSSAPSSSSQLTHLKSPFKAVKYTPLPSSRSKSSSFSVSCTIAKDPPVLMAAGSDPALWQ
RPDSFGRFGKFGGKYVPETLMHALSELESAFYALATDDDFQRELAGILKDYVGRESPLYFAERLTEHYRRE
NGEGPLIYLKREDLNHTGAHKINNAVAQALLAKRLGKKRIIAETGAGQHGVATATVCARFGLECIIYMGAQ
DMERQALNVFRMRLLGAEVRGVHSGTATLKDATSEAIRDWVTNVETTHYILGSVAGPHPYPMMVRDFHA
VIGKETRKQALEKWGGKPDVLVACVGGGSNAMGLFHEFVNDTEVRMIGVEAAGFGLDSGKHAATLTKG
DVGVLHGAMSYLLQDDDGQIIEPHSISAGLDYPGVGPEHSFFKDMGRAEYYSITDEEALEAFKRVSRLEGII
PALETSHALAYLEKLCPTLSDGTRVVLNFSGRGDKDVQTVAKYLDV
>BRASSICA_RAPA_XM_009112467
MSSTKIQIRGQPLFKVLTRNHRMINSVVCGVPIKRQHRVSNVLRTSDPPLGSVPTRTDESQFLRGDGNGRFG
RFGGKFVPETLMSPLRDLEDEFDFVLNDHEFQEELTTALRDYVGRETPLYFAGRLTEHYKNISQTTGGGPEI
YLKREDLSHCGSHKINNALGQAMIARRLGCKRVVAATGAGQHGVATAAACAKLSIECTVFMGTTDIEKHS
SNVLSMKLLGAQVKSVEGTFKDASSEAIRNWVGNLETTYYLSGTVVGPHPNPLMVREFQSVIGKETRRQA
KQLWGGKPDVLVACVGSGSNALGLFHEFLGDEDVRLVGVEAAGLGLDSGKHSATLAVGDVGVYHGSMS
YLLQDDQGQILKPHSIGVGLEYPGVGPEISFLKESGRAEFYTATDQEAVQACMLLSRLEGIIPALEASHALAF
LDKLVPTLRDGAKVVVNCSGRGDKDLDTLIQRGMPSSLC
>BRASSICA_RAPA_XM_009139385
MATSGTASTFRPSVSASSRLTHLRSPPSKVPLFTPLPSSRSRSFSVSCTIAKDPTFLMAEAENTKTAGSDPTLW
KRPDSFGRFGKFGGKYVPETLMHALSELETAFYSLATDDDFQRELAGILKDYVGRESPLYFAERLTEHYRR
ENGEGPLIYLKREDLNHTGAHKINNAVAQALLAKRLGKKRIIAETGAGQHGVATATVCARFGLQCIIYMGA
QDMERQALNVFRMRLLGAEVRGVHSGTATLKDATSEAIRDWVTNVETTHYILGSVAGPHPYPMMVRDFH
AVIGKETRRQAMEKWGGKPDVLVACVGGGSNAMGLFHEFVDDTEVRMIGVEAAGFGLDSGKHAATLTK
GDVGVLHGAMSYLLQDDDGQIIEPHSISAGLDYPGVGPEHSFLKDMGRAEYYSVTDEEALEAFKRVSRLEG
IIPALETSHALAHLEKLCPTLPDGARVVLNFSGRGDKDVQTAIKYLEV
>ACTAEA_CIMICIFUGA_CNA0013768_67886
MAISTNASLKTSSASSLSKPSFSSHRKPFNFGKFLLNPSSITPTSISCTLTRASIQPMEVEEVYSTILQRPDSFGR
FGKFGGKYVPETLMYALSELESAFRLLASDEDFQKELGGILKDYVGRESPLYFAERLSEHYKRPNGEGPDV
YLKREDLNHTGAHKINNAVAQALLAKRLGKKRIIAETGAGQHGVATATVCARFGLECIIYMGAQDMERQ
ALNVFRMRLLGAEVRAVHSGTATLKDATSEAIRDWVTNVESTHYILGSVAGPHPYPMMVREFHAVIGKET
RKQAMEKWGGKPDVLVACVGGGSNAIGLFHEFVDDKDVRLIGVEAAGFGLDSGKHAATLTKGEVGVLH
GAMSYLLQDDDGQIIEPHSISAGLDYPGVGPEHSFLKDIGRAEYYSITDEEALEAFQRLSRLEGIIPALETSHA
LGYLEKLCPTLPNGAKVVLNCSGRGDKDVDTVIKHLQV
>BERBERIS_FORTUNEI_CNA0013806_133709
MAISASVSCRNPNPQTPKTHFSSSNYLSIPSRILNFPSCPSTMNHASISCSASTNSVLQVAEEETKSTVSQRPDS
FGRFGKYGGKYVPETLMYALTELESAFHLLAKDEGFQRELDGILRDYVGRESPLYFAERLTEHYKRPNGK
GPLVYLKREDLNHTGAHKINNAVAQALLAKRLGKKRIIAETGAGQHGVATATVCARFGLECIIYMGAQDM
ERQSLNVFRMRLLGAEVRAVHSGTATLKDATSEAIRDWVTNVESTHYILGSVAGPHPYPMMVREFHAVIG
KETRKQALEKWGGKPDVLVACVGGGSNAMGLFHEFVDDEDVRLIGVEAAGFGLDSGKHAATLTKGEIGV
LHGAMSYLLQDDDGQIIEPHSISAGLDYPGVGPEHSYLKDIGRAEYYCVTDQEALEAFKRLSRLEGIIPALET
SHALAYLEKLCPTLPNGTKVVLNCSGRGDKDVHTAIKHLQV
>COPTIS_CHINENSIS_CNA0013793_176826
MAFSTTSISSIKTPSNFHKLISSPSAIKQPSSSMPCTLTTKVASPLQIEVNEKNTNSTILQRPDSFGRFGKFGGK
YVPETLMHALTELESAFRLLASDEAFQKELDGILKDYVGRETPLYFAERLTEHYKGPSGEGPDIYLKREDLN
HTGAHKINNAVAQALLAKRLGKKRIIAETGAGQHGVATATVCARFGLECIIYMGAQDMERQSLNVFRMRL
LGAEVRAVHSGTATLKDATSEAIRDWVTNVESTHYILGSVAGPHPYPMMVREFHAVIGKETRKQAIEKWG
GKPDVLVACVGGGSNAMGLFHEFVDDSDVRLIGVEAAGHGVDSGRHAATLTKGEVGVLHGAMSYLLQD
DDGQIIEPHSISAGLDYPGVGPEHSYLKDIGRAEYYSITDDEALEAFKRLSQLEGIIPALETSHALAYLEKLCP
TLPNGTKVVLNCSGRGDKDVQTAIKYLGADL
>CLEMATIS_MONTANA_CNA0013865_8094
MAISATTATYKTPASPSFLPKPSIPSPNSIQRLSLKPTLTRVSSSLKMEAKETNSDSNSTVLQRPDSFGRFGKF
GGKYVPETLMYALTELENAFRLLANDRDFQEELAGILRDYVGRESPLYFAERLTEHYKRANGEGPHVYLK

REDLNHTGAHKINNAVAQALLAKRLGKKRIIAETGAGQHGVATATVCARFGLECIIYMGAQDMERQSLNV
FRMRLLGAEVRPVHSGTATLKDATSEAIRDWVTNVESTHYILGSVAGPHPYPMMVREFHAVIGKETRKQA
MEKWGGMPDIIVACVGGGSNAMGIFHEFVDDKEVRLIGVEAAGFGLDSGKHAATLTKGEVGVLHGAMSY
LLQDDDGQIIEPHSISAGLDYPGVGPEHSFLKDLGRAEYYSITDTEALEAFKRLAQLEGIIPALETSHALGYLE
KLCPTLPNGTKVVLNCSGRGDKDVQTVIKHMQV
>PAPAVER_SOMNIFERUM_XM_026566589
MAFSSSTVTCKNPNSNLLQKPYLSSSSSFNPLRVHFQKFPLPSSSSSIKSSSISCAVTEEKSKMSTISQRPDSFG
RFGKFGGKYVPETLMYALTELESAFRSLAADQDFQEELSGIFKDYVGRETPLYFAERLTEHYKSANGGGPH
IYLKREDLNHTGAHKINNAVAQALLAKRLGKKRIIAETGAGQHGVATATVCARFGLECIIYMGAMDMERQ
ALNVFRMRLLGAEVRPVHSGTATLKDATSEAIRDWVTNVESTHYILGSVAGPHPYPMMVREFHAVIGKET
RKQAMEKWGGMPDVLVACVGGGSNAMGLFHEFVDDKEVRLIGVEAAGFGIDSGKHAATLTKGEVGVLH
GAMSYLLQDDDGQIIEPHSISAGLDYPGVGPEHSFLKDLGRAEYYSITDEEALEAFKRVSRLEGIIPALETSH
ALAYLEKLCPTLPDGTKVVVNFSGRGDKDVHTAIKHLKL
>ZEA_MAYS_NP_001384061
MATTAAAAIRSPTRAAAPGPAAPAQQRAVLRVASSRTSSARPRRAAAAVAAAAMQPAKAVAAEAASPAA
VEMGNGAAAPGLQRPDAMGRFGRFGGKYVPETLMHALTELESAFHALATDDEFQKELDGILKDYVGRES
PLYFAERLTEHYKRADGTGPLIYLKREDLNHTGAHKINNAVAQALLAKRLGKQRIIAETGAGQHGVATAT
VCARFGLQCIIYMGAQDMERQALNVFRMRLLGAEVRAVHSGTATLKDATSEAIRDWVTNVETTHYILGSV
AGPHPYPMMVREFHKVIGKETRRQAMDKWGGKPDVLVACVGGGSNAMGLFHEFVEDQDVRLVGVEAA
GHGVDTDKHAATLTKGQVGVLHGSMSYLLQDDDGQVIEPHSISAGLDYPGVGPEHSFLKDIGRAEYDSVT
DQEALDAFKRVSRLEGIIPALETSHALAYLEKLCPTLADGVRVVVNCSGRGDKDVHTASKYLDV
>SORGHUM_BICOLOR_XM_002443775.2
MAAATIRSPTRAAAAGGPAAAPTQQRSVLRVASSAASASSSSARTRRGAAAMQPAKAVAAEAASPAVEM
GNGAAVAGMQRPDAMGRFGKFGGKYVPETLMHALTELENAFHALATDEEFQKELDGILKDYVGRESPLY
FAERLTEHYKRADGTGPLIYLKREDLNHTGAHKINNAVAQALLAKRLGKQRIIAETGAGQHGVATATVCA
RFGLQCIIYMGAQDMERQALNVFRMRLLGAEVRAVHSGTATLKDATSEAIRDWVTNVETTHYILGSVAGP
HPYPMMVREFHKVIGKETRRQAMDKWGGKPDVLVACVGGGSNAMGLFHEFVEDQDVRLIGVEAAGHGV
DTDKHAATLTKGEVGVLHGSMSYLLQDDDGQVIEPHSISAGLDYPGVGPEHSFLKDIGRAEYDSVTDQEAL
DAFKRVSRLEGIIPALETSHALAYLEKLCPTLPDGVRVVVNCSGRGDKDVHTASKYLDV
>FRAXINUS_CHINENSIS_CNA0013754_10043
MAFSSAAPCRTQTYLPSSADSHSLLPLKVDKITSPCCGLNPKPHSICCTLAKQQPMTMTEQQLEPVVMQRP
DSFGRFGQFGGKYVPETLMYALTELETAFKSLTFDQEFQKELDGILRDYVGRESPLYFAERLTEHYKRPDG
KGPHIYLKREDLNHTGAHKINNAVAQALLAKRLGKKRIIAETGAGQHGVATATVCARFGLQCIIYMGAQD
MERQALNVFRMRLLGAEVRPVHSGTATLKDATSEAIRDWVTNVESTHYILGSVAGPHPYPMMVREFHSVI
GKETRKQALEKWGGKPDVLVACVGGGSNAIGLFHEFVDDKNVRLIGVEAAGFGLDSGKHAATLSKGEVG
VLHGAMSYLLQDDDGQIIEPHSISAGLDYPGVGPEHSFLKDIGRAEYYSITDEEALEAFKRLSRLEGIIPALET
SHALAYLEKLCPMLPDGTKVVLNCSGRGDKDVQTAIKQLKL
>FRAXINUS_CHINENSIS_CNA0013754_87781
MACNNLKAIVINGVNFRATSESLNSSKRLGFAMHSQTDRGLMVCSAIGHNTLNSDESRLSNPTRSSDILRLI
DQCEKTLESSGKFGRFGGVFVPETLITCLKKLEAEFKWVLHDPEFQMELSIALKDYVGRETPLYFADRLTSY
YKNENGEGPKIYLKREDLTHGGAHKINNAIAQVMLAKRMGRKIIVAATGAGQHGVATAAACAKLGLECTI
FMGKIDMERQSSNVLLMKHLGAKVKSVAGTFKDATSEAIRNWVGNLEGSYYLAGTAVGPHPCPSMVREF
QTVIGKETRKQAMEKWGGEPDVLVACVGSGSNALGLFHEFVRDENVRLIGVEAAGNGIDSGKHSATLAK
GELGVYHGTMSYLLQDDEGQIIGPHSVGVGLEYPGVSPELSFLKDIGRAEFYTVTDEEALEAYTRLCRLEGI
FPALEAAHALAYLEKLCPDLANGTKVVVNCSGRGDKDAASVFKYQEDKIKRGHKESN
>CATHARANTHUS_ROSEUS_UOYN_SCAFFOLD_2042758
MAFSSSTAVRPNTTAAPPQQLPKPYRPSSAAALFFQKPTKFNQFSSSKAASISCVLTKEAMVTKEDLEPVVL
QRPDSFGRFGKFGGKYVPETLMYALSELEDAFKSLSTDREFQEELDGILKDYVGRESPLYFAERLTEYYRRP
DGEGPLIFLKREDLNHTGAHKINNAVGQALLAKRLGKKRIIAETGAGQHGVATATVCARFGLECIIYMGAQ
DMERQALNVFRMRLLGAEVRPVHSGTATLKDATSEAIRDWVTNVESTHYILGSVAGPHPYPMMVREFHA
VIGKETRKQALEKWGGKPDVLVACVGGGSNAMGLFHEFVDDKDVRLVGVEAAGFGLDSGKHAATLTKG
EVGVLHGAMSYLLQDDDGQIVEPHSISAGLDYPGVGPEHSFLKDIGRAEYYSVTDEEALEAFKRASRLEGII
PALETSHALAYLEKLCPTLPDGTKVVLNFSGRGDKDVQTAIKHLQL
>CATHARANTHUS_ROSEUS_UOYN_SCAFFOLD_2004866
MACSSNSLRFCLTRGGGEVKFSTESNFAAAVSAVPSLCNKRASLISCSAVNHQEISSVVSAADSLISSQLEAL
TRSLSQSSQQRLNNLQQIEKEISSEQVYDVHTNNGKFGRFGGKFVPETLITSLNMLEAEFNFILNNADFQKEL

EITLRDYAGRETPLYFAERLTDYYKNSEGEGPEIYLKREDLVHGGSHKMNNAIAQAMLAKRIGRNSIITATG
AGQHGVATAAACAKLSLECTVFMATADIERQPSNVLLMKHLGAQVKVVDGSFKDAISEAIRNWVGELESS
YFLAGTAVGPHPCPTMVREFQSVIGKEIRKQANEKWGGKPDVLVACVGSGSNALGMFHEFIRDEDVRLIG
VEAAGTGLDEGLHSATLARGEVGVYHGTMSYLLQDDEGQILVPHSIGVGLEYPAVSPELSFLKETGRAEFY
SATDEEALDAYERLCRLEGIFPALETCHALAYLEKLCPTLKDGTKVVVNCSGRGDKDAAIVFNRTR
>NANDINA_DOMESTICA_YHFG_SCAFFOLD_2008512
MATSLTTSWRNPNLQISKPYRTTSNSLGLPFEFGKFPLYPSSLKPSSISCSVTTSSALQMEEKDTKSTISQRPD
SFGRFGKYGGKYVPETLMYALTELESAFHLLAKDEAFQKELAGILKDYVGRETPLYFAERLTEHYKHPNGE
GPEVYLKREDLNHTGAHKINNAIAQALLAKRLGKKRIIAETGAGQHGVATATVCARFGLECIIYMGAQDM
ERQALNVFRMRLLGAEVRAVHSGTATLKDATSEAIRDWVTNVESTHYILGSVAGPHPYPMMVREFHAVIG
KETRKQAMEKWGRKPDVLVACVGGGSNAMGLFHEFVDDEDVRLIGVEAAGFGLDSGKHAATLTKGEIG
VLHGAMSYLLQDDDGQIIEPHSISAGLDYPGVGPEHSFLKDIGRAEYYSITDEEALEAFKRLSRLEGIIPALET
SHALAYLEKLCPTLPNGTKVVLNCSGRGDKDVHTAIKHLQV
>CAMELLIA_SINENSIS_XM_028268887
MATVSSPNPSCRTIHISTSLPKPYPSFFNPFILPKFCVSPRSSSISCTFTRDSPTRPMELQDQSLVRPRLGVLQRP
DSFGRFGKFGGKYVPETLMYALTELESAFNALAADQDFQKELDVILKDYVGRESPLYFAKRLTEHYKRPN
GEGPHIYLKREDLNHTGAHKINNAVAQALLAKRLGKKRIIAETGAGQHGVATATVCAQFGLECVIYMGAQ
DMERQALNVFRMQLLGAEVIPVHSGTATLKDATSEAIRDWVTNVESTHYILGSVAGPHPYPMMVREFHAV
IGKETRKQALEKWGGKPDVLVACVGGGSNAMGLFHEFVNDEEVRLIGVEAAGFGLDSGKHAATLTKGEV
GVLHGAMSYLLQDEDGQIVEPHSISAGLDYPGVGPEHSFLKDLGRAEYYSITDEEALEGFKRLSRLEGIIPAL
ETSHAIAYLEKLCPTLPNGTKVVLNCSGRGDKDVQTAIKYLQI
>CAMELLIA_SINENSIS_XM_028209702.1
MIMASSPRNVFLQGGNTTEFSEPVLKLRRLPFSAPLGSPYIKAADRPRAPISCSAVADKMPATVGGALDSNS
LTTRRLMTQTFIDQAELMTVSSTGKFGRFGGKFVPETLISCLSKLEAEFILALHDENFQEELATALRDYVGR
ESPLYFAKRLTDYYKNSEGKGPEIYLKREDLVHTGAHKINNAIAQAMLAKRIGRKRIVAATGAGQHGVAT
AAACAKLSLECTIFMGSADVGRQGANVLLIKNLGAQVKVVDGSFKDAMSEAIRNWVGDLETSYFLSGTAV
GPHPCPTIVREFQSVIGKETRKQAEEKWGGKPDVLVACVGSGSNALGMFHEFIRDEDVRLIGVEAAGVGLD
SGAHSATLARGDVGVYHGSMSYLLQDEEGQIIGPHSIGVGLEYPGVSPELSFLKDNGRAEFYAVTDDEALD
AYRRLCQLEGIFPALEASHALAYLEKLCPGLEDGAKVVVNCSGRGDKDAATVYKHNEERV
>RHEUM_LACINIATUM_CNA0013760_28769
MAVSTGSALKAQSSSPGYSTAHNTAFNLSSLPSNSLNFSQYSLSRRPTSISCIMTRDELGAKNLKGEMSDHE
TGLLQRPDSFGRYGKFGGKYVPETLMYALTELESAFHALSKDPLFQSELDGILRDYVGRESPLYFAERLTEH
YKRSNGEGPHIYLKREDLNHTGAHKINNAVAQALLAKKLGKKRIIAETGAGQHGVATATVCARFGLECIV
YMGAQDMERQSLNVFRMRLLGAEVRPVHSGTATLKDATSEAIRDWVTNVETTHYILGSVAGPHPYPMMV
RDFHAVIGKETRKQAQDKWGGIPDVLVACVGGGSNAMGLFHEFVDDKEVRLIGVEAAGFGLDSGKHAAT
LTKGEVGVLHGAMSYLLQDDDGQVIEPHSISAGLDYPGVGPEHSFLKDIGRAEYYSITDEEALEAFKRLSRL
EGIIPALETSHALAYLEKLCPTLSDGTRVVLNCSGRGDKDVHTAIKYLKM
>RHEUM_LACINIATUM_CNA0013760_22600
MSSTNILRNRFFAGNGVGTTTQSCTICRKTDVCLPSRNRSSVATMARIRKEMKTITCDTEVEEEEMVKLSGK
YGKFGGKFVPETLINWLTVLEAEFKLALHDPQFQEELATALRDYVGRETPLYYAERLTDYYREKNKGVGP
EIYLKREDLCHGGAHKINNAIAQVMLAKRLGKKKIIAATGAGQHGVATAAACAKHSLECVIFMGNSDMER
QSSNVHLIELLGAQVKRINGSFKDAASDAFRERIADLDSNFLLVGTVVGPHPCPTMVQEFQSVIGKEVRRQA
MGKWGGKPDVLVACVGSGSNALGLFHDFIGDEDVRLIGVEAAGSATLTRGSVGVYHGAMSYLLQDEEGQ
IVGPHSIGVGLEYPGVSPELSYLMEMERAEFHAVTDEEALHAFNRLCKLEGIYPALESAHALAYLEKLCPT
MADGAKVVVNISGRGDKDAATVFQCMSI
>FALLOPIA_MULTIFLORA_CNA0013822_104185
MVKTVACNLNNSSLGGANAIGLMFPQSLALSRKMKIGFRTQNLKGLAVVAMARQDVKAMATSSAVLVK
APNPMTETSALIAVPLKEKKQNKEEERKERVSINRESGKYGKFGGKFVPETLMTWLTILEAEFNLALHDSHF
QEELSTALRDYVGRETPLYHAQRLTSYYKDKNGGLGPEIYLKREDLCHGGAHKINNAIGQAMLAKRLGKT
KIVVATGSGQHGVAVAAASAKLDLECTVFMGTSIMERQSSNVALMNLLGAQVVPVNGSFKDATSDAFRES
LTDFNNNYYLQGSGIGPHPYPMMVQEFQSVIGKEVRKQSMERWGGKPDILLACVGGGSNALGLFHEFVQD
ENVRLIGVEAAGHGLETGFHSATLTKGSVGVYHGAMSYLLQDDEGQILEPHSIGVGLECPGVSPELSYLKD
TGRAEFHAITDIEAVNAYKRLCKLEGIFPALESAHALAYLEELCPTLPNGTKVVVNLSGRGDKDATTVFQC
NLMEQWTSII
>FALLOPIA_MULTIFLORA_CNA0013822_127398

MAVSSGSALKAQTSSSGYSIAQRGYNNAFNLSALSSNSLKFSRNSLSRRPRSISCIMTREELGAKDLKGEMS
DRESGLLQRPDSFGRFGKFGGKYVPETLMHALTELESAFHALAKDPQFQSELDGILRDYVGRESPLYFAER
LTEHYKRSNGEGPHIYLKREDLNHTGAHKINNAVAQALLAKRLGKTRIIAETGAGQHGVATATVCARFGL
QCIVYMGAQDMERQALNVFRMRLLGAEVRPVHSGTATLKDATSEAIRDWVTNVETTHYILGSVAGPHPYP
MMVRDFHAVIGKETRKQAQEKWGRKPDVLVACVGGGSNAMGLFHEFVDDKEVRLIGVEAAGFGLDSGK
HAATLTKGEVGVLHGAMSYLLQDDDGQVIEPHSISAGLDYPGVGPEHSFLKDIGRAEYYSVTDEEALEAFK
RLSRLEGIIPALETSHALGHLEKLCPTLADGTKVVLNCSGRGDKDVHTAIKHLKM
>LAMIUM_GALEOBDOLON_CONTIG_9143
MAESCFLSSSLKPRLSLRGDDQWLSCFPPNIRVCNLRNLPNSGVSKSVSCTADPKPIGIPRQWYNLIADLPVL
PPPPLHPKTFVPIKPEDLSPLFPDELIKQEGSTERFIDIPEEVIDVYRLWRPTPLIRAKRLEKLLDTPARIYYKYE
GGSPAGSHKPNTAVPQVWYNAQEGVKNVVTETGAGQWGSSLAFACSLFDLNCEVWQVRASYDQKPYRR
MMMETWGAKVHPSPSTVTEYGRRILEKYPSSPGSLGIAISEAVEVAAMNADTKYCLGSVLNHVLLHQTVI
GEECIKQMEAIGETPDVIIGCTGGGSNFGGLVFPFIREKLSGKINPIIRAVEPTACPSLTKGVYAYDYGDTAG
MTPLMKMHTLGHDFIPDPIHAGGLRYHGMAPLISHVYELGFMEAISIAQTECFEGAIQFARTEGLIPAPEPTH
AIAATIREARRCRETGEAKVILMAMCGHGHFDLPAYEKYLKGDLVDLSFSEQRIKESLANIPQL
>APHELANDRA_SQUAROSA_AS_145
MAQSVFLTPSANARCSIQGYKHQLGYFASKEKPCHLKLVPKSSLSPTAAQIFNSRAIEIPRQWYNLVADLPV
KPPPPLHPKTFEPVKPDDLTPLFPDELIKQEATLERFIDIPDEVVDVYRLWRPTPLIRAKRLEKLLDTPARIYY
KYEGGSPAGSHKPNSAVPQAWYNAQQGVKNVVTETGAGQWGSSLAFACSLFGLNCEVWQVRASYDQKP
YRKLMMQTWGAKVHPSPSDITEAGRRILEHDASSPGSLGIAISEAVEVAAANPDTKYCLGSVLNHVLLHQT
VIGEECIKQMEAIGETPDVIIGCTGGGSNFGGLAFPFIREKLNGKINPVIRAVEPEACPSLTKGVYAYDYGDT
AGMTPLLKMHTLGHDFIPDPIHSGGLRYHGMAPLISHVYELGFMETMAIAQTECFEGAIKFARTEGIIPAPEP
THAIAATIREACRCRETGESKVILTAMCGHGHFDLPAYEKYLQGDLVDLSFSEESIKASLAKIPRT
>CONSOLIDA_ORIENTALIS_TRINITY_DN14752_C0_G1_I2
MADSLVTKTPPRFSLPHRVNNRWIGGYSPKTKLMHVKCSFSRQVRAGATSRTNTKAIEVPCQWYNLVADL
AIKPPPFLHPMTKEAVKPEDLSPLFSDELISQEFSNERFIDIPDEVVDVYNLWRPTPLIRAKRLEKLLDTPARI
YYKYEGGSPAGSHKPNTAVPQVWYNAQQGVKNIVTETGAGQWGSALAFACSLFDVNCEVWQVRASYDQ
KPYRKLMMQTWGAKVHPSPSDLTNAGRRILETDPSSPGSLGIAISEAVESAVKNGDTKYCLGSVLNHVLLH
QTIIGEECIKQMEEIGETPDVIIGCTGGGSNFGGLAFPFIREKLSGKINPLIRAVEPTACPSLTKGVYAYDYGD
TAGMTPLMKMHTLGHDFIPDPIHSGGLRYHGMSPLISHIYELGFMEALAIPQIECFRGAIQFARTEGLIPAPEP
THAIAATIREALHCKETGEAKVILMAMCGHGHFDLPAYEKYLQGNMVDLSFSDEKMQASLADLPNIS
>NICOTIANA_TABACUM_XM_016610964
MALSPPNLYAKGDAYGIKYFEIKTKPSQLKLCFSCRARAKAALSTRSSSIEVPRQWYNLVADLPIKPPPPLHP
KTFQPIKPEDLSPLFCDELIKQEASIDQFIDIPEEVLDVYSLWRPTPLIRAKRLEKLLDTPARIYYKYEGGSPAG
SHKPNTAVPQAWYNKMGSVKNVVTETGAGQWGSALSFACSLFGLNCEVWQVRASFDQKPYRKMMMQT
WGAKVHPSPSDLTEAGRTILRMDPSSPGSLGIAISEAVEIAATNADTKYCLGSVLNHVLLHQTVIGEECIKQ
MEDFGETPDVIIGCTGGGSNFAGLAFPFIREKLKGKINPLIRAVEPAACPSLTKGVYAYDYGDTAGMTPLMK
MHTLGHDFIPDPIHAGGLRYHGMAPLISHVYELGFMEAISIPQTECFKGAIQFARSEGLIPAPEPTHAIAATIR
EALRCKERGESKVILMAMCGHGHFDLSSYDKYLQGSLVDLSFSEEKIKASLAKIPQPMS
>ORYZA_SATIVA_NM_001422904
MAAAVGNPNAAAAASISASRVGAGALRAGGLRVAAGRRGAGAVVAAAMRPAKAVASPAKEAAGEVNG
AASGGFARPDAFGRFGKFGGKYVPETLMHALTELEAAFHALAGDEDFQKELDGILKDYVGRETPLYFAER
LTEHYKRADGTGPMIYLKREDLNHTGAHKINNAVAQVLLAKRLGKERIIAETGAGQHGVATATVCARFGL
QCIIYMGAQDMERQALNVFRMKLLGAEVRAVHSGTATLKDATSEAIRDWVTNVENTHYILGSVAGPHPYP
MMVREFHKVIGKETRRQAMEKWGGKPDVLVACVGGGSNAMGLFHEFVDDQDIRMIGVEAAGYGVDTD
KHAATLTKGEVGVLHGSLSYVLQDDDGQVIEPHSISAGLDYPGVGPEHSFLKDIGRAEYDSVTDQEALDAF
KRVSRLEGIIPALETSHALAYLEKLCPTLPDGVRVVVNCSGRGDKDVHTASKYLDV
>HORDEUM_VULGARE_XM_045100595
MAASAIRNPSPAAAVSAPPPSRAVLRMLTPSARRGASVVASASMRPVKAVAAEAPSPVSERVNGAEVAGA
GIARPDALGRFGKFGGKYVPETLMHALTELEAAFHALADDEDFQKELDGILKDYVGRESPLYFAERLTEHY
KRADGTGPLIYLKREDLNHTGAHKINNAVAQALLAKKLGKKRIIAETGAGQHGVATATVCARFGLECIIYM
GAQDMERQALNVFRMKLLGAEVRPVHSGTATLKDATSEAIRDWVTNVETTHYILGSVAGPHPYPMMVRE
FHKVIGKETRRQAMDKWGGKPDVLVACIGGGSNAMGLFHEFVDDQDVRLIGVEAAGHGVDTDKHAATL
TKGEVGVLHGSLSYVLQDADGQVIEPHSISAGLDYPGVGPEHSFLRDIGRAEYDSVTDQEALDAFKRTSRL
EGIIPALETSHALAYLEKLCPTLPDGVRVVLNCSGRGDKDVHTASKYLDV
>BRACHYPODIUM_DISTACHYON_XM_003573303

MAASAIRNPSPASAAAVAASRSAVSTSRSILRVPAASRRSLVVAAGMRPAKAVAAEAPTPVAERVNGAEV
ARPDAMGRFGKFGGKYVPETLMHALTELEAAFHALADDQDFQTELDGILKDYVGRESPLYFAERLTEHYK
RADGTGPLIYLKREDLNHTGAHKINNAVAQALLAKRLGKQRIIAETGAGQHGVATATVCARFGLQCVIYM
GAQDMERQALNVFRMRLLGAEVRAVHAGTATLKDATSEAIRDWVTNVETTHYILGSVAGPHPYPMMVR
EFHKVIGKETRRQAMDKWGGKPDVLVACVGGGSNAMGLFHEFVDDQDVRLIGVEAAGHGVDTDKHAAT
LTKGEVGVLHGSLSYVLQDADGQVIEPHSISAGLDYPGVGPEHSFLRDIGRAEYDSVTDQEALDAFKRTSR
LEGIIPALETSHALAYLEKLCPTLPDGVRVVLNCSGRGDKDVHTASKYLDV
>TRITICUM_AESTIVUM_XM_044575364
MAASAIRNPSPAAAVAAPLPSRAVLRMVTPSARRGASVVASASMRPAKAVAAEAPSPVAERVNGAEVAG
AGIARPDALGRFGKFGGKYVPETLMHALTELEAAFHALADDEDFQKELDGILKDYVGRESPLYFAERLTEH
YKRADGTGPLIYLKREDLNHTGAHKINNAVAQALLAKKLGKQRIIAETGAGQHGVATATVCARFGLQCIIY
MGAQDMERQALNVFRMKLLGAEVRPVHSGTATLKDATSEAIRDWVTNVETTHYILGSVAGPHPYPMMV
REFHKVIGKETRRQAMDKWGGKPDVLVACVGGGSNAMGLFHEFVDDQDVRLIGVEAAGHGVDTDKHAA
TLTKGEVGVLHGSLSYVLQDADGQVIEPHSISAGLDYPGVGPEHSFLRDIGRAEYDSVTDQEALDAFKRTS
RLEGIIPALETSHALAYLEKLCPTLPDGVRVVLNCSGRGDKDVHTASKYLEV
>TRITICUM_AESTIVUM_XM_044567734
MATALRPPRLPAVPEQASSLHRLPKHRVAVTGRRSFAARAGSYPGNVGVPKQWYNLIADLPVKPPPMLHP
GTHQPLNPSDLAPLFPDELIRQELTEERFIDIPDEVRDVYELWRPTPLIRAKRLEKLLGTPAKIYYKYEGTSPA
GSHKGNTAVPQAWYNAAAGVKNVVTETGAGQWGSALSFASTLFGLNCEVWQVRASYDQKPYRRLMME
TWGAKVHPSPSDVTEAGRRLLAADPSSPGSLGMAISEAVEVAATNADTKYCLGSVLNHVLLHQTVIGEEC
LEQLAAIGDTPDVVIGCTGGGSNFGGLAFPFMREKLAGRMNPQFRAVEPAACPTLTKGVYAYDYGDTAGL
TPLMKMHTLGHDFVPDPIHAGGLRYHGMAPLISHVYELGFMEAMSIQQTECFEAALQFARTEGIIPAPEPTH
AIAAAIREALECKRTGEEKVILIAMCGHGHFDLAAYDRYLRGDMIDLSHSSEKLKESLGAIPKV
>HORDEUM_VULGARE_XM_045105492
MATASTALRPPRLPAVAEQASPLHRLPKNRVVAVNVRRSFAARAGSYPGNVGVPKQWYNLIADLPVKPPP
MLHPGTHQPLNPSDLAPLFPDELIRQELTEERFIDIPDEVRDVYELWRPTPLIRAKRLEKLLGTPAKIYYKYE
GTSPAGSHKGNTAVPQAWYNAAAGVKNVVTETGAGQWGSALSFASTLFGLNCEVWQVRASYDQKPYRR
LMMETWGAKVHPSPSDVTEAGRRLLAADPSSPGSLGMAISEAVEVAATNADTKYCLGSVLNHVLLHQTVI
GEECLEQLAAIGDTPDVVIGCTGGGSNFGGLAFPFMREKLAGRMNPQFKAVEPAACPTLTKGVYTYDYGD
TAGLTPLMKMHTLGHDFVPDPIHAGGLRYHGMAPLISHVYELGFMEAMSIQQTECFEAAIQFARTEGIIPAP
EPTHAIAAAIREALECKRTGEEKVILIAMCGHGHFDLAAYDRYLRGDMIDLSHSSEKLKESLGAIPKV
>BRACHYPODIUM_DISTACHYON_XM_003563542
MAAASIALHPPRLQGPGKAALPPRIPNRSRVTVSGTRSFATRAGSNPGNVSIPKQWYNLVADLPVKPPPQLH
PQTHQPLKPSDLSPLFPDELIRQELTEERFVDIPEEVRDVYELWRPTPLIRAKRLEKLLGTPAKIYYKYEGTSP
AGSHKANTAVPQAFYNAAAGVKSVVTETGAGQWGSALSFASTLFGLTCEVWQVRASYDQKPYRRLMME
TWGAKVHPSPSEATESGRKLLAADPSSPGSLGMAISEAVEVAATNGDTKYCLGSVLNHVLLHQTVIGEECL
EQLAAIGDTPDVVIGCTGGGSNFGGLAFPFMREKLAGRINPVFKAVEPAACPTLTKGVYAYDYGDTAGLTP
LMKMHTLGHGFVPDPIHAGGLRYHGMAPLISHVYELGFMEAMSIQQTECFEAALQFARTEGIIPAPEPTHAI
AAAIREALECKRTGEEKVILIAMCGHGHFDLAAYDRYLRGDMVDLSHSSEKLKESLAAIPKV
>ORYZA_SATIVA_XM_015788505
MATTASVRPPLLRQAAGSEKASLLCKPKQRASVRRRSFTARASSNPVSIPKQWYNLVADLPVKPPPPLHPQ
THQPLNPSDLSPLFPDELIRQEVTEERFIDIPEEVAEVYKLWRPTPLIRARRLEKLLGTPAKIYYKYEGTSPAG
SHKPNTAVPQAWYNAAAGVRSVVTETGAGQWGSALSFASSLFGLTCEVWQVRASYDQKPYRRLMMETW
GATVHPSPSAATESGRRILERDPASPGSLGIAISEAVEVAARDADTKYCLGSVLNHVLLHQTVIGEECLEQL
AAAGDVPDVVIGCTGGGSNFGGLVFPFMREKLAGRMSPAFKAVEPAACPTLTKGVYAYDFGDTAGLTPL
MKMHTLGHGFVPDPIHAGGLRYHGMAPLISHVYELGFMEAIAIQQTECFDAALKFARTEGIIPAPEPTHAIA
AAIREAMECKRTGEKKVILMAMCGHGHFDLASYEKYLRGDMVDLSHSDEKLQEALAAVPKI
>SORGHUM_BICOLOR_XM_021448695
MAAAAALRPALSQAAGPEHRASLLCTPKHRVSASASRRSLRFTARASSNPGAQVSIPKQWYNLIADLPVKP
PPPLHPQTHQPLNPSDLSPLFPDELIRQEVTDERFVDIPEEVIDVYKLWRPTPLIRARRLEKLLGTPAKIYYKY
EGTSPAGSHKPNTAVPQAWYNAAAGVKNVVTETGAGQWGSALSFASSLFGLNCEVWQVRASFDQKPYR
RLMMETWGAKVHPSPSTATEAGKRILEADPSSPGSLGIAISEAVEVAATNADTKYCLGSVLNHVLLHQTVI
GEECLEQLAALGETPDVVIGCTGGGSNFGGLAFPFLREKLRGNMSPAFRAVEPAACPTLTKGVYAYDFGDT
AGLTPLMKMHTLGHGFVPDPIHAGGLRYHGMAPLISHVYELGFMDAIAIQQTECFQAALQFARTEGIIPAPE
PTHAIAAAIREALECKRTGEEKVILMAMCGHGHFDLAAYEKYLRGDMVDLSHPAEKLEASLAAVPKV
>CONSOLIDA_ORIENTALIS_BX1

MALAITSSAFSLVCQKPAVIQKSSETRGSLTISPSSLTISPSSVSISETFASLRQQGKVALVPYITAGDPDLSTT
AEALKVLDYCGSDVIELGVPCTDPFLDGPVIQAACKRSLGGGANMKSIFSMLQKVSPQLSCPILLFTYYKQI
LKCGIGRFMAATNDAGVRGLLVPDAPLEHTEVLRAEASKYGIEIVLLTTPITPKERMKKIVQVAQGFVYLVS
SVGVTGARPSVNPRVQSLLQEXKEVTNKPVVVGFGISKPEHVKQIARWGADGVIVGSAMVKLLGEAKTAN
EGLKELEAFTMSLKTALSENNSLLMI
>ZEA_MAYS_BX1
MAFAPKTSSSSSLSSALQAAQSPPLLLRRMSSTATPRRRYDAAVVVTTTTTARAAAAAVTVPAAPPQAPAP
APVPPKQAAAPAERRSRPVSDTMAALMAKGKTAFIPYITAGDPDLATTAEALRLLDGCGADVIELGVPCSD
PYIDGPIIQASVARALASGTTMDAVLEMLREVTPELSCPVVLLSYYKPIMSRSLAEMKEAGVHGLIVPDLPY
VAAHSLWSEAKNNNLELVLLTTPAIPEDRMKEITKASEGFVYLVSVNGVTGPRANVNPRVESLIQEVKKVT
NKPVAVGFGISKPEHVKQIAQWGADGVIIGSAMVRQLGEAASPKQGLRRLEEYARGMKNALP
>CARICA PAPAYA_XM_022031690
MICANIPLSYLPKSPKIFKCPLICSAMTEETILTREAKPIIQKLPEISVRSVPSTPGKFGKFGGKFVPETLMTSLS
NLEAEFNLVLKDSEFQEELATALRDYVGRETPLYFAERLTDHYKKSKNGEGPEIYLKREDINHTGAHKINN
AIAQAMIAKRMGRKTISCATGAGQHGVATAAACAKLGLECIVFMGTADMEKQASSVTFMKLLGAQVKGV
EGSFKDASSEAIREWVGNLESVYYLTGTVVGAHPCPSMVREFQSVIGKETRRQAMEKWGGKPDVLLACIG
SGSNALGLFHDFIGDEHVRLIGVEAAGFGLDTTKHSATLATGHLGVYHGAMTFLLQDEEGQILPPYSIALGL
EYPGVGPEISFLKDSGRAEFYSVTDQEALNAYVRVCRLEGILPSLEAAHALAFLEKLCPTLPNGTKVVVNCS
GRGDKDAALVLQHIKDSIHQ
>CARICA PAPAYA_XM_022031689
MICANIPLSYLPKSPKIFKCPLICSAMTEETILTREAKPIIQKLPEISVRSVPSTPGKFGKFGGKFVPETLMTSLS
NLEAEFNLVLKDSEFQEELATALRDYVGRETPLYFAERLTDHYKKSKNGEGPEIYLKREDINHTGAHKINN
AIAQAMIAKRMGRKTISCATGAGQHGVATAAACAKLGLECIVFMGTADMEKQASSVTFMKLLGAQVDTC
MHRLFLSQIYGFNYLFKLWLQVKGVEGSFKDASSEAIREWVGNLESVYYLTGTVVGAHPCPSMVREFQSVI
GKETRRQAMEKWGGKPDVLLACIGSGSNALGLFHDFIGDEHVRLIGVEAAGFGLDTTKHSATLATGHLGV
YHGAMTFLLQDEEGQILPPYSIALGLEYPGVGPEISFLKDSGRAEFYSVTDQEALNAYVRVCRLEGILPSLEA
AHALAFLEKLCPTLPNGTKVVVNCSGRGDKDAALVLQHIKDSIHQ
>CARICA PAPAYA_XM_022037282
MAASTCRPYAYSQRKSLASSSRPSYTSTSSRFTFNFSKFTPRPPSKSPLSLSCTLTRDPAIQMEDPAQWLRPD
SFGRFGKFGGKYVPETLMYALTELESAFHSLSADDVFQRELAGILKDYVGRESPLYFAERLTEHYRRANGE
GPVIYLKREDLNHTGAHKINNAVAQALLAKRLGKKRIIAETGAGQHGVATATVCARFGLECIIYMGAQDM
ERQALNVFRMRLLGAEVRAVHSGTATLKDATSEAIRDWVTNVETTHYILGSVAGPHPYPTMVRQFHAVIG
KETRAQALEKWGGKPDVLVACVGGGSNAMGLFHEFVNDKDIRLIGVEAAGFGLDSGKHAATLTKGEVGV
LHGAMSYLLQDEDGQIIEPHSISAGLDYPGVGPEHSFLKDAGRAEYYSVTDDEALEAFKRLSQLEGIIPALET
SHALAYLETLCPTLPNGTKVVVNCSGRGDKDVQTAIKHLQV
>CARICA PAPAYA_XM_022047687
MSLFSANPLHLQPSSSFISSKADEQQFSCFSLKRNLLYLRSSNGCRLRATAALNSDCKSVIGIPHQWYNLIAD
LPIKPPPPLHPKTFQPIKPEDLSPLFPDELIKQEAANDRYIDIPDEVLDVYRLWRPTPLIRAKRLEKLLGTPARI
YYKYEGVSPAGSHKPNTAVPQAFYNAQQGIKNVVTETGAGQWGSSLAFACCLFGLDCEVSNTMISSDTIH
AGGLRYHGMAPLISHVYELGFMEAIAIPQIECFQGAIQFARSEGIIAAPEPTHTIAATIREALCCKESGEAKVI
LMGVCGHGHLDLPSYDKFLQGKLVDLSFEGKKIQESLAKIPQVVP
>ARABIDOPSIS THALIANA_AT4G27070
MATASTAATFRPSSVSASSELTHLRSPSKLPKFTPLPSARSRSSSSFSVSCTIAKDPAVVMADSEKIKAAGSDP
TMWQRPDSFGRFGKFGGKYVPETLMHALSELETAFYSLATDEDFQRELAEILKDYVGRESPLYFAERLTEH
YRRENGEGPLIYLKREDLNHTGAHKINNAVAQALLAKRLGKKRIIAETGAGQHGVATATVCARFGLQCIIY
MGAQDMERQALNVFRMRLLGAEVRGVHSGTATLKDATSEAIRDWVTNVETTHYILGSVAGPHPYPMMV
RDFHAVIGKETRKQAMEKWGGKPDVLVACVGGGSNAMGLFHEFVDDTEVRMIGVEAAGFGLDSGKHAA
TLTKGDVGVLHGAMSYLLQDDDGQIIEPHSISAGLDYPGVGPEHSFLKDVGRAEYFSVTDEEALEAFKRVS
RLEGIIPALETSHALAHLEKLCPTLPDGARVVLNFSGRGDKDVQTAIKYLEV
>CAMELINA SATIVA_XM_010460261
MSSSTKIQVRGKLLPPARNHRMIHSVVYRVSTKRHHRESSVLSSESCPSSDSVPTKTDKSQFCCGNVDGKFG
RFGGKFVPETLMSRLKDLEEEFNFLLSDHKFQDELTTALRDYVGRETPLYFAGRLTEHYKNIARTIGGGPEI
YLKREDLSHCGSHKINNALAQAMIARRLGCSRVVAATGAGQHGVATAAACAKLSLDCTVYMGAPDIEKQ
FSNVLSMKLLGAQVKSVEGTFKDASSEAIRHWVENLDSTYFLLGTVVGPHPCPIMVREFQSVIGKETRRQA
NQLWGGKPDVLVACVGSGSNALGLFHEFVEDEDVRLVGVEAAGLGLDSGKHSATLAVGHVGVYHGSMS

YLLQDDQGQILRPHSVGVGLEYPGVGPEISFLKETGRAEFYTATDQEAIQACMRLSRLEGIIPALETSHALAF
LDKLIPTLRDGAKVIVNCSGRGDKDLDTLIQRGISFPNC
>CONSOLIDA ORIENTALIS_TRINITY_DN2392_C0_G2_I1
MATPIYKTSSSSSCCCLFLKPSLLPKPSTFDKFPYLTPRGCSSKPSPSISCTIASEVEVQQRPDSFGRFGKFGGK
YVPETLMSALSDLESAFNLLAADHHFQKELDEILKDYVGRESPLYFAERLTEHYKRPNGEGPHVYLKREDL
NHTGAHKINNAIAQALLAKRLGKRRIIAETGAGQHGVATATVCARFGLECIIYMGAQDIERQALNVFRMRL
LGAEVRPVHSGTATLKDATSEAIRDWVTNVESTHYILGSVAGPHPYPMMVRDFHAVIGKETRRQAMDKW
GGKPDVLVACVGGGSNAIGLFHEFVDDEDVRLIGVEAAGFGLDSGKHAATLTKGEVGVLHGAMSYLLQD
DDGQIIEPHSISAGLDYPGVGPEHSFLKDIGRAEYYSITDEEALEAFKRLAQLEGIIPALETSHALAYLEKLCP
TLPNGTKVVLNCSGRGDKDVHTAIKHLQV

**Data S3.**

Sequences used for phylogenetic analysis of Fig S8A, provided in FASTA format.

>NICOTIANA TABACUM_XM_016610964
MALSPPNLYAKGDAYGIKYFEIKTKPSQLKLCFSCRARAKAALSTRSSSIEVPRQWYNLVADLPIKPPPPLHP
KTFQPIKPEDLSPLFCDELIKQEASIDQFIDIPEEVLDVYSLWRPTPLIRAKRLEKLLDTPARIYYKYEGGSPAG
SHKPNTAVPQAWYNKMGSVKNVVTETGAGQWGSALSFACSLFGLNCEVWQVRASFDQKPYRKMMMQT
WGAKVHPSPSDLTEAGRTILRMDPSSPGSLGIAISEAVEIAATNADTKYCLGSVLNHVLLHQTVIGEECIKQ
MEDFGETPDVIIGCTGGGSNFAGLAFPFIREKLKGKINPLIRAVEPAACPSLTKGVYAYDYGDTAGMTPLMK
MHTLGHDFIPDPIHAGGLRYHGMAPLISHVYELGFMEAISIPQTECFKGAIQFARSEGLIPAPEPTHAIAATIR
EALRCKERGESKVILMAMCGHGHFDLSSYDKYLQGSLVDLSFSEEKIKASLAKIPQPMS
>NICOTIANA TABACUM_XM_016633459
MAFSSSTAQTASPLSSKHCCRLSSSAASSASYFPKFQIPFKFDKTTSCPSSISCVLTKQESMAAQEAAEPAVLL
RPDSFGRFGKFGGKYVPETLMHALDELETAFKSLATDEAFQKELDGILRDYVGRESPLYFAERLTEHYKRP
DGEGPLIYLKREDLNHTGAHKINNAVAQALLAKRLGKKRIIAETGAGQHGVATATVCARFGLECIIYMGAQ
DMERQALNVFRMRLLGAEVRGVHSGTATLKDATSEAIRDWVTNVETTHYILGSVAGPHPYPMMVREFHA
VIGKETRKQALEKWGGKPDVLVACVGGGSNAMGLFHEFVDDKDVRLIGVEAAGFGIDSGKHAATLTKGE
VGVLHGAMSYLLQDEDGQIVEPHSISAGLDYPGVGPEHSFLKDLGRAEYYSITDEEALEAFKRLSRLEGIIPA
LETSHALAYLEKLCPTLPNGTKVVLNCSGRGDKDVHTAINYLKV
>NICOTIANA TABACUM_XM_016602914
MACNIEVIVRQAKIAEPRLSYSRKWKGKFAIFSVATGPSRVTELPGKLVYHEKKRAIFSNEKFGIFGGKFVPE
TLISSLTKLDYEFNSALHDPQFQMVLGVALRDYVGRETPLYLAERLTDNYKSRNGGKGPDLYLKREDLNH
VGAHKINNAIAQTMLAKRMDCKNIIAATGAGQHGVATAAACAKLSLECTVFMGSLDMERQPSNVILMKH
LGAKVKSVKGSFKDAVSEGIRHWVNNLETSYFLGGAAIGPHPCPTMVREFQSVIGKETRKQAMEKWGGKP
DVLVACVGSGSNALGLFHEFIEDKDVRLIGVEAGGVGLDTGKHSATMARGQVGVYHGAMSYLLQDDEGQ
IIEPHSIGVGLEYPGVSPELSFLKDTGRAEFYTVTDEQALEAYKRLCRLEGIFPALEASHALAFLDRLCPTLED
GEKVVVNLSGRGDKDAATVFNHTTKNE
>APHELANDRA SQUARROSA_AS_143
MSYSKCLPSNSFLHSNGFYSNSDPKLATRRLNFGRKIAGDKSLTVSSVMTTQDVRTPLNDDQATHSRIRSD
VVRLIEQADEKLSSTGKFGRFGGVFVPETLITCLNKLAAEFNLILHDRGFQAELRTALRDYVGRETPLYYAK
RLSDHYRNGKGEGPDIYLKREDLNHGGAHKINNAIAQAMIAKRMGRMRVVAATGAGQHGVATASACAQ
LGLECTVFMGNVDMERQPSNVLLMKILGAQIKSVEGSFKDATSEAIRHWVGDLDNGYFLTGMAVGPHPLP
TMVREFQAVIGKETRRQAREKWGGKPDVVVACVGSGSNALGIFHEFVKDEDVRLIGVEAAGSGIDTGKHS
ATLSTGDVGVYHGAMSYLLQDDEGQIIGPHSIGVGLEYPGVSPELSFLKDIGRAEFHTVTDEEALDAYAVL
CRLEGIIPALEAAHALAYLGKLCKTLPDGAKVVVNCSGRGDKDAATVFNHQQQKQ
>SOLANUM LYCOPERSICUM_SOLYC10G005320.3.1
MACNINVESILGQGIFATTSSKKLQAFPSHHTYKANVISCVAIGPTPIPQLPWKLVFHEKERQSLLSNEKFGIY
GGKFVPETLISPLTKLDYEFNSALRDPQFQMNLQVALKDYVGRETPLYFAQRLTDYYKSLNKGIGPDIYLK
REDLNHGGAHKINNAIAQAMLAKRMGCKNVVASTGAGQHGVATAAACAKLSLECTIFMGSLDMERQPSN
VLLMNHLGAKVKCVEGSFKDAMSEGIRNWVNNLETSYFLAGAAIGPHPCPTMVREFQSIIGKETRKQAMD
KWGGKPHVLVACVGSGSNALGLFHEFIQDVDVRLIGVEAGGIGLDSGKHSATMARGEVGVYHGAMSYLL
QDEEGQIIGPHSIGVGLEYPGVSPELSYLKDIGRAEFSTVTDEEAIKAYKRLCILEGIFPALESCHALAFLDKL
CSTLKDGEKVIVNLSGRGDKDAEAVFNHTPKHK
>SOLANUM TUBEROSUM_PGSC0003DMT400029363
MACNINVESILGQGIFATTSSKKLQAFPSHNYKPNVISCVAIGPTHKVPQLPWKLVFHEKERPLLSNEKFGIY
GGKFVPETLISPLTKLDYEFNSALRDPQFQMDLQVALKDYVGRETPLYFAQRLTDHYKSLNKGIGPDIYLK
REDLNHGGAHKINNAIAQAMLAKRMGCKNVVASTGAGQHGVATAAACAKLSLECTIFMGTLDMERQPS
NVLLMNHLGAKVKSVEGSFKDAMSEGIRNWVNDLETSYFLAGAAIGPHPCPTMVREFQSIIGKETRKQAM
DKWGGKPHVLVACVGSGSNALGLFHEFIQDEDVRLIGVEAGGTGLDSGKHSATIARGEVGVYHGAMSYL
LQDEEGQIIGPHSIGVGLEYPGVSPELSYLKDIGRAEFSMVTDEEAIKAYKRLCRLEGIFPALESSHALAFLDK
LCSTLKDGEKVIVNLSGRGDKDAEAVFNHTPKHE
>CAPSICUM ANNUUM_CA10G05380
MACCNVETIFGLGNIGRKLQVFPSHNYKASVISCVAVGPSQTPQVPWKLVFHEKERSLLSNEKYGIYGGKF
VPETLISPLTKLDYEFNSALRDPLFQMELEVTLRDYVGRETPLYFAQRLTDYYKSINRGTGPDIYLKREDLN
HGGAHKINNAIAQAMLAKRMGCKSVMAATAAACAKLSLECTIFMGSLDMERQPSNVLLMKHLGAKVKS
VNGNFKDAVSEGIRNWVNDLETSYFLAGAAVGPHPCPTMVREFQSIIGKETRKQAMEKWGGKPDVLVAC

VGSGSNALGLFHEFILDQDVRLIGVEAGGTGLDSGKHSATMARGEVGVYHGAMSYLLQDEEGQIIGPHSIG
VGLEYPGVSPELSYLKDIGRAEFSSVTDEQALEAYKRLCRLEGIFPALESSHALAFLGKLCSTLKDGEKVVV
NLSGRGDKDAAAVFNHTPKHE
>SOLANUM MELONGENA_SMEL4.1_04G005180.1.01
MAGNVKGILGQGIFAKSSTKVQGFPSHTYKANVISCVAIGPSQVPQTPWKLIFHEKERPLLSNEKFGIYGGK
FVPETLISSLTKLDYEFNSALRDPLFQMELEVALRDYVGRETPLYFAQRLTDYYKKMNRGIGPDIYLKREDL
NHGGAHKINNAIAQAMLAKRMGCKNVVAPTGAGQHGVATAAACAKLSLECTIFMGSLDMERQPSNLLL
MKHLGAKVKSVEGSFKDAVSEGIRNWVSDLDANYFLAGGAIGPHPCPTMVREFQSIIGKETRKQAMDKW
GGKPHVLVACVGSGSNALGLFHEFIQDEDVRLIGVEAGGTGLDSGKHSATIARGEVGVYHGAMSYLLQDE
EGQIIGPHSIGVGLEYPGVSPELSFLKDIGRAEFASVTDEEALEAYKRLCRLEGIFPALESCHALAFLDKLCST
LKDGEKVIVNLSGRGDKDAAAVFNHTANMDEKKIF
>PETUNIA AXILLARIS_PEAXI162SCF00950G00611.1
MACNVEQVIFRRGKFATCGEARLSSNRKWQGKFVVSSVAVGPSRGTLTTADQEQQSPWKLVYHGKKGPS
SNEKFGIFGGKFVPETLISSLTKLNYEFNSALHDPQFQMELGVALRDYVGRETPLYFAKRLTEHYKSSNSGK
GPDIYLKREDLNHGGAHKINNAIAQAMLAKRMDCKSIVAATGAGQHGVATAASCAKLSLACTIFMGSLD
MERQPSNVLLMQHLGAKVQKIYWVTNLDTSYFLSGAAIGPHPCPTMVREFQSVIGKETRKQAMEKWGGK
PDVLVACVGSGSNALGLFHEFIEDEDVRLIGVEAGGNGLDSGKHSATIARGQVGVYHGAMSYLLQDEEGQ
INVPHSIGVGLEYPGVSPELSFLNDIGRAEFSSVTDEEALEAYKRLCRLEGILPALEASHALAFLDKLCPTLN
DGQKVVVNCSGRGEKDAARVFCHTPTHE
>SOLANUM PENNELLII_SOPEN10G001310.1
MDLQVALKDYVGRETPLYFAQRLTDYYKSLNKGIGPDIYLKREYLNHGGAHKINNAIAQAMLAKRMGCK
NVVASTGAGQHGVGTAAACAKLSLECTIFMGSLDMERQPSNVLLMNHLGAKVKCVEGSFKDAMSEGIRN
WVNDLETSYFLAGAAIGPHPCPTMVREFQSIIGKETRKQAMDKWGGKPHVLVACVGSGSNALGLFHEFIQ
DVDVRLIGVEAGGIGLDSGKHSATMARGEVGVYHGAMSYLLQDEEGQLIEPHSIGVGLEYPGVSPELSYLK
DIGRAEFSTVTDEEAIKAYKRLCILEGIFPALESCHALAFLDKLCSTLKDGEKVIVNLSGRGDKDAEAVFNH
TPKHK
